# The representational geometry of cognitive maps under cognitive control

**DOI:** 10.1101/2023.02.04.527142

**Authors:** Seongmin A. Park, Maryam Zolfaghar, Jacob Russin, Douglas S. Miller, Randall C. O’Reilly, Erie D. Boorman

## Abstract

Recent work has shown that the brain abstracts non-spatial relationships between entities or task states into representations called cognitive maps. Here, we investigate how cognitive control enables flexible top-down selection of goal-relevant information from multidimensional cognitive maps retrieved from memory. We examine the relationship between cognitive control and representational geometry by conducting parallel analyses of functional magnetic resonance imaging data and recurrent neural network models trained to perform the same task. We find both stable map-like representations in a medial temporal lobe and orbitofrontal cortical network that reflect both task-relevant and irrelevant dimensions and context-dependent, orthogonal representations of only task-relevant dimensions in a frontoparietal network. These representational motifs also emerge with distinct temporal profiles over the course of training in the recurrent neural network, with map-like representations appearing first. We further show that increasing control demands due to incongruence (conflicting responses) between current task-relevant and irrelevant dimensions impact the geometry of subjective representations, and the degree of this effect further accounts for individual differences in cognitive control. Taken together, our findings show how complementary representational geometries balance stability and behavioral flexibility, and reveal an intricate relationship between cognitive control and cognitive map geometry.

## Introduction

Generalizing previous experiences to make flexible decisions according to current goals is a hallmark of adaptive behavior. This behavioral flexibility depends on the development of representations that can be reused across multiple tasks, as well as the capacity for flexible top-down selection of information according to changing task goals ^1–10^. Recent findings suggest that the hippocampus (HC), entorhinal cortex (EC), and orbitofrontal cortex (OFC) organize relationships between items, individuals, events, or abstract task states into a unitary combined representation, even when multiple task dimensions are sampled in different task contexts ^11–18^. Recent computational models of cognitive control suggest that the brain trades off the benefits of low-dimensional representations for fast learning and generalization across multiple tasks against the benefits of high-dimensional representations for flexible decision making ^4,5,19–24^. An intriguing but unexplored possibility is that an analogous tradeoff may determine the geometry of cognitive maps.

Previous studies of cognitive control have revealed the mechanisms by which the brain biases attention toward explicitly presented sensory features, such as color, motion, shape, or orthographic features, while participants integrate sensory inputs providing evidence favoring different, sometimes conflicting, responses ^8,9,25–27^. While these studies have made substantial progress towards understanding the top-down biasing of sensory information based on current task sets, relatively little is known about how cognitive control may apply to dimensions from mnemonic representations, particularly cognitive maps. When information is retrieved from memory, the brain may favor high-dimensional population coding enabling any specific dimensions to be flexibly selected based on the current goal. Moreover, the representational geometry of the cognitive map defined by the neuronal population may play a central role in shaping control demands when they are not determined by incoming sensory feature evidence, but instead by learned relationships retrieved from memory, as is commonplace in the real world.

To investigate the interaction between changing task goals and the representational geometry of cognitive maps in guiding flexible decision making, we examined how neural representations of two-dimensional (2-D) social hierarchies are influenced by task demands and their potential impact on cognitive control (Fig.1a). When making relational inferences between entities – e.g., who is more senior, or, on the other hand, more popular, amongst colleagues – control demands increase due to interference caused by incongruence between currently task-relevant and irrelevant dimensions. For example, while José is more senior than Gabriela and has more influence at work, Gabriela is more popular and, hence, has more influence at a social gathering.

**Figure 1.**
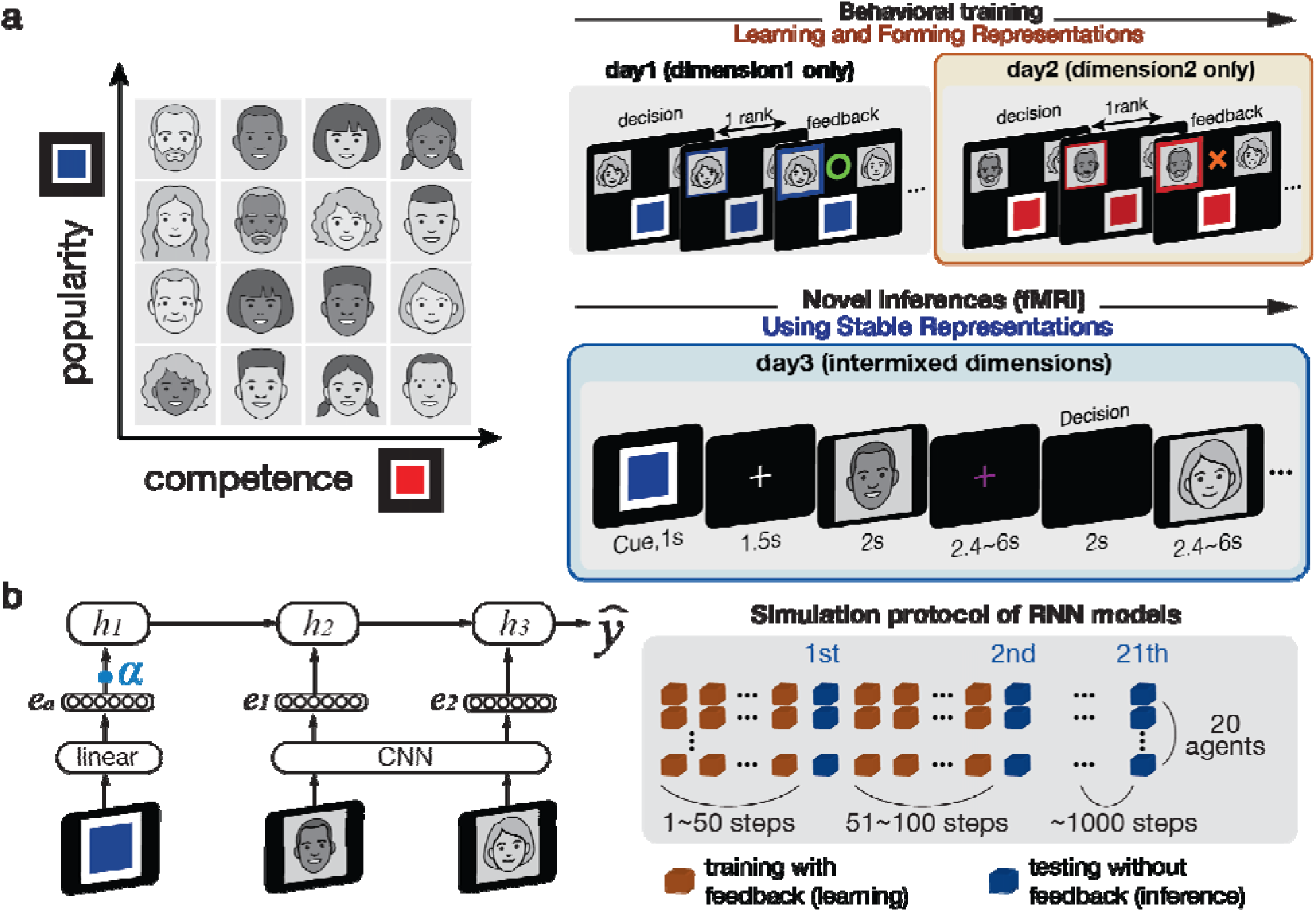
Experimental task. **a**. Participants learned the relationships between 16 faces through feedback in two independent social hierarchy dimensions (competence and popularity). The task used photographs of human faces; the stimuli shown here have been replaced with author-created illustrations. To represent the relational task structure, the brain could construct a representation that integrates relationships across both dimensions, enabling a task-independent (context-invariant) representation shared across the two task contexts (left). During behavioral training (right top), participants learned neighboring relationships (i.e., pairs separated by one rank) in each social hierarchy dimension on separate days. In the fMRI task (right bottom), faces were presented sequentially, and participants made novel inferences about their relationship, selecting the higher-ranked face in the cued social hierarchy dimension upon presentation of the second face. The task-relevant dimension was signaled by a colored context cue at the beginning of each trial, with cue colors counterbalanced across participants and dimensions randomly intermixed across trials. Because no feedback was provided during inference, the representations constructed during learning could be maintained without updating. **b**. The architecture of the recurrent neural network (RNN) trained to perform the same task as human participants. The model used a convolutional neural network (CNN) to process images and long short-term memory (LSTM) to process the sequence of inputs over time. *e*: embeddings; *h*: hidden layers; : outcome; : the level of degradation associated with additional ablation simulation. During training, the RNN was allowed to update the activity patterns associated with each face under different task contexts based on feedback. This update process was paused during the test phase, corresponding to the inference phase for human participants, during which the RNN was not allowed to modify its representational structure based on feedback. The representational geometry was evaluated repeatedly throughout the test phase, every 50 training steps over 1,000 training steps. The ablation analysis examined how degrading context information influenced the formation of representational geometry. Specifically, the context input was degraded across all trials during training but remained intact during test steps.

Recent findings suggest that the representational geometry of task representations might be determined by task demands such that they facilitate generalization to similar contexts that share a decision policy (i.e., choice rule) ^5,23,28–32^. When the same policy can be applied to different task contexts, the brain may construct a shared representation along a single dimension that supports rapid learning and generalization between contexts (Fig.2a, left panel). In contrast, when each task context requires an independent policy, the brain should theoretically form separate representations for each context—or organize them along orthogonal dimensions—such that representations of one task context do not interfere with those of another (Fig.2a, right panel). In everyday learning, however, we often encounter uncertainty as to whether the underlying task structure can be best captured along a shared dimension or instead orthogonal dimensions. This ambiguity poses a fundamental challenge for the brain in determining how to organize task representations to balance useful generalization and context-specificity that mitigates interference.

**Figure 2.**
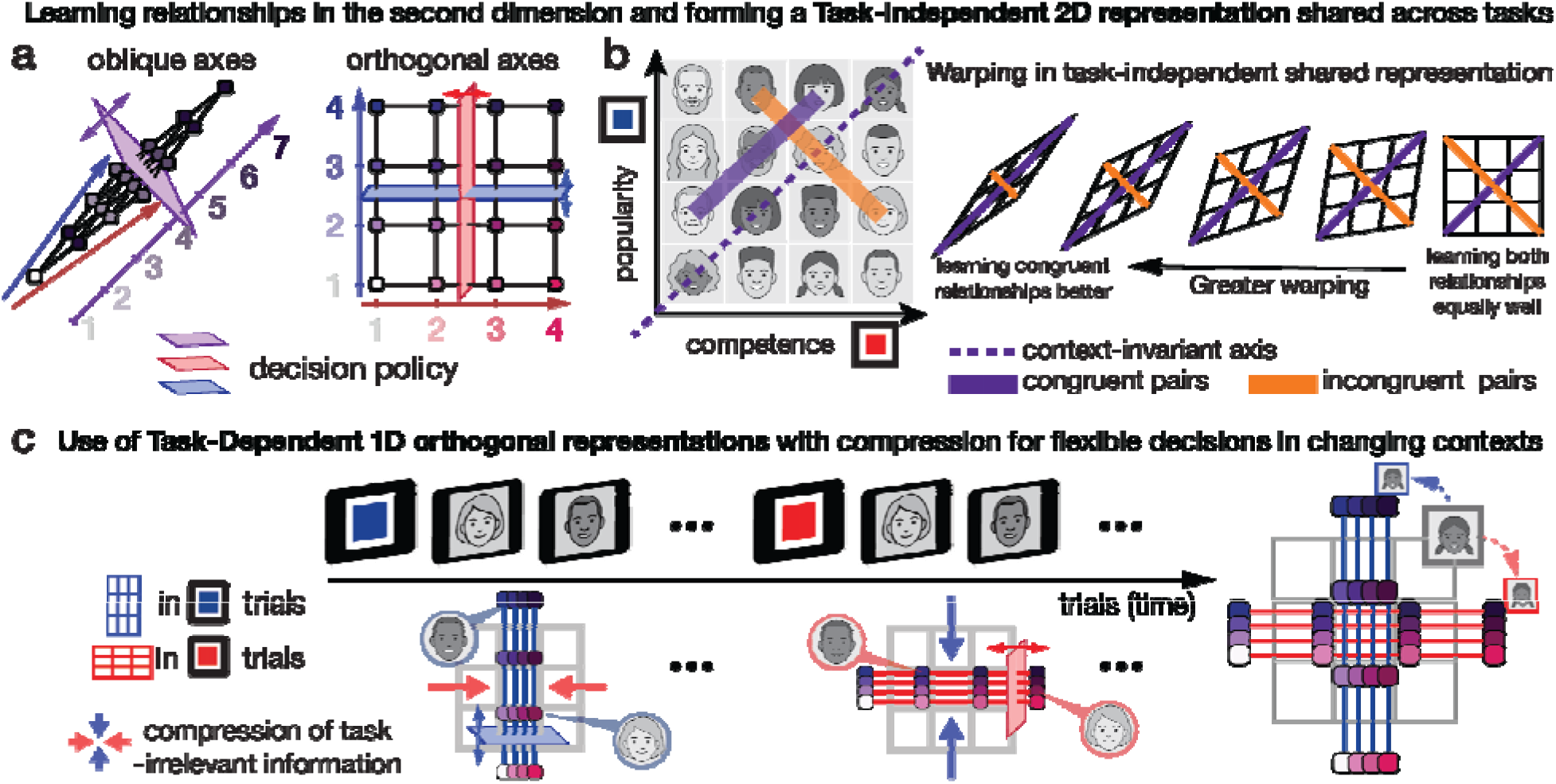
The interaction between cognitive control and representations of relational structure. **a**. During behavioral training, different levels of cognitive control in learning and updating congruent and incongruent relationships could shape the geometry of context-invariant representations in the brain. The brain could construct a representation with oblique axes (left) by updating face ranks based on overall win–loss ratios across comparisons, thereby integrating information across dimensions. Alternatively, it could construct a representation with orthogonal axes (right) by maintaining separate ranks for each task-relevant dimension, reducing interference from task-irrelevant information on incongruent trials. Both representations capture relationships across dimensions and support generalization across task contexts. However, the oblique representation allows the same decision policy (purple plane) to be applied across dimensions, whereas the orthogonal representation supports distinct decision policies (red and blue planes). Thus, representational geometry may influence accuracy in inferring incongruent relationships and behavioral flexibility when task contexts change. **b**. Individual differences in task-independent two-dimensional (2-D) representations. During behavioral training, participants may differ in their ability to maintain the previously learned rank relationships along the first dimension while updating relationships in the second, which was learned on a different day. Because learning incongruent relationships typically requires greater cognitive control than learning congruent ones, weaker control may lead to less accurate representations of incongruent relationships. These differences may alter the relative distances between faces associated with congruent (purple) and incongruent (orange) relationships, resulting in individual differences in representational geometry and varying degrees of warping along the context-invariant axis (purple dashed line). **c**. Task-dependent one-dimensional (1-D) orthogonal representations. To make correct inferences, the brain must flexibly extract and compare ranks along the task-relevant dimension according to the continuously changing task context. To facilitate effective value coding for the current goal, the brain may expand the multidimensional map-like representation along the task-relevant axis and/or compress task-irrelevant information according to the current goal. As a result, we hypothesize that the brain may employ a context-dependent 1-D representation along the orthogonal axes, alongside the context-independent 2-D representation, simultaneously when making inferences about relationships based on the representation retrieved from memory.

Cognitive control may play an important role not only in overcoming interference between competing responses, but also in the construction of such task representations ^*3,33*^. During the construction of a multidimensional representation, congruence or incongruence stems from whether the newly learned relationships align with or conflict with those established in the previously learned representations. If the control demands associated with learning incongruent versus congruent relationships shape the geometry of subjective representations, we would expect the brain to adopt a versatile representational strategy (Fig.2b). That is, congruence would favor shared representations that support generalization across dimensions, whereas incongruence would favor segregated or orthogonalized representations that minimize interference and enable flexible switching across contexts with divergent response requirements.

To facilitate effective value computations at the time of decision making, control- and decision-related brain regions may collapse or compress the multidimensional map-like representations into one-dimensional (1-D) representations according to the current task goal (Fig.2c). When features of sensory stimuli guiding responses are presented explicitly, previous studies have shown top-down control from the prefrontal cortex (PFC) gating the inputs of task-relevant information on the early sensory cortex ^7,27,34^ or attenuating task-irrelevant information relative to task-relevant information, even in proficient participants who underwent intensive training ^8,9^. Other studies have instead proposed that currently task-irrelevant evidence is simultaneously and orthogonally encoded faithfully in prefrontal cortex without any top-down gating ^25,26,35^. Whether or how these competing mechanisms apply to inferences over internal representations of cognitive maps, where the task-relevant dimensions are not determined by features of sensory inputs, but by the learned relationships between entities retrieved from memory, is unclear.

To address these questions, we combined recurrent neural network (RNN) models with human fMRI data on a task that required subjects to make context-dependent inferences about relationships between entities (Fig.1b). We found 2-D representations that were stable across behaviorally-relevant contexts in the hippocampus (HC), entorhinal cortex (EC), and orbitofrontal cortex (OFC), even though only 1-D relationships were behaviorally-relevant for current decisions. In parallel, we found context-dependent, orthogonal1-D representations with compression of task-irrelevant dimensions relative to task-relevant dimensions in the dorsomedial frontal cortex (dmFC), lateral prefrontal cortex (lPFC), posterior cingulate cortex (PCC), and inferior parietal lobule (IPL). Consistent with human fMRI, we found both context-invariant 2-D representations and context-dependent 1-D representations in the RNN trained on the same task. Moreover, we found evidence of warping in both HC representations and the RNN such that representational geometries were skewed along the congruent versus incongruent axis. Notably, our results show parallels between representations of cognitive maps in human brains and those that emerge in the RNN, and thus provide insight into the neural mechanisms that support the construction and geometry of cognitive maps, the flexible prioritization of goal-relevant information, and their interplay.

## Results

To examine how representational geometry and cognitive control may interact to form subjective cognitive maps, we performed additional behavioral and neural analyses on previously collected fMRI data ^11^ and compared neural representations in humans to those in hidden layers of RNN models trained on the same task (both not published previously). Both human participants (*n*=27) and RNN models (classic long short-term memory [LSTM] networks) ^36^ learned the relative ranks of 16 individual faces on two social hierarchy dimensions (Fig.1a)—competence and popularity — using a similar previously detailed training protocol ^37^. On each trial, participants selected which of two sequentially presented faces ranked higher along one of the two dimensions, which was indicated by a colored context cue. During training both human participants and the RNN learned only about relationships between neighboring entities – i.e., individuals whose rank differed only by one level along the given dimension -through feedback on their decisions. Human participants learned the relationships in each of the two dimensions on different days, but the model learned them in a single block with interleaved trials (see Methods) to avoid catastrophic forgetting ^38,39^. After training, both human participants (Fig.1a) and the RNN (Fig.1b and Eq.16-18) were asked to infer relationships that were not seen during training. During this test, contexts were randomly interleaved for the human participants as well as the model, thus requiring subjects to flexibly switch between contextually defined task dimensions. It is important to note that neither participants nor the model were provided with any information about the true 4×4 grid structure of the social hierarchy, nor were they asked to construct it spatially.

Using both human fMRI data and hidden states of the RNN, first, we examined unitary map-like representations of the social hierarchies in both dimensions (context-invariant representations) and tested for the effects of cognitive control modulating representational geometries to facilitate execution of the current task context while resolving interference from task-irrelevant information (context-dependent representations). Second, we tested for hypothesized warping in the subjective 2-D map-like representations. Third, we explored the relationship between cognitive control and different levels of warping in the representations across individual participants. Finally, we performed ablation experiments on the RNN that informed additional fMRI analyses that together shed light on the mechanisms that produced the observed warping.

### 1. Context-invariant 2-D and orthogonal 1-D context-dependent representations

In our previous work ^11^, we provided evidence suggesting that the brain builds a unitary, context-invariant representation across different social hierarchy dimensions and uses it to make inferences, even when only one dimension is relevant for current decisions. Specifically, we found that the Euclidean distances between individuals in the true 2-D social hierarchy structure explained the decision-related univariate BOLD activity in the EC and ventromedial prefrontal cortex (vmPFC), as well as reaction times (RTs), better than 1-D task-relevant rank differences alone. Moreover, multivariate analyses of BOLD patterns elicited by face presentations identified 2-D map-like representations of the social hierarchy in the HC, EC, and OFC, such that closer individuals in the true hierarchy (measured by pairwise Euclidian distance) were represented increasingly and linearly more similarly (measured by pairwise pattern similarity).

While sharing representations across multiple task contexts may benefit generalization and allow faster learning across contexts, the brain may also benefit from flexibly transforming this higher-dimensional representation into 1-D representations according to the current task goals to effectively resolve interference between information in the task-relevant and task-irrelevant dimensions. Motivated by recent evidence in the perceptual selective-attention literature ^8,9,25,26,35^, we tested for evidence that the two dimensions were represented orthogonally in any additional brain regions, with information along the currently irrelevant dimension being compressed. To test these hypotheses while controlling for alternative representational geometries in the brain, we examine the extent to which each of several candidate model representational dissimilarity matrices (RDMs) explains the pattern dissimilarity associated with individuals in the social hierarchy using a general linear model (GLM) (Fig.3a). Specifically, in the same GLM, we inputted the pairwise Euclidean distances in the 2-D social hierarchy and those computed from orthogonal 1-D representations with compression according to the task context (1D orthogonal), as well as other control RDMs (1-D task relevant rank representation (1D task-relevant), task context, and learning group; see Methods), to pit them against one another to explain variance in pattern dissimilarity (Eq.1). Note that while both 1D orthogonal and 1D task-relevant models sought to identify brain areas whose activity patterns represent rank in the current task-relevant dimension, the 1-D orthogonal representation (1*D* in Eq.1) differs from the 1-D task-relevant representation (1*R* in Eq.1) in an important way. While the activity patterns associated with rank in the 1-D task-relevant rank representation can be generalized across the two social hierarchy dimensions, those in the 1-D orthogonal representation are maximally decorrelated, thereby favoring segregation and minimizing interference, and thus cannot be generalized across dimensions (Fig. 3a and Supplementary Fig.S1 in supporting information (SI)).

**Figure 3.**
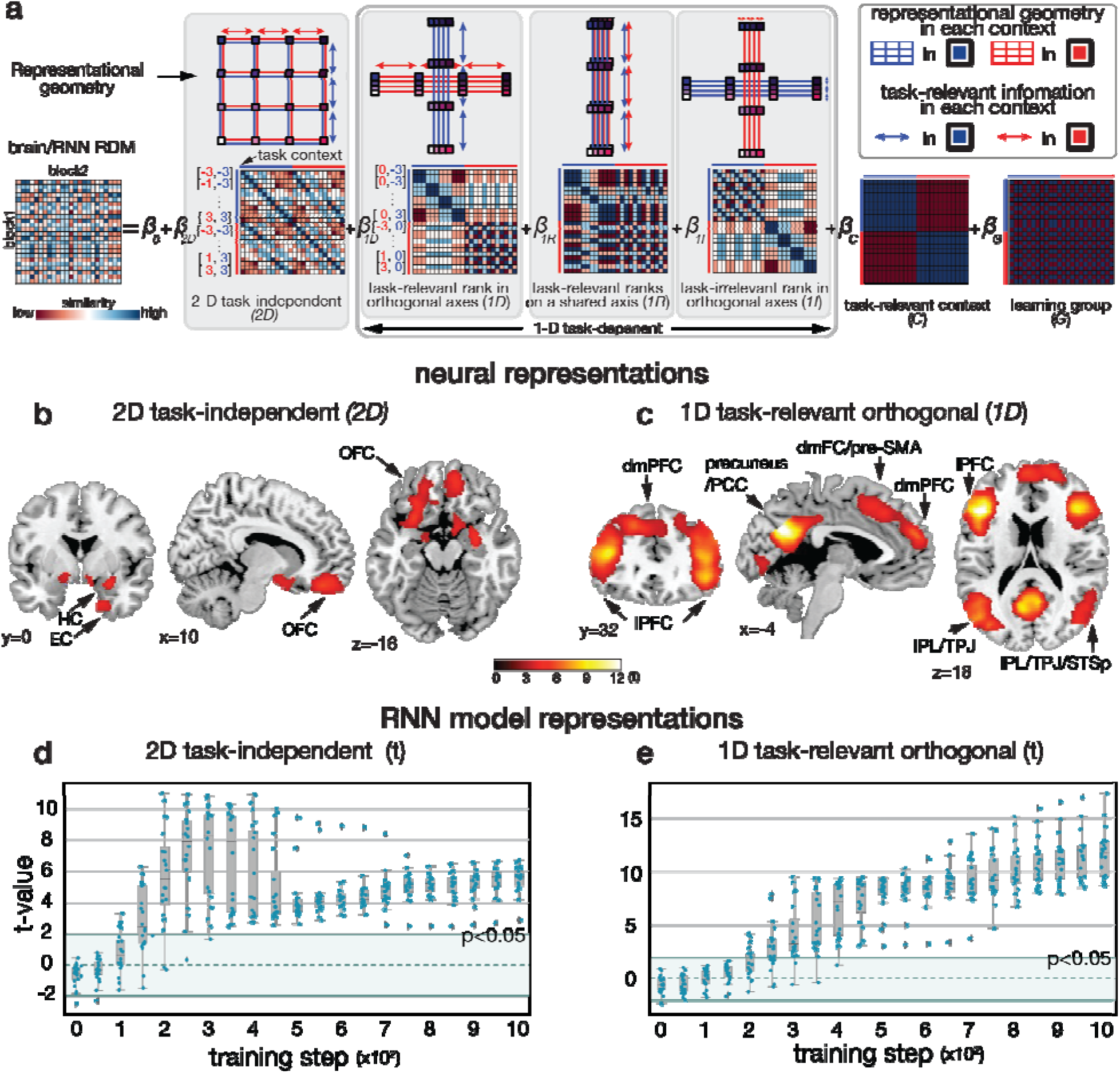
Simultaneous 2-D context-invariant and 1-D orthogonal context-dependent representations in the brain and the RNN. **a**. Simultaneous representations of different geometrical structures were found in both the brain and the RNN. Different representational geometry models were fit against each other within a single general linear model (GLM) and tested with representational similarity analysis (RSA) by estimating pairwise distances between individuals from their activity patterns’ dissimilarity across the entire brain or in the hidden layers of the RNN. Illustrated from left to right is the general linear model (GLM) we tested. In order from left to right, we tested for the representational geometry of the 2-D context-invariant representation across contexts (*2D*), the task-relevant information in the 1-D context-dependent orthogonal representation (*1D*), the 1-D task-dependent rank representation that can be generalized across contexts (*1R*), the task-irrelevant information in the 1-D context-dependent orthogonal representation (*1I*), task context (*C*), and learning group (*G*). In the RDM templates, the colored-line indicates the relevant context dimension (popularity or competence), and the geometric form shows the shapes of the idealized representations capturing our hypotheses and how distances were computed. **b**. RSA results of the 2-D context-invariant representation RDM are shown in the hippocampus (HC), entorhinal cortex (EC), and orbitofrontal cortex (OFC). **c**. RSA results of the task-relevant 1-D orthogonal representation. Significant effects are shown in the lateral prefrontal cortex (lPFC), dorsomedial frontal cortex (dmFC) and prefrontal cortex (dmPFC), posterior cingulate cortex (PCC), and inferior parietal lobule (IPL). **b**. and **c**. All the t-statistic brain maps are displayed at a cluster-corrected threshold p_TFCE_<0.05 over the whole brain for all brain regions. **d**. The same analysis of the RNN shows changes in the level of 2-D representation in the hidden layers over the course of training. **e**. Timecourse of emergent 1-D orthogonal representation according to the current task demand during RNN training. **d. and e**. Box plots show the levels of representations (t values) over 20 runs (a dot indicates each run). Box, lower and upper quartiles; line, median; whiskers, range of the data excluding outliers; ⍰ sign, the whiskers’ range of outliers.

Consistent with our previous study ^11^, this analysis found the 2-D map-like representations in the HC (peak=[28,-2,-18], *t*=4.67), EC ([20,0,-36], *t*=4.02) and vmPFC encompassing the medial OFC ([14,42,-22], *t*=5.83), and lateral OFC ([28, 22, -22], *t*= 3.96 in right and [-24, 26, - 16], *t*= 4.55 in left) (pTFCE<0.05, df=26 for all group-level fMRI analyses; Fig.3b). In parallel, we found 1-D orthogonal representations of the relevant dimensions with the irrelevant dimension compressed in the dorsomedial PFC (dmPFC, [-4,54,24], *t*=5.99), the dorsomedial frontal cortex (dmFC) encompassing the pre-supplementary motor area (pre-SMA) ([4,14,48], *t*=5.59), lateral PFC (lPFC, [42,26,18], *t*=9.03 in right; [-42,24,20], *t*=13.48 in left), precuneus/posterior cingulate cortex (PCC, [-4,-64,28], *t*=13.60), right inferior parietal lobule (IPL, [46,-64,22], *t*=5.20) extending into temporoparietal junction area (TPJ), and left IPL extending into TPJ and posterior superior temporal sulcus (STSp) ([-48,-72,18], *t*=6.72) and fusiform gyrus ([30,-84,-10], *t*=5.54 in right; [-40,-60,-20], *t*=6.44 in left) (pTFCE<0.05; Fig.3c). See Supplementary Table 1 in supplementary information for a full list of brain areas surviving correction.

Importantly, follow-up control analyses confirmed that these effects were specific to the 1-D orthogonal representation, and could not be explained by the 1-D task-relevant representation alone. First, these effects remained even when considering only cross-context pattern dissimilarity, where the RDMs make divergent predictions, and thus were not solely due to within-context pattern dissimilarity, where distinguishing between the 1*D* and 1*R* models becomes challenging (Supplementary Fig.S1). Moreover, within these brain areas showing significant effects of the 1-D orthogonal representation, we did not find evidence of weaker but still significant effects of the task irrelevant information (1*I*), even at a lenient threshold (*p*<0.05 uncorrected) (Supplementary Fig.S1e).

Taken together, these representational similarity analyses (RSA) show that the HC, EC and OFC exhibit stable, task-invariant 2-D map-like representations, in parallel to task-dependent 1-D orthogonal representations with compression along the irrelevant dimension, in a frontoparietal network ideal for computing context-dependent decision values. Indeed, the extent to which neural activity patterns reflected 2-D and 1-D representations was neither positively nor inversely correlated with each other across participants (Supplementary Table 2), supporting the interpretation of simultaneous representations of both types of representational geometry.

### 2. 2-D and orthogonal 1-D representations emerge in the RNN

To test if, under what conditions, and when during training these representations might emerge as an efficient solution to our task, we performed the same GLM-based RSA analyses of our RNN model (Fig.3a). We also found the emergence of both representations in the neural network model, even though we did not design the models with any components specialized to produce these representations. Just as in the fMRI data, the pairwise distances between representations in the neural network model were explained by the 2-D Euclidean distances between individuals in the context-invariant 2-D social hierarchy structure (Fig.3d), as well as those in the context-dependent 1-D orthogonal representations with compression (Fig.3e). The 2-D map-like representations emerged at a relatively early phase of training (the representation was significant [*p*<0.05] at 150 steps of training [the median run: *t*=4.35, *p*<0.001; df=19 for all RNN representation analyses in Fig.3]; the peak was at training step 300 [the median run: *t*=8.31, *p*<0.001]) and were maintained throughout training (*t*=5.63, *p*<0.001 at the last training step). While the magnitude of the effect decreased by the end of training, this reduction was not statistically significant (*t*=-1.24, *p*=0.22). Simultaneously, the model also utilized task-dependent orthogonal representations, and this tendency for compression increased continuously over training (significant at training step 250 [the median run: *t*=4.53, *p*<0.001]; peak at training step 1000 [the median run: t =11.75, *p*<0.001]). Thus, both shared 2-D and orthogonal 1-D representations emerged as complementary, efficient representations to solve our relational inference task, but with different time courses.

### 3. Effects of cognitive control on representational geometry

Based on empirical and theoretical evidence that task demands during learning might shape a task space’s subjective representation ^5,23,28–32^, we considered the extent to which cognitive control demands might influence the relational configuration between entities and hence the representational geometry of the 2-D cognitive map. Specifically, we test the hypothesis that increasing control demands for incongruent compared to congruent pairs produces warping along the congruent axis in the subjective representation of the 2-D social hierarchy. We tested this hypothesis in four independent ways, including one behavioral analysis and three neural-level analyses.

#### 3.a . Behavioral evidence of differential cognitive control demands on incongruent versus congruent trials

Through the utilization of four multiple linear regression models (Fig.4a; Eq.2-5), we evaluated the potential influence of warping in the 2D representation by leveraging the variability of RTs, if such warping were to exist. Within a warped representation, the rank differences between faces would undergo alterations, not only in the task-relevant dimension but also in the task-irrelevant dimension, with both being contingent upon the congruence of relationships. Consequently, there would be an increase in pairwise 2-D Euclidean distances between congruent pairs and a decrease between incongruent pairs, reflecting the different demands for cognitive control.

**Figure 4.**
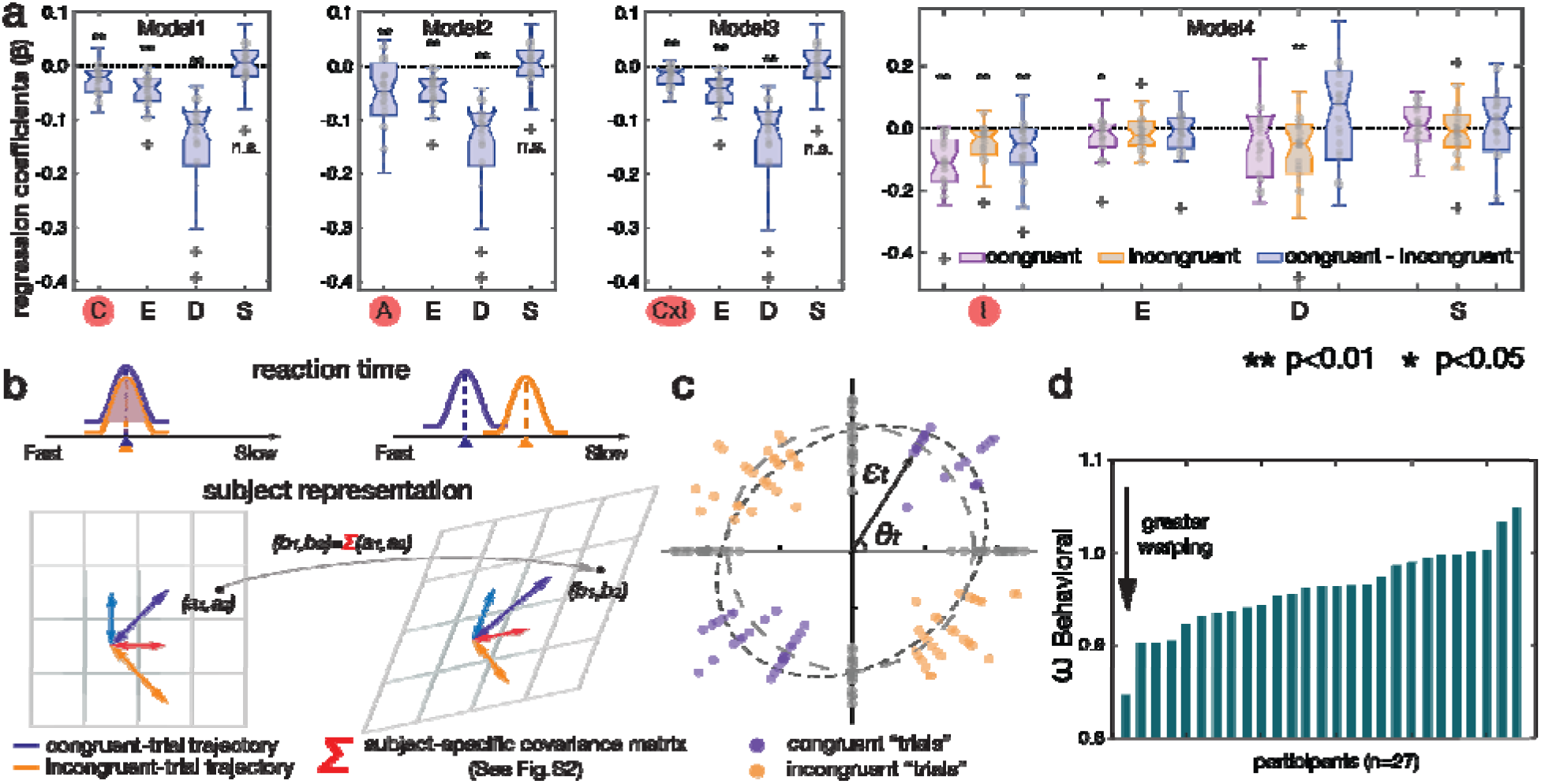
Behavioral results suggesting potential warping of representations. **a**. Four multiple linear regressions tested the predicted effects of inference rank differences, Euclidean distances between faces, and congruence on reaction times (RTs). Model 1 showed that context congruency (C), 1-D rank difference in the task-relevant dimension (D), and 2-D Euclidean distance (E), but not context switching (S), significantly explained RT variance. Negative effects indicate faster RTs with increasing rank differences and on congruent versus incongruent trials. Model 2 yielded similar findings, with context congruency represented as angles (A) of inferred trajectories relative to the context-invariant axis. Models 3 and 4 further showed that task-irrelevant information (I) was modulated by context congruency (C). Regression coefficients were estimated separately for each participant using all correct trials (up to 208 per participant), with group-level significance assessed using two-sided one-sample t-tests (n = 27). Box plots show coefficient distributions; boxes indicate quartiles, horizontal lines medians, whiskers the most extreme non-outlier values, and + symbols outliers. **b**. Schematic rationale for approximating the subjective representation from RTs. RTs are inversely correlated with Euclidean distances of inferred trajectories (Fig. 3a). If the representation is not warped (left), RTs for congruent (purple) and equidistant incongruent (orange) relationships should not differ. In a warped representation (right), congruent distances are elongated and incongruent distances shortened. Thus, warping along the context-invariant axis should produce faster RTs for congruent than incongruent inferences across decision trajectories. **c**. Example warping estimate from one participant. To test whether RTs varied with trajectory angles (θ) while controlling for Euclidean distance, we computed a covariance matrix (Σ) from the logarithmic distribution of −RTs, normalized by trajectory distances (ε) across trials (Eqs. 7–12). An unbiased circular distribution of ε indicates no warping, whereas elongation along the context-invariant axis indicates warping. We quantified warping (ω^***Behavioral***^) as the ratio of incongruent to congruent distances. This covariance-based measure captures systematic angle-dependent biases while reducing sensitivity to individual-trial fluctuations and components unrelated to decision angles. **d**. Individual differences in behavioral warping (ω^***Behavioral***^). Smaller values indicate greater warping; values below 1 indicate warping along the context-invariant axis.

Analysis of log-transformed RTs revealed a significant effect of the congruence of relationships between individuals across the two task dimensions, even after accounting for other distance measures of inferred trajectories, including the 1-D rank difference in the task-relevant dimension (Fig.4a and Supplementary Table 3). This effect was robust, whether we defined congruence as a categorical value (Model 1:-0.027±0.005(SE), *t*=-5.164, *p*<0.001; df=26 throughout all RT regression models) or as a continuous value derived from the angles of inferred trajectory for inferences (Model 2:-0.053±0.013, *t*=-4.14, *p*<0.001), indicating that people were significantly slower for incongruent than equidistant congruent inferences. Furthermore, to explore the effects of rank differences in task-relevant and irrelevant dimensions separately, we tested two additional models. In these models, we decomposed the 2-D Euclidean distances into two constituent 1-D rank differences, one for the task-relevant dimension and the other for the task-irrelevant dimension. We discovered a significant interaction effect between task-irrelevant information and congruence on RTs (Model 3:-0.02±0.004, *t*=-5.34, *p*<0.001). To unpack this interaction, we confirmed that the impact of rank difference in the task-irrelevant dimension was more pronounced, resulting in faster RTs, during congruent trials (-0.118±0.019) compared to incongruent trials (-0.047±0.013). This difference was statistically significant (Model 4:-0.070±0.021, *t*=-3.45, *p*=0.002) (Fig.4a).

Last, model comparison using BIC revealed that integrating the interaction effect between context congruency and task-irrelevant rank difference significantly improved model fit. Model 3, which included this interaction term, outperformed Models 1 and 2, which considered only context congruency, in explaining the variance in reaction times (compared to Model 1: *t*=2.546, *p*=0.017; compared to Model 2: *t*=3.178, *p*=0.004). These analyses indicate that participants exhibited slower RTs on incongruent trials compared to congruent ones, even when accounting for distances in both the 1-D task-relevant dimension and the 2-D Euclidian distance incorporating both dimensions. See SI Table 3 for the results of these four models predicting accuracy, which showed limited variability due to high overall accuracy levels, using logistic regressions with the same regressors.

Subsequently, by computing effects of the degree of congruence (defined as the vector angles of inferred trajectories, *θ*) on the relationship between Euclidian distance and RTs (*ε*), we approximated the hypothesized subjective representational geometry and the implied level of warping per participant (Fig.4b and Eq.7-8). The levels of potential warping (*ω*_*Behavioral*_) were estimated as the ratios of the incongruent diagonal distances to the congruent diagonal distances in the 4×4 structure of the social hierarchy after applying the covariance estimated from the distribution of [ε, θ] of each participant (Fig.4c). The *ω*_Behavioral_ was quantified using the subject-specific covariance matrix (Σ; Eq.9-10). The Σ captures the level of distortion in the influence of Euclidean distances on RTs by considering the angles of inference trajectories (Eq.11-12). The mean ratio between diagonals (*ω*_*Behavioral*_) was 0.961±0.008 (SE). We confirm that this effect is significant compared to a shuffled baseline (*t*=2.93, *p*=0.007, 1000 permutations; Supplementary Fig.S2), suggesting consistent effects of context congruence along the direction of the context-invariant axis. We further validated that this procedure could estimate the level of warping in subjective representations if warping exists, through confirmatory simulations with synthetic data (Supplementary Fig.S2; Eq.13). The distribution of the estimated level of warping across participants is shown in Fig.4d. We used *ω*_*Behavioral*_, the diagonal ratio (relative values) to index individual differences in cognitive control in further neural analyses.

Note that subjective warping estimates should be robust to the specific model used to estimate trial-wise RT residuals. Because decision values were symmetrically distributed across directions in the current study, applying the same linear or nonlinear mapping from decision value to RT could change the overall scale of RT predictions, but should not selectively distort the estimated warping. Consistent with this assumption, residuals from the linear model were highly correlated with residuals from an alternative DDM-based model (Eq.14; mean r±SE=0.97±0.0035, 95% CI=0.96–0.98), and the resulting rank order of warping estimates across participants remained consistent across models (Spearman’s ρ=0.92, *p*<0.001).

#### 3.b. Neural evidence of warped representations

Next, we tested if the inferred trajectories over the hypothesized warped representations better explain the neural encoding of decision values for inferences in the EC and vmPFC during decisions. We defined *a priori* regions of interest (ROIs) anatomically based on our previous findings^11^ and found that the Euclidean distances in the *warped* representations inferred from RTs could explain additional variance in activity in the bilateral EC (peak [x,y,z]=[28,-16,-32], *t*=4.06, pTFCE<0.01; [-22,-14,-32], *t*=4.84, pTFCE<0.01 in the EC ROI) and the vmPFC ([2,24,-12], *t*=4.99, pTFCE<0.01 in the vmPFC ROI), after controlling for the variance accounted for by true Euclidean distances between individuals in the 2-D social hierarchy and, importantly, RTs across trials (Fig.5a). This finding indicates that the hypothesized warped representations explain additional variance in the effect of Euclidian distance on decision-related univariate activity in these brain regions. It is important to note that the behavioral warping measure was derived from the entire covariance matrix (Σ), ensuring that this effect is not determined by the RT indices of difficulty on the current decision triaI. Additionally, we tested an alternative, but non-mutually exclusive hypothesis that changes in cognitive control might stem from the costs of task switching, rather than warped neural representations. Notably, we did not find statistically significant effects of switching task contexts on RTs (*S* in Fig.4a), or on neural activity across the whole brain even at a lenient threshold (*p*<0.005, uncorrected). Moreover, incongruent trials following congruent trials did not show statistically significantly slower RTs during inference or during learning of relationships in the second dimension (one-sided t-tests, *p*=0.990 and *p*=0.843, respectively), providing no statistically significant evidence of an additional cost associated with the prior-trial control state ^40^.

**Figure 5.**
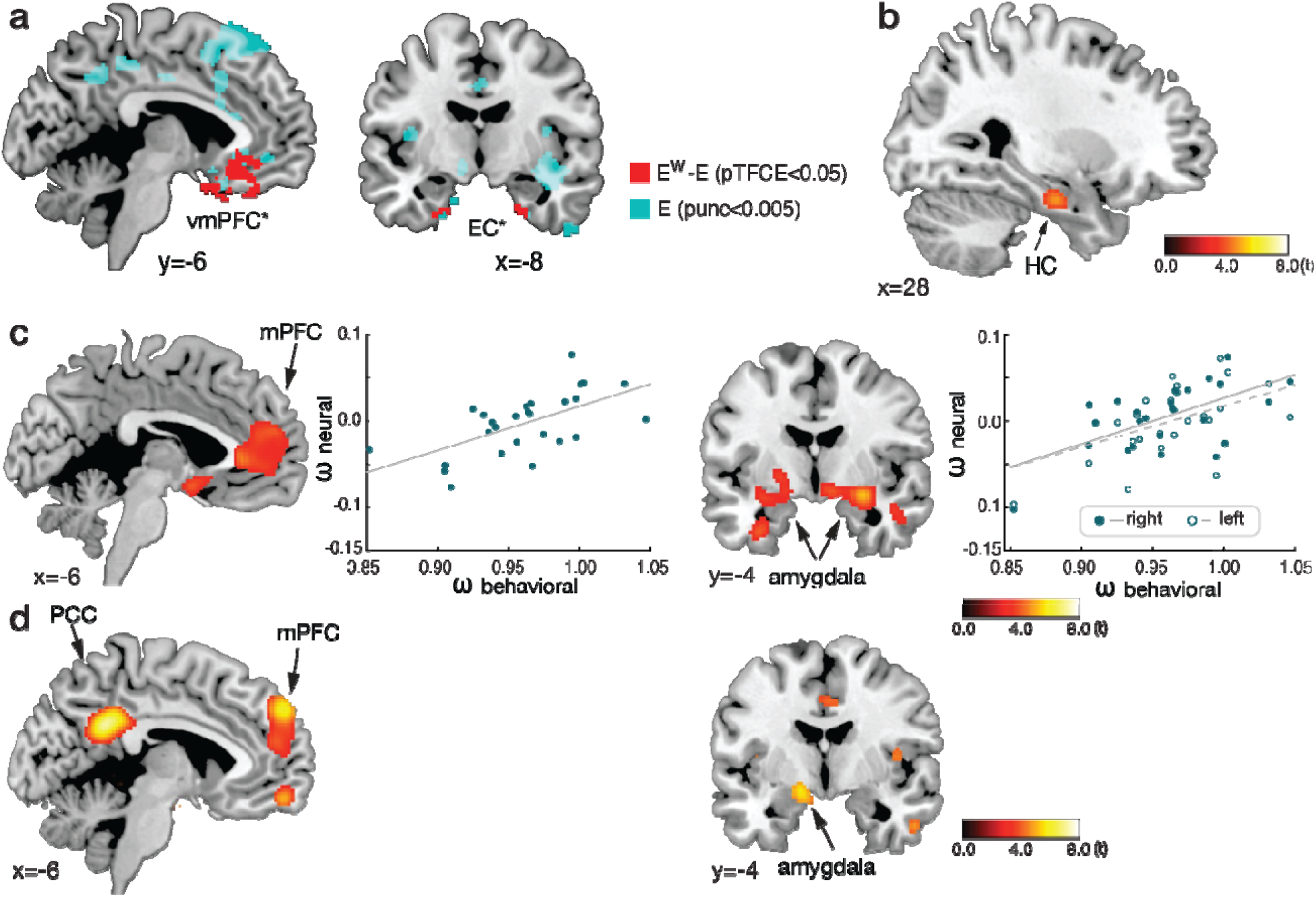
Neural evidence of representational warping. **a**. The entorhinal cortex (EC) and ventromedial prefrontal cortex (vmPFC) encode the Euclidean distance of the inferred trajectory in subjective warped representations. Additional variance in BOLD activity at the time of decisions explained by Euclidian distances in subjective representations (E^W^) over and above the Euclidian distance in the true (non-warped) 4×4 social hierarchy (E) is shown in red (p_TFCE_<0.05 small volume corrected within anatomically defined independent ROIs; denoted by ⍰ next to EC and vmPFC). Within the same model, teal highlights the neural encodings of Euclidean distance (E) based on true (non-warped) representations (*p*<0.005 uncorrected for visualization purposes). These effects control for reaction times (RTs) across trials. **b**. Warping was observed at the group level in HC activity patterns (_Neural_), among other regions, along the context-invariant axis. Maps are displayed at a cluster-corrected threshold of pTFCE<0.05 over the whole brain for all brain regions. **c**. The level of warping (_Neural_) in the medial prefrontal cortex (mPFC) and amygdala correlated with individual differences in behaviorally estimated warping (_Behavioral_). Each dot indicates a participant in the scatter plots. Maps are displayed using conventions in (d). **d**. Intersubject RSA (IS-RSA) analysis showed that the representational geometry in the mPFC, posterior cingulate cortex (PCC), and amygdala reflects individual differences in behavioral warping (_Behavioral_) across participants. Displayed clusters of activation correspond to t-values compared to baseline computed from 1000 permutations with random shuffling at the threshold p_FDR_<0.05.

For the critical test of warped map-like representations in human participants, we directly compare distances between *neural representations* of congruent and incongruent pairs (not trials) across blocks. The results of our RSA of fMRI data consistently showed warping along the congruent compared to incongruent axis — lower pattern similarity between representations of congruent pairs of faces than equidistant incongruent pairs of faces in the true 2-D social hierarchy. Specifically, whole-brain searchlight-based RSA (Supplementary Fig.S3 and Eq.15) of fMRI data revealed this warped geometry at the group level representation in the right HC ([28,-12,-22], *t*=4.50), among other regions (pTFCE<0.05 whole-brain corrected, Fig.5b; See Supplementary Table 4 for a full list of brain areas). The HC effect was also significant after excluding ‘boundary’ individuals who either had the highest or lowest rank in both social hierarchy dimensions (*t*=4.06, pTFCE=0.05 corrected in the anatomical HC ROI), indicating that the observed warping could not be explained by specific faces, such as those at the boundaries (highest or lowest ranks in both dimensions).

Finally, we utilized our inter-individual behavioral warping measure to identify brain regions exhibiting greater warping in participants who showed higher RT costs for incongruent pairs compared to congruent pairs. Specifically, we examined whether participants with greater behavioral warping (*ω*_*Behavioral*_)—indicating a tendency toward longer RTs as the level of context incongruence increased—also demonstrated greater neural warping (*ω*_*Neural*_) in their neural activity patterns, as estimated from RSA (Supplementary Fig.3 and Eq.15), across all participants. This analysis revealed that individual differences in our behavioral measure of warping correlate with the levels of neural warping observed in the amygdala (right [28,0,-12], *t*=6.67; left [-20,-8,-16], *t*=3.30), ventromedial/medial prefrontal cortex (vmPFC/mPFC) ([-5,34,2], *t*=4.52), dorsomedial PFC (dmPFC, [22, 48, 16], *t*=9.90), lateral OFC (lOFC, [34, 32, -18], *t*=4.68), and fusiform gyrus (right [28, -54, -6] *t*=5.88; left [-36,-32,-18], *t*=4.24) (pTFCE<0.05 whole-brain corrected; Fig.5c; See Supplementary Table 4 for a full list of brain areas). This finding suggests those individuals with greater representational warping in these regions experienced greater demands on cognitive control during decisions. This relationship between individual differences in representational geometry and cognitive control during inferences was also supported by the results of another independent analysis, termed intersubject representational similarity analysis (IS-RSA) ^41–43^. The IS-RSA allowed identification of brain areas sharing similar representational geometries of the social hierarchy for participants who showed similar behavioral patterns in cognitive control during inferences, without assuming any specific underlying structure of the representations (Supplementary Fig.S4). Consistent with the previous analysis, we found that individual differences in cognitive control were captured by differences between pattern similarity in the left amygdala ([-16,-2,-12], *t*=5.73), mPFC/dmPFC ([-8,56,38], *t*=6.28), fusiform gyrus (right [24,-50,-6], *t*=7.05; left [-20.-54.-10] *t*=4.30) and posterior cingulate cortex (PCC) ([-8,-48,28], *t*=7.28) across participants (Fig.5d; See Supplementary Table 5 for a full list of brain areas surviving at permutation-based pFDR<0.05 based on 1000 permutations). Control leave-one-out analyses indicated that the results were not driven by the level of warping of any single subject (Supplementary Fig.S4). Collectively, the above lines of evidence converge to suggest people formed warped representations of the 2-D social hierarchy, reflecting the asymmetric demands of cognitive control, and the resulting geometry of subjective representations accounts for individual differences in cognitive control.

#### 3.c. Consistently warped representations emerge in the RNN

To explore under what conditions such representations might emerge during learning, we then performed identical RSA analyses on the hidden layers of the RNN (h_2_ and h_3_ in Fig.1b) throughout learning. Motivated by the human fMRI data, we specifically tested for warping in the 2-D map-like representation in the hidden layers by comparing distances between representations of congruent and incongruent pairs. Consistent with the fMRI results, we found evidence for warping in the hidden layers (*ω*_RNN_; Eq.21) along the congruent compared to incongruent axis that emerged during early to intermediate training timesteps (the peak: training ste*p*=300 [the median run: *t*=6.86, *p*<0.001]; the last training ste*p*=1000 [the median run: *t*=5.29, *p*<0.001]) (Fig.6a). This finding was remarkable, as the warping phenomenon emerged in the RNN simply as a consequence of training it on the same task and without building any additional constraints into the architecture. Notably, warping in the representations persisted even after inference performance had reached an asymptote (Fig.6b). Control analyses indicated that warping in the model’s representations was still present when excluding boundary faces who had the highest or lowest rank in both dimensions (Supplementary Fig.S5). Thus, the warping effect could not be due to the representations of any specific face, nor these outlier face positions. Together, these findings suggest that warping emerges naturally in a standard recurrent neural network trained via backpropagation on our task.

**Figure 6.**
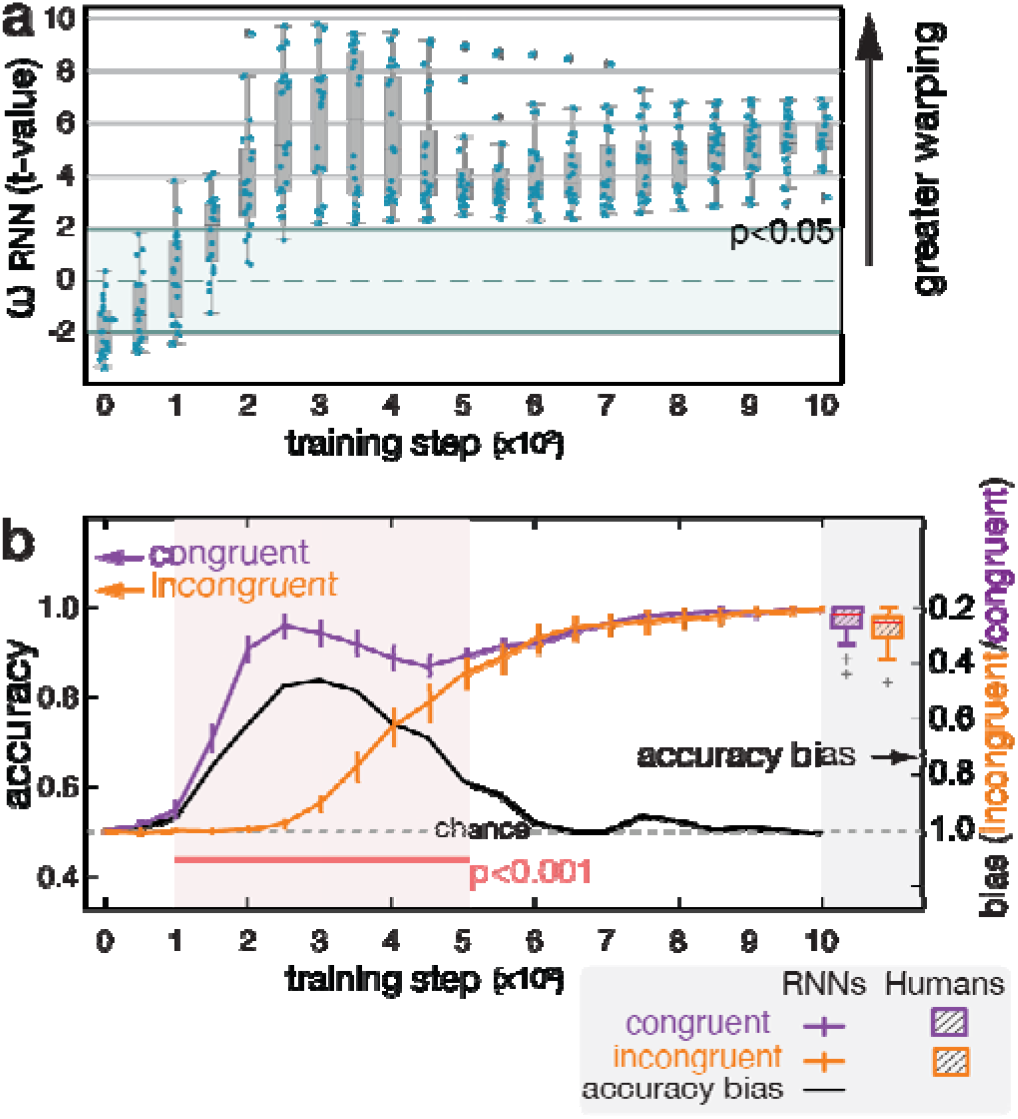
Representational warping in an RNN model. **a**. Timecourse of changes of the level of warping (_Neural_) estimated from the hidden layer representations in the RNN across training. Notably, the RNN representation remained warped even when the model performed inferences with a high-level accuracy. Each point represents an independent RNN training run. At each training stage, 20 independent training runs were performed, with each run initialized using a different random seed. Plotted t values quantify the level of warping _RNN_) and were obtained from two-sided independent-samples t-tests comparing hidden-layer Euclidean distances between congruent and incongruent face pairs (36 pairs each; df = 70) at each training stage. Box, lower and upper quartiles; line, median; whiskers, range of the data excluding outliers; ⍰ sign, the whiskers’ range of outliers. **b**. Timecourse of the mean inference performance (left label) of the RNN across training on the congruent (purple line) and incongruent (orange line) trials. The black line indicates the ratio (right label) between them, indicating the RNN’s relative performance accuracy in the incongruent compared to congruent trials. For comparison, human participants’ accuracy on congruent and incongruent trials is shown as boxplots. The similarity between human and RNN performance supports the interpretation that both reach near-ceiling accuracy with minimal congruency effects. This accuracy bias correlates with timecourse changes in the level of warping in the RNN representation along the context-invariant axis (Fig.6a), suggesting that the tendency to learn congruent compared to incongruent relationships earlier could cause warping in the representations (_RNN_).

We conducted additional analyses to examine whether the key representational findings were robust across different RNN configurations. The results confirmed that the emergence of both 2D and orthogonal 1D representations, as well as the warping effects, remained significant across a wide range of learning rates, hidden layer sizes, and weight initialization gains (Supplementary Fig.S6). These results suggest that the observed representational structures are robust and reflect general computational principles of learning rather than specific parameter choices.

### 4. Inefficient use of context for learning incongruent relationships induces warping

Having found evidence that the brain’s 2-D map-like representation is warped according to subjective control demands, we next examined the hypothesis that cognitive control may determine the level of warping in the subjective representations during learning. Specifically, we examined how incongruence between dimensions experienced throughout learning may cause the observed warping and result in different levels of warping across individuals. We focused on the effects of (in)congruence for the following reasons: On congruent trials, context information that indicates the task-relevant dimension does not strictly need to be maintained for decision making because the correct answer does not depend on the current context. On incongruent trials, on the other hand, context information is required to select the appropriate action (Fig.7a). To investigate why warping emerges, we interrogated the trajectory of the warping in emergent representations in the RNN over the course of training and performed causal experiments to assess the impact on these representations.

Examining performance on congruent and incongruent trials separately revealed that the RNN started performing relatively better on inferences of congruent compared to incongruent trials in the early stages of training (Fig.6b; See Supplementary Table 6 for the results of logistic regression). Moreover, this bias in performance toward congruent trials (black line in Fig.6b) correlated with changes in the levels of warping (*ω*_RNN_) (Eq.21; *β* =0.78±0.10 (SE), *t*=7.85, *p*<0.001). While these effects were relatively large early in training, the representations remained warped even after the RNN’s performance equated for congruent and incongruent trials (Fig.7b). This observation suggests that the warping in the RNN might be linked to an initial tendency to ineffectively utilize context information that would mitigate the interference on incongruent trials. To test the relationship between the use of context information and warping in the RNN, we performed an ablation experiment. To mimic individual differences in the insufficient use of context information, we degraded the embedding vector of the context cue (by setting *α* to 1 to 10; *e*_*c*_ in Fig.1b and Eq.20) during both training and testing, and examined changes in the geometry of the hidden layer’s representations of the faces (h_2_ and h_3_ in Fig.1b and Eq.17). The results of the ablation experiment showed increases in warping in the representations as a function of the degree of degraded context cue information (*β* =-0.97± 0.20 (SE), *t*=-4.84, *p*<0.001) (Fig.7c), consistent with the notion that the inefficient use of context produced the observed warping. To test whether the observed representational warping emerged during training or testing, we selectively degraded the context input either only during training or only during testing. These analyses showed that degrading context input during training specifically induced warping in the representations, whereas degrading context during the test phase had minimal influence on representational warping (Supplementary Fig.S7). As predicted, degrading context input only during test selectively impaired performance on incongruent trials, with minimal impact on congruent trials, supporting the need to use context information specifically during incongruent inferences. With minimal changes to the representations during testing, behavioral performance during inference depended primarily on the representational structure formed during training and the use of context information.

**Figure 7.**
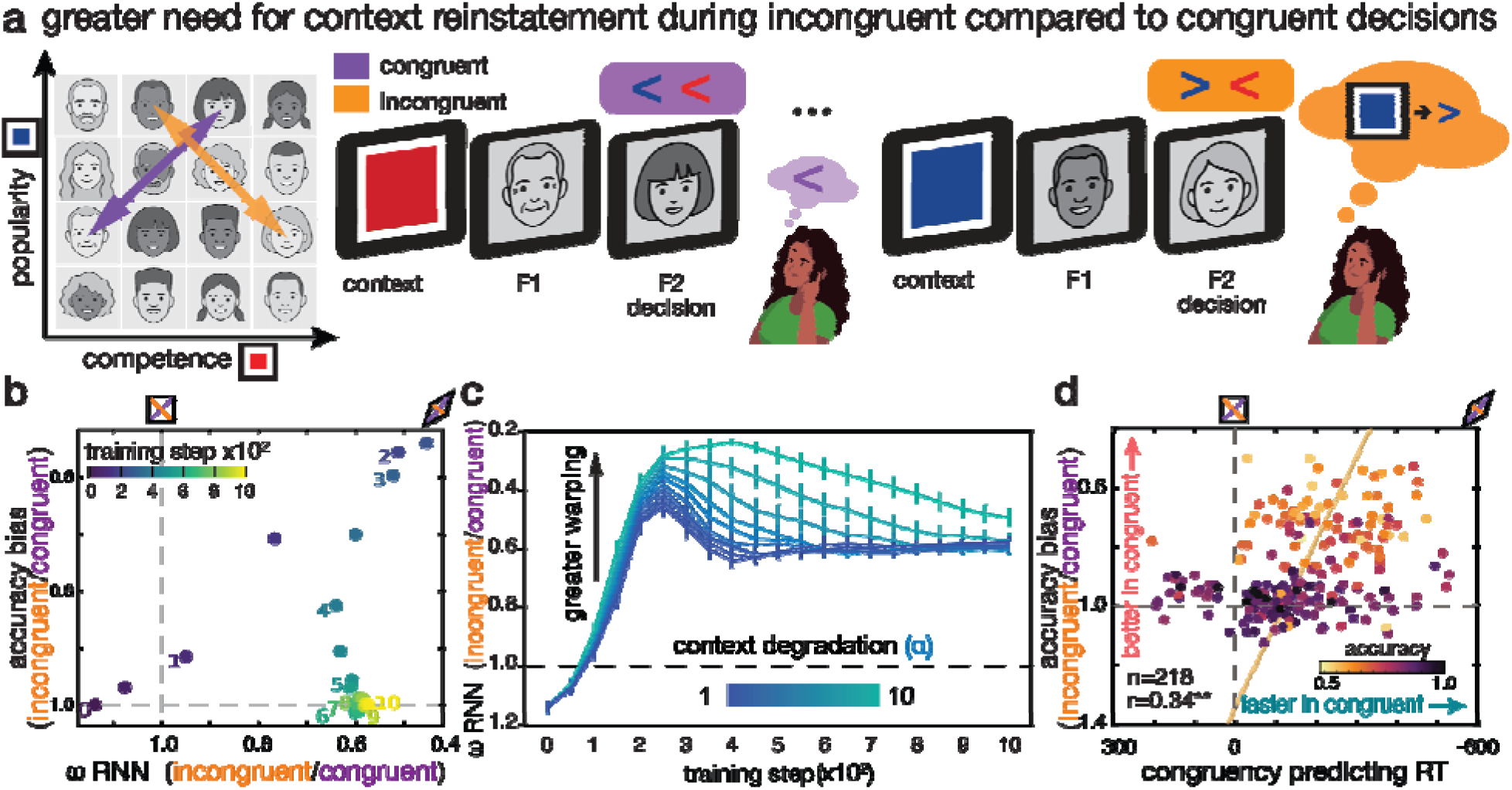
Relationship between context and warping in subjective representations. **a**. Conceptual schematic illustrating the hypothesis that the context cue information is reinstated on incongruent decisions to overcome interference between conflicting responses. **b**. The trajectory of the accuracies and accuracy bias between incongruent compared to congruent inferences for the RNN as a function of the level of warping measured with RSA of hidden layer units (_RNN_). Each dot indicates the mean accuracy bias across runs at different training steps (0: the beginning to 10: the last training step) and their mean level of warping at the same training step. **c**. Results of ablation simulation. Each line represents the mean level of warping (RNN) across n = 20 independent RNN runs (simulation replicates, each initialized with a different random seed) at a given level of context degradation (= 1–10; Fig.1b; Eq.20), plotted across training steps. Greater degradation of the task-relevant context information in the RNN (increases in in Fig.1b) incurs greater warping in the 2-D representation in the RNN across training steps (increasingly light blue lines), supporting the causal influence of accurate task-relevant context information on the construction of less warped (veridical) 2-D representations (Fig.2a). Error bars indicate SEM. Dashed line indicates no warping. **d**. Relationship between individual differences in accuracy bias during behavioral training (n=218) and the regression coefficient for context congruence predicting RTs from a multiple linear regression controlling for other distance measures, including decision difficulty (analogous to Fig. 4a). Participants with greater RT advantages for congruent over incongruent trials, indicative of greater representational warping, showed greater accuracy for congruent but lower accuracy for incongruent relationships. This relationship reflects more than simply faster RTs for congruent inferences; instead, it indicates that subjective representational warping, indirectly inferred from RT patterns, selectively biases inference accuracy, analogous to the relationship between representational warping and accuracy bias observed in the RNN (Fig. 7b). Darker purple dots indicate greater accuracy. Vertical and horizontal dashed lines indicate no accuracy bias and no RT advantage for congruent trials, respectively.

Inspired by this observation in the RNN model, we performed additional behavioral and fMRI analyses. First, in a larger cohort of participants trained behaviorally, we found that participants whose accuracy was more asymmetric (higher in congruent compared to incongruent pair trials) likewise showed RTs that were more strongly explained by the effects of congruence, potentially consistent with more warped representations (*r*=0.34, *p*<0.001, *n*=218; Fig.7d and Supplementary Fig.S8). Participants with lower behavioral performance exhibited stronger congruency effects in RT and were more likely to fail on inferences involving incongruent relationships. This relationship remained significant after controlling for each participant’s tendency to rely on a warped representation (partial *r*=0.25, *p*<0.001). This tendency was estimated using a logistic regression predicting choice, as the relative contribution of context-independent decision values, reflecting a fully warped representation (Fig.2a, left), compared with context-dependent decision values, reflecting an unwarped representation (Fig.2a, right). These individual differences likely reflect variability in the use of context information to learn the two task dimensions independently. Insufficient use of context leads to greater representational warping and selective performance biases in incongruent trials. This result and our RNN analyses point to a key role for effectively using context information to guide decisions specifically on incongruent trials.

Second, we tested the hypothesis of greater neural reinstatement of the current context information, specifically at the time of decisions, on incongruent than congruent trials. We first defined ROIs that showed significant context representations at the time of the context cue presentation, which included the bilateral HC, mPFC, anterior insula, dorsal anterior cingulate cortex (dACC)/pre-SMA, retrosplenial cortex (RSC)/PCC and premotor cortex (pTFCE<0.05 whole-brain corrected). Next, we tested for stronger reinstatement of these context-specific activity patterns at the time of decision making (F2 presentation) on incongruent trials than congruent trials, which identified significant clusters in the left HC ([-26,-8,-22], *t*=3.25), mPFC([-12,48,-4], *t*=3.19), dACC/pre-SMA([-6,12,44], *t*=6.59), anterior insula ([-30, 18, -8], *t*=5.48 in right and [36, 22, -2], *t*=6.44 in left), and RSC/PCC ([-4,-28,28], *t*=6.35), among other areas (pTFCE<0.05 corrected in the combined ROI) (Fig.8a; See Supplementary Table 7 for additional regions outside of these ROIs). Importantly, this GLM controlled for the potential effects of the context, incongruence, and motor responses by including these as control RDMs (Supplementary Fig.S9). These findings support the hypothesis that there is greater reinstatement of context information during inferences in the presence of incongruence, precisely when it needs to be used to overcome interference.

**Figure 8.**
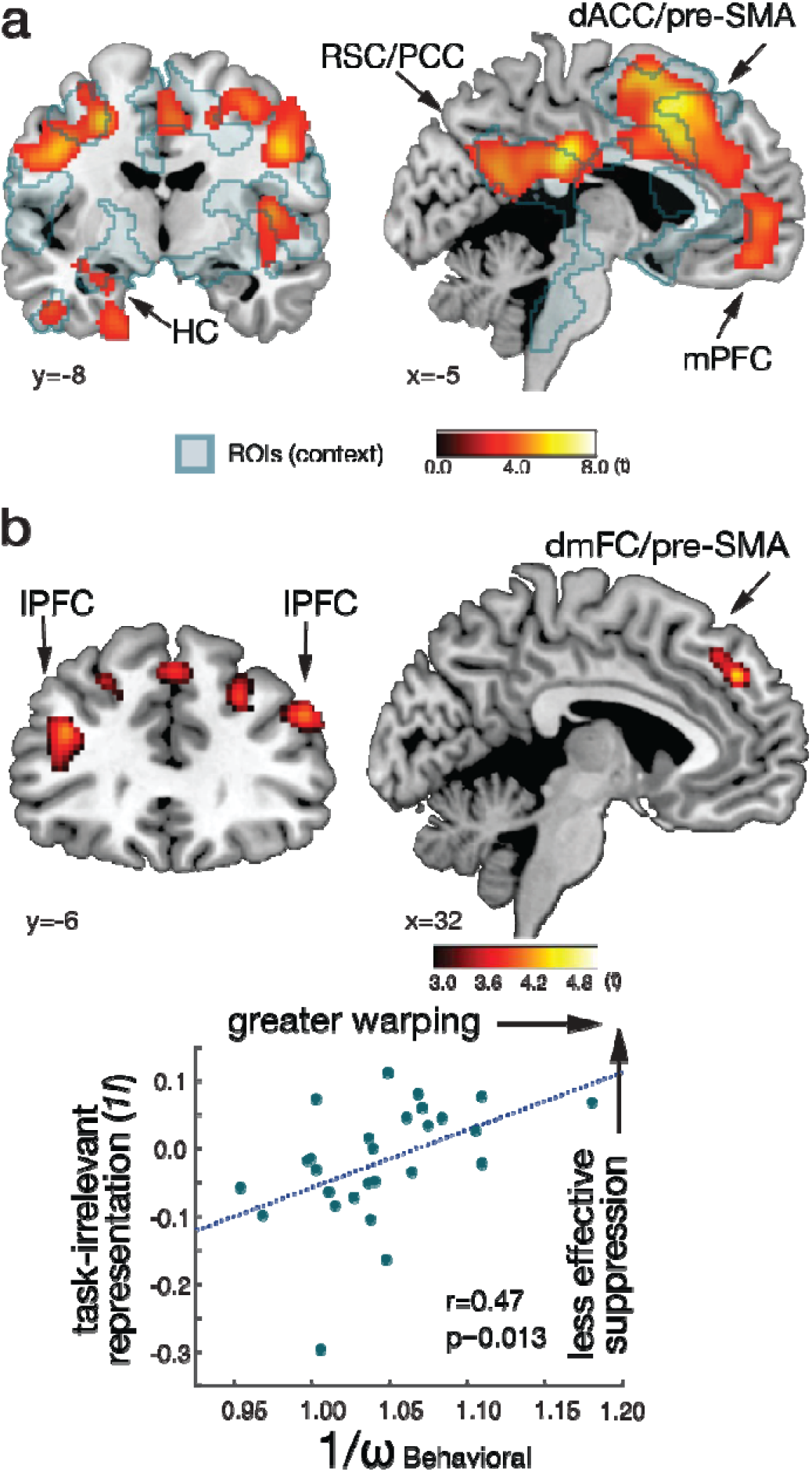
Context reinstatement and representational warping in neural representations. **a**. T-statistic map showing activity patterns specific to the current task-relevant context information were reinstated greater at the time of inferences on the incongruent than congruent trials in the hippocampus (HC), medial prefrontal cortex (mPFC), dorsal anterior cingulate cortex / pre-supplementary motor area (dACC/pre-SMA), and retrosplenial cortex / posterior cingulate cortex (RSC/PCC). Statistical significance was assessed using one-sided one-sample t-tests on the contrast images. The maps are displayed at a cluster-corrected threshold p_TFCE_<0.05 over the whole brain. The independently defined ROIs including all brain areas showing context-specific activity patterns at the time of context cue presentation are shown in green outline. **b**. Left: T-statistic map showing regions with a significant correlation between the neural representation of the task-irrelevant rank () and the behaviorally estimated level of warping (_Behavioral_) across participants. Statistical significance was assessed using one-sided one-sample t-tests on individual participants’ RSA maps. The map is displayed with a threshold of *p*<0.005 (uncorrected). Right: Scatterplot illustrating the correlation between the behavioral index of warping and the task-irrelevant dimension information. The level of 1*I* in the scatter plot is estimated from a functional ROI defined from the results of an independent analysis that identifies brain areas containing task-relevant information within the 1-D orthogonal representation (Fig. 2c).

Third, context input is essential for effective compression in the task-dependent 1D representation. Without it, the representation cannot be appropriately compressed, and the brain cannot extract information along the task-relevant dimension. Based on this logic, we reasoned that those participants showing smaller behavioral costs from the task-irrelevant dimension on incongruent relative to congruent trials (i.e., less behaviorally implied warping) would exhibit less intrusion of the task-irrelevant dimension in the 1D orthogonal representations within the frontoparietal network. As expected, we found a positive correlation between the behaviorally estimated level of warping in the 2-D representation (*ω*_Behavioral_) and the extent to which the neural activity patterns were explained by the task-irrelevant information in the 1-D context-dependent orthogonal representation in the dmFC/pre-SMA ([26,24,50], *t*=3.78]) and lPFC (left: peak [-42,22,42], *t*=4.74; right: peak [50, 38, 30], *t*=4.1; pTFCE<0.05 corrected in combined ROI) (Fig.8b and Supplementary Table 8). This finding suggests that participants who were better at forming accurate, less warped 2-D representations of the task showed neural activity patterns that reflected a greater ability to suppress task-irrelevant information in these regions. This mechanism likely supported their greater flexibility in decision making across different contexts. Together with the RNN and fMRI analyses, these findings suggest a critical link between the utilization of contextual information for effective cognitive control and the observed representational warping.

## Discussion

Our findings show how latent non-spatial, abstract cognitive maps are represented when learning relationships piecemeal from two different task contexts, and how they are used to make flexible decisions in only one task-relevant dimension at a time. To address these questions, we performed parallel analyses of an RNN model and human fMRI data and have shown three key phenomena about the relationships between the representational geometry of cognitive maps interacting with changing task goals and ensuing cognitive control demands. First, both a HC-EC-OFC/vmPFC network and the RNN learn multidimensional (2-D) representations of the latent task structure such that the relational information learned from different task contexts were represented as different axes of the same 2-D representation that was shared across contexts. Second, to facilitate effective selection of the task-relevant dimension, this multidimensional representation was modified into one of two orthogonal 1-D rank representations, suppressing the task-irrelevant dimension in a frontoparietal cortical network and in the RNN. Finally, individual differences in cognitive control—likely operating during learning and the formation of context-specific representations—were associated with greater warping along the context-invariant (congruent) axis, and this warped neural geometry was in turn associated with increased behavioral control demands during incongruent inferences. The current findings characterize cognitive control as a process whereby an individual’s representational geometry is both sculpted by and shapes cognitive control when retrieving specific dimensions of endogenous representations from memory.

Although human participants were never explicitly instructed about the underlying latent structure, representations in HC, EC, and OFC/vmPFC captured this structure in that the similarity between representations of each entity (individual faces) was correlated with their true pairwise Euclidean distances in the 2-D social hierarchy ^11,12^. The stable 2-D representations support efficient generalization across multiple task contexts. By encoding relationships between entities according to their ranks within a common multidimensional space, the brain can infer novel relationships along different dimensions without needing to switch representations based on current task demands. In addition, we also found simultaneous 1-D representations in which the axes of the two different task dimensions were represented orthogonally when they were behaviorally-relevant, with the irrelevant dimension compressed, in a separate network of brain regions including the dmFC, lPFC, PCC, and IPL. Importantly, these orthogonal 1-D representations afford behavioral flexibility by minimizing interference due to compression of task-irrelevant dimensions relative to task-relevant dimensions. This finding suggests that the human brain may use similar principles for top-down biasing or selection of task-relevant from irrelevant dimensions as previously proposed for explicitly presented perceptually signaled features ^8,9^ to perform inferences over particular task-relevant dimensions of cognitive maps retrieved from memory. Notably, we also found both 2-D and orthogonal 1-D representations emerge in the representations of faces in the hidden layers of our RNN, with distinct time courses. Consistent with the representations found in the human brain, task-general 2-D representations formed in the RNN at earlier phases of training and remained even after extensive training. The RNN more gradually developed the 1-D orthogonal representations as training progressed, and this tendency to utilize task-specific representations increased throughout training.

Our results demonstrate that the representational transition previously observed in perceptual context-dependent decision-making extends to memory-guided decision-making, in which decision values must be inferred from mnemonic relationships rather than selected from perceptual features. Previous work ^8,9,25,26^ showed that context-dependent behavior can be supported by compressed low-dimensional representations that are orthogonal across task contexts, allowing task-relevant information to be accessed while minimizing interference. Our findings are consistent with this orthogonal-coding principle, but extend it in two key ways. First, because faces carried no intrinsic feature-to-rank mapping, participants had to infer values from pairwise-learned relational structure rather than from perceptual dimensions. Second, we found that task-dependent orthogonal 1D codes in frontoparietal regions coexisted with task-independent 2D relational representations in HC-EC and vmPFC. This suggests that memory-guided decisions may rely on high-dimensional relational maps as internal models that are compressed or selectively accessed into orthogonal low-dimensional subspaces according to current task demands.

While other studies have observed multidimensional task structure representations ^5,11,12,31,44^, little is known about the factors determining the representational geometry of such cognitive maps. If neural systems construct separate representations for separately learned task dimensions, these could afford behavioral flexibility but in theory, should incur costs for task switching between different cognitive maps. On the other hand, having a task-independent representation shared across multiple dimensions (a stable multidimensional representation) (Fig.2a) allows for fast learning in a new task context via generalization (which works particularly well for learning congruent relationships in our task) and a single computation across dimensions, thereby mitigating costs of task/context switching ^3,22,24^. Our results indeed showed that both medial temporal and interconnected orbital and ventromedial prefrontal brain regions ^45^ and RNNs can construct a shared multidimensional representation that can serve generalization across dimensions, and flexible readout of information depending on the current goal. In the context of our task, computing shared decision vectors across dimensions is beneficial for inferences on congruent relationships but not for inferences on incongruent relationships. Convergent pieces of evidence suggest that an initial shared representation developed at early training steps more likely represented relationships in different dimensions along a collapsed low-dimensional number line. This representation started to transition toward orthogonal high-dimensional representations wherein the relationships in the two dimensions became represented on separate axes of a single 2-D representation (Fig.c, 3i and 3j). Specifically, we found that those participants who showed greater accuracy asymmetries on incongruent compared to congruent trials also exhibited RTs better explained by the context-independent rank differences (or context-free values based only on win/loss ratio regardless of the task-relevant dimension), whereas RTs of those who showed lower asymmetries were better explained only by the context-dependent task-relevant values (Fig.7d and Supplementary Fig.S8). Complementing these behavioral data, the simulation results from the RNN also support this interpretation by showing that the shared representations that initially enabled accurate inferences on the congruent relationships better than incongruent relationships evolved into a 2-D structure enabling accurate inferences on both with increased training (Fig.7b). This finding suggests that learning how many dimensions are relevant to potential behavior and how they are related to each other gradually transforms the representational geometry. We acknowledge the possibility that factors beyond value mapping influence this transformation. For example, prior work shows that response mappings can bias how learners organize task structure ^46,47^, raising the possibility that alignment or conflict between response and value mappings may also shape representational dimensionality. Taken together, these and our findings imply that the dimensionality of the task stimulus space and response mappings both likely contribute to the neural representational geometries that emerge, as opposed to the view that tasks containing two relevant dimensions are naturally mapped onto rigid 2D representations in the hippocampal formation.

Our findings contribute to the debate over whether task representations in the brain are high- or low-dimensional, suggesting that these formats are not mutually exclusive but rather complementary and potentially co-evolving. A high-dimensional relational scaffold may arise from representations initially constructed across combined dimensions to encode multiple task features along orthogonal axes, while concurrent compression into low-dimensional, task-specific formats promotes unbiased and accurate encoding that supports context-dependent decision-making. Consistent with this view, both two-dimensional and orthogonal one-dimensional representations emerged naturally in our human data and in the RNN models without requiring additional architectural or cost-function constraints. The RNN in the current study was designed to test how the task’s learning demands shape internal representations and performance, while leaving response speed outside the scope. Future work could incorporate evidence-accumulation mechanisms ^48^, which model RTs through gradual evidence accumulation toward a decision, to examine how learned representations are translated into choices and RTs. In the present model, representational warping emerged from the learning asymmetry itself: congruent associations aligned with prior structure, whereas incongruent associations deviated from it and imposed greater learning demands. This suggests that multiple representational formats can coexist and jointly contribute to inference. It is also worth noting that the current findings cannot resolve which representational format—2D or 1D— contributes more strongly to correct inferences, nor whether their relative influence changes across learning. An important direction for future work will be to test whether, when, and how these high- and low-dimensional structures interact, compete, or complement one another during learning. Such investigations will be critical for revealing how multiple representational formats jointly support flexible cognition.

Representational warping in the present study arises from the demands of learning an unknown decision structure under competing decision policies—a challenge common in everyday cognition, where latent relationships must be inferred from experience rather than navigated within a known space. Unlike tasks in which the two dimensions are explicitly defined as independent (e.g., spatial navigation or orthogonal feature spaces) ^49–54^, our task required participants to learn each dimension in isolation, without ever experiencing them simultaneously, and subsequently integrate these separately acquired relationships to construct the latent two-dimensional structure. This learning process naturally produced congruent and incongruent relationships, requiring the resolution of interference between competing relational mappings. These demands closely parallel classic cognitive control tasks, in which selecting task-relevant information requires suppressing competing alternatives ^8,9,25–27^. We note, however, that such distortions could in principle also reflect experience-dependent value learning inherent to the task structure ^*55,56*^. Our interpretation does not exclude this possibility; rather, we propose that individual differences in how effectively participants use task context to constrain learning determine the extent to which these experience-driven biases become incorporated into the learned representational geometry.

We tested for the effects of congruence on the representational geometry of cognitive maps in the human brain by comparing distances between representations of congruent and incongruent pairs sampled from different trials across blocks. We found warping in the HC representations along the congruent compared to incongruent axis □ greater differences in pattern similarity between representations of equidistant congruent pairs of faces than incongruent pairs of faces. Additionally, the levels of warping estimated in the amygdala, mPFC, and PCC were correlated with individual differences in our behavioral measure of cognitive control, suggesting those individuals with greater representational warping in these regions experienced greater demands on cognitive control during decisions. These results were also supported by an independent IS-RSA analysis. These findings accord with previous studies which have reported sensitivity to transitively inferred social hierarchy rank in activity in the amygdala, HC, and mPFC, and with evidence amygdala gray matter volume correlated with individual differences in transitive inferences in the social domain ^57,58^ and social group size ^59–63^.

Our findings indicate that both the utilization of context and the suppression of irrelevant information play a vital role in maintaining the integrity of representations. We observed a correlation between participants who displayed less bias in their response times, suggesting a lower degree of distortion in their representations, and their enhanced ability to suppress task-irrelevant information in brain regions that contain a one-dimensional task-dependent orthogonal representation. This suggests that the failure to fully employ contextual information can hamper the ability to suppress irrelevant dimensions during the process of making inferences, consequently impacting the level of cognitive control required. Previous studies have demonstrated the distortion of value comparison processes caused by task-irrelevant information ^64–67^. Our findings extend this understanding by demonstrating that the insufficient suppression of task-irrelevant dimensions can lead to distortions in the representational geometry. This effect is particularly pronounced when decision values are not exclusively determined by the explicit features of stimuli but rather rely on the retrieval of hidden relational structures from memory.

Our findings further suggest that the warped representations along the congruent versus incongruent axis further influence the control demands which arise from interference between conflicting responses. To resolve this conflict, the brain needs to deploy control resources to select the single task dimension relevant to the current goal signaled by one of two discrete color cues—a process consistent with models emphasizing strengthened context representations under conflict ^*1,68,69*^. The results of the ablation simulation support the importance of contextual representations by showing that increasingly degrading the contextual input impaired learning of incongruent relationships in particular and subsequently caused greater warping in 2-D social hierarchy representations. We found consistent findings in the human fMRI data: the HC, mPFC, anterior insula, and RSC/PCC representations specific for a single contextual cue were reactivated to a greater extent during inferences on incongruent compared to congruent pairs. Importantly, among these ROIs, the level of reactivation in the HC specifically correlated with the level of warping estimated from behavioral RTs across participants. This suggests that efficient use of contextual information resolves conflict by orthogonalizing dimensions in the shared representations, and the subjective representational geometries likely affect the control demands required for flexible decision making. An intriguing open question for future studies concerns how the contextual representations and 2-D map-like representations are related during goal-directed selection of specific information from cognitive maps. Here, we estimated the geometry of the latent 2D space indirectly through decision-time patterns rather than through an independent behavioral reconstruction of the latent 2D representational space; complementary approaches using explicit similarity judgments or spatial reconstruction tasks may further refine these estimates in future work.

We found evidence of complementary representational geometries for efficiently encoding abstract relational information and flexibly selecting behaviorally-relevant dimensions from those representations in the human brain as well as the neural network model. Our findings suggest that cognitive control shapes the representational geometry of cognitive maps which can create warping in the subjective representations. We further showed that biases in context-dependent inferences could arise from the subjective representational geometry of cognitive maps. This insight provides an approach to better discern individual differences in learning and decision making, and could provide a framework to understand and potentially overcome irrational biases in decision making, social prejudices, and psychiatric conditions ^70–73^. Moreover, the current study demonstrates the value of integrating a neural network modeling approach with neuroimaging through the interrogation of representations that emerge through learning, and may help to address current limitations of modern neural networks used for artificial intelligence ^74,75^, which often exhibit excessive reliance on contextual information ^76^, and lack human-like control processes ^77^.Taken together, our results reveal an intricate relationship between cognitive control during cognitive map formation, the resulting representational geometry, and its role on subsequent control during decisions.

## Methods

### Participants

33 participants (age range: 19–23, normal or corrected to normal vision) were recruited to the fMRI experiment via the University of California, Davis online recruitment system. Six participants were excluded due to strong head movements (> 3 mm) and technical failure (2 subjects). In total, we analyzed the scans of 27 participants (16 female, mean age: 19.37±0.26, standard error mean [SEM]). The study was approved by the University of California, Davis, IRB. Participants provided written informed consent before the experiment and received course credit and performance-based monetary compensation for their participation. The sample size was estimated based on a power analysis assuming a medium to a large effect (d = 0.6), resulting in a sample size of 24 to achieve a statistical power of 80% (a = 0.05, two-tailed test). Participant sex was self-reported at recruitment; gender was not collected. Sex and gender were not considered in the study design, and no sex- or gender-based analyses were performed because the study was not powered to detect such differences. All participants completed the same within-participant task. Blinding was not applicable because there were no experimental group allocations.

### fMRI experiment

In the current study, we performed additional analyses using the dataset collected in a previous study (see ^11^ for detailed information on the experiment design). During fMRI participants made novel inferences about the relationships between pairs of individuals in one of two social hierarchy dimensions – “competence” and “popularity.” In each trial of the fMRI experiment, participants were given a context cue (1 s) with the color indicating the social hierarchy dimension that relevant to the task in the current trial, and two face stimuli (F1 and F2) followed sequentially. Each face stimulus was shown for 2 s. Participants were asked to make a binary decision by selecting the face associated with a higher rank in the given social hierarchy dimension while F2 was on the screen. The experiment used photographs of human faces; however, to protect privacy, the face stimuli shown in this article have been replaced with author-created illustrations. Before fMRI, participants learned the relative ranks between these 16 individuals in two social hierarchy dimensions. Unbeknownst to participants, F1 and F2 were selected from the pairs that had not been directly compared in the given dimension during training. This allowed us to present 12 faces while excluding 4 “hub” faces (see behavioral training) as F1 or F2 in each of two social hierarchy dimensions. To make a correct inference, therefore, participants needed to make transitive inferences using comparisons that they have been trained on. Importantly, we limited the number of pairs that participants learned about during training. This forced participants to make novel inferences through a unique trajectory. As a control experiment, at the end of each trial, a third face was shown and participants were asked to press a button according to the gender of the face. The inter-stimulus interval (ISI) was 1.5 s between the context cue to F1 and 2-5 TRs between faces. The inter-trial interval (ITI) was 2-5 s. To acquire responses to all inferences, the trial in which participants did not make a decision was presented again after a random number of trials. The fMRI experiment comprised two 104 trial blocks. The presentation order was randomized across participants.

### Behavioral training

Participants were asked to learn the relationships of 16 individuals in two social hierarchy dimensions for 3 days of behavioral training. The training was comprised of learning and test blocks. For the first two days, participants learned the relationships in each of the two social hierarchy dimensions on different days. The dimension learned on the first day was counterbalanced across participants. During behavioral training, participants were presented with a pair of individuals and asked to select the one with a higher rank in the given social hierarchy dimension. During learning blocks, participants received feedback on their decisions. No feedback was given during the test blocks. Importantly, during behavioral training and the fMRI experiment, no information hinting at the structure of the social hierarchy was given to participants. Instead, participants were only informed that the pairs shown in the learning blocks have one rank difference in the given dimension. This allowed participants, in principle, to infer the overall social hierarchy structure through transitive inferences. During test blocks, to test if participants could infer the relationships that were not directly compared during learning, participants were asked to infer relationships of pairs of individuals whose ranks differed by one or more rank differences. At the end of day 2 training, after learning/testing relationships between individuals in both dimensions, participants performed an additional test block (called “flexible test”) in which participants make inferences of relationships between pairs in the given social hierarchy dimension while the task-relevant dimensions were randomly interleaved across trials. Participants who successfully inferred the relationships in both social hierarchy dimensions and exceeded the predetermined accuracy threshold (participants who successfully distinguished the second- and third-rank individuals above chance and also reached > 85% overall accuracy in the flexible test) continued to participate in the day 3 training. This threshold was determined and used in order to exclude participants who may be using heuristics such as wins/losses regardless of the current task-relevant dimensions rather than building relational structure to infer the unlearned relationships. The 3 days of behavioral training created a relational structure of 16 faces in two social hierarchy dimensions. The inferred trajectories between individuals were determined by the learned relational structure. The details in behavioral training can be found in the protocol paper ^37^. Unbeknownst to participants we paired an individual only with a limited number of other individuals during the first two days of training. This inherently created two eight members groups (learning group). That is, an individual was compared only to the same group members but not to those in the other group. During day 3 training, right before the fMRI scanning, participants learned between-group relationships only from selected pairs for the first time. In particular, only certain faces called “hubs” were paired with members of their own group (on days 1 and 2) as well as members of the other group (on day 3). To infer the unlearned between group relationships, therefore, participants need to make transitive inferences through a unique hub face paired with both group members. This allowed us to create a unique trajectory for a transitive inference during the fMRI experiment. Among 282 participants who initially signed up for the participation, 218 participants completed the first two days of training. Among 85 participants whose accuracy exceeded the predetermined threshold, 65 participants continued to the day 3 behavioral training. Of the original 65 participants, 51 demonstrated performance above a predetermined threshold on the inferences during the final training session on day three. Thirty-three of these 51 participants then went on to participate in the fMRI experiment portion of the study. Separately, 18 of the original 51 participants who reached criteria during behavioral training took part in a behavioral-only version of the inference task. Refer to Park et al. ^11^ for further details on participant retention.

### Functional imaging acquisition

We acquired T2-weighted functional images using Siemens Skyra 3 Tesla scanner. We used gradient-echo-planar imaging (EPI) pulse sequence, with a multi-band acceleration factor of 2. We set the slice angle of 30° relative to the anterior-posterior commissure line to minimize the signal loss in the orbitofrontal cortex region (Weiskopf et al., 2006). The 38 slices were acquired with 3mm thickness with the following parameters: repetition time (TR) = 1200 ms, echo time (TE) = 24 ms, flip angle = 67°, field of view (FoV) = 192mm, voxel size = 3 × 3 x 3 mm^3^. Contiguous slices were acquired in interleaved order. We used a field map to correct for potential deformations with dual echo-time images covering the whole brain with the following parameters, TR = 630 ms, TE1 = 10 ms, TE2 = 12.46 ms, flip angle = 40°, FoV = 192mm, voxel size = 3 × 3 x 3 mm^3^. A T1-weighted structural image was acquired using a magnetization-prepared rapid gradient echo sequence (MPRAGE) with the following parameters: TR = 1800 ms, TE = 2.96 ms, flip angle = 7°, FoV = 256mm, voxel size = 1 × 1 x 1 mm3.

### Pre-processing

The preprocessing of functional imaging data was performed using SPM12 (Wellcome Trust Centre for Neuroimaging). Images were corrected for slice timing, realigned to the first volume, and realigned to correct for motion using a six-parameter rigid body transformation. Inhomogeneities created using the phase of nonEPI gradient echo images at 2 echo times were coregistered with structural maps. Images were then spatially normalized by warping subject-specific images to the reference brain in MNI (Montreal Neurological Institute) coordinates (2mm isotropic voxels).

### Testing simultaneous context-invariant and context-dependent representations

We performed a whole-brain searchlight representational similarity analysis (RSA) ^78,79^ to test whether the brain represents the 2-D context-invariant and 1-D orthogonal context-dependent representations simultaneously. Specifically, we estimated the mean activity patterns associated with each of the faces during correct inference trials, combining F1 and F2 presentations. This was accomplished using a general linear model (GLM) while including the onsets of the two contexts and button press, as well as the 6 motion regressors as regressors of non-interest. The mean activity patterns were separately estimated for each face identity and for each task context, corresponding to the social hierarchy dimension relevant to the current goal. By comparing the context-dependent activity patterns of faces in one run with those of another independent run, we eliminated the impact of temporal autocorrelation on our estimation of pattern dissimilarity. Based on these representations, we computed the representational dissimilarity matrix (RDM) per region of interest (ROI) across the whole brain. We quantified the representational dissimilarity using the Euclidean distance between the standardized mean activity patterns of representations obtained from two independent fMRI blocks. To assess representational structure, we computed cross-run RDMs, comparing beta estimates for each condition across independent blocks. Each stimulus was presented an equal number of times across runs and participants, minimizing potential biases from unequal sampling. To control for differences in magnitude between neural patterns, we applied z-scoring to the mean activity patterns evoked by each face in each task context. We subsequently predicted the RDM estimated from the patterns of neural activity in each of the ROIs using a GLM in which we included six idealized model RDMs (Supplementary Fig.1 and Eq.1). While our group-level analysis relies on point estimates of RDM–model correlations and does not incorporate uncertainty in beta estimates, this two-step approach has commonly been used in representational similarity analysis ^*78–80*^. We acknowledge that hierarchical modeling may offer advantages in explicitly modeling uncertainty and individual variability ^81^, and we consider this a valuable direction for future work.

The six model RDMs include: (1) 2*D*: Pairwise Euclidean distances between faces in a context-invariant 2-D social hierarchy. This RDM allows us to identify brain areas whose activity patterns represent a stable 2-D social hierarchy; (2) 1*D*: Pairwise distances between individuals in a 1D representation where the two task dimensions are encoded along orthogonal axes, with distances in the task-irrelevant dimension suppressed. This RDM allows us to identify brain areas whose activity patterns form a 1-D social hierarchy that changes across contexts, focusing on task-relevant rank differences. This representation assumes that the brain uses distinct sets of codes to represent rank along the current goal-relevant dimension, while compressing ranks of goal-irrelevant information along an orthogonal axis. Under this model, stimuli with the same rank in different contexts would not share neural representations, and the overall representational geometry would differ across contexts; (3) 1*R*: Pairwise distances between individuals in a 1D representation where the two task dimensions are encoded along a single shared axis (Akin to a number line). This representation posits that the brain uses a single, shared axis to encode ranks in the dynamically changing goal-relevant dimension, while its representations are insensitive to ranks in the goal-irrelevant dimension. As a result, in the 1*R* representation, stimuli with the same rank in different contexts are predicted to show similar neural representations, leading to a common representational topology across contexts. (4) 1*I*: Pairwise distances between individuals in a 1-D representation where distances in the task-relevant dimension are suppressed; (5) *Context*: Indicates if faces were from the same or different task-relevant social hierarchy. This captures whether activity patterns for two faces were measured under the same or different social hierarchy dimension; (6) *Group*: Indicates if ranks of two faces were compared within the same training group.

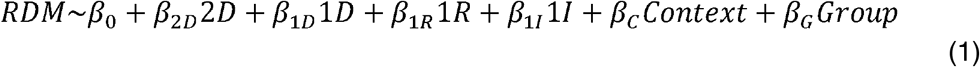

Fig. 2a and Supplementary Fig.1 depicts the difference between 1*D* and 1*R*. The two RDMs, 1*D* and 1*R* test different hypotheses about the geometry of the 1-D social hierarchy representation across contexts: 1*D* tests the hypothesis that the geometry changes across contexts, while 1*R* tests the hypothesis that the geometry remains stable across contexts. When comparing faces within the same context, predictions from the 1*D* and 1*R* models are the same because both represent ranks of faces within the same task dimension. The critical test lies in how representations change across task contexts (Supplementary Fig. 1c). The 1R model predicts that stimuli will shift flexibly along a shared axis—common “number line”, resulting in relatively small cross-context representational changes. In contrast, the 1D model predicts that stimuli will be represented on a different axis in each context, producing larger cross-context changes in geometry. Thus, examining representational shifts across contexts is essential for distinguishing between the two models. When comparing faces within the same context, 1D and 1R are similar because both represent ranks of faces within the same task dimension. However, 1D and 1R differ when comparing faces across contexts: 1R represents 1-D distances between faces on a common “number line” that is shared across contexts; In contrast, 1D represents 2-D distances between faces located in two distinct, orthogonal 1-D axes that vary across contexts. For example, a face ranked 2 in competence and 3 in popularity would be represented as (3,0) and (2,0) across the two contexts under the 1R model, but as (3,0) and (0,2) under the 1D model, because 1R assumes a shared rank axis whereas 1D assumes context-specific rank axes (Fig.2c).

The 1*I* estimates the extent to which neural activity patterns represent task-irrelevant information in the orthogonal representation. This further allows us to address two questions: (1) Do activity patterns representing context-dependent 1-D orthogonal representations fully suppress task-irrelevant information, or do they compress task-irrelevant distances within a 2-D structure while representing both task relevant and irrelevant information? (2) Is the degree of warping in the task-independent 2-D representation associated with different levels of suppression across individuals?

The *Context, Group*, and 1*I* RDMs served as control RDMs of non-interest. To resolve issues arising from multicollinearity between explanatory variables, we employed partial correlation to regress out the shared variance explained by other models. To identify brain areas containing the 2-D context-invariant and 1-D context-dependent orthogonal representations, we mapped back the *β*_*2D*_,*β*_*1D*,_*β*_*1R*_, *β*_*1I*_, and *β*_*1*I_ the extent to which the brain RDM of the activity patterns of each region of interest (ROI) was explained by the pairwise Euclidean distances in the model RDMs on the central voxel of each of the searchlight ROI across the whole brain to create continuous maps per subject. We smoothed the images using an 8-mm full-width at half maximum (FWHM) Gaussian kernel. We further performed one-sample t-tests for statistical inference in group-level analyses.

We conducted additional analyses to rigorously test competing hypotheses about representational structure. First, to compare our proposed task-dependent compressed orthogonal representation (1*D*) versus one involving solely the task-relevant dimension (1*R*), we examined how well each model explained the pattern of similarity between brain regions, especially for comparisons across different task contexts where 1*D* and 1*R* made divergent predictions. We estimated the average fit of each model (β values) within an independently-defined anatomical ROIs and used paired t-test to compare the model fits at the group level.

Secondly, we examined whether a stronger representation of task-irrelevant information correlated with increased warping in the subjective representation of social hierarchy, which was quantified by *ω*_*Behavioral*_ as described later in the “Estimating Representational Warping” section. To investigate this, we assessed the extent to which the activity patterns represented task-irrelevant information (*β*_1*I*_ in Eq.1) in the brain areas where the activity patterns represented the task-relevant information (*β*_*1R*_ in Eq.1), exhibiting the hypothesized 1-D task-dependent orthogonal structure. The region of interest (ROI) was defined using a threshold of pTFCE<0.05. This analysis aimed to shed light on the connection between subjective representation and the ability to suppress task-irrelevant distances, thus facilitating flexible decision making.

### Warping in representations of the cognitive map

#### The regression analyses to test the effects of warping on reaction times (RTs)

Using the reaction times for each trial (*RT*_*t*_) as an index of choice difficulty, we conducted several multiple regression analyses to examine the impact of potential warped representation on demand of cognitive control in inferential tasks. Each model used a different measure of distance between faces. These distance measures could not be included in a single model due to multicollinearity. For the behavioral analyses, we only included trials in which the participant made correct inferences among the congruent and incongruent trials. The number of congruent trials equaled the number of incongruent trials in the task (60 trials respectively).

We chose to analyze RTs over accuracy due to the limited trial-by-trial variability of accuracy in carefully trained participants. Specifically, all participants underwent extensive behavioral training to ensure they exceeded a predefined performance threshold before participating in the fMRI experiment.

In the first model, using the reaction times of each trial (*RT*^*t*^) as an index of choice difficulty, we tested for effects of context congruence (*C*_*t*_; 1 for congruent trials; -1 for incongruent trials; 0 for neutral trials), while controlling for the effects of the distances of the inferred trajectory, using multiple linear regression (Eq.2). The inferred trajectory of each inference trial was defined by a unique hub face enabling the transitive inferences of the relationship between F1 and F2. In addition to *C*_*t*_, the regression included the rank difference in the given task-relevant dimension (*D*^*t*^), and the Euclidean distance between F1 and F2 (*E*^*t*^), the task switch (*S*_*t*_, *S*_*t*_ was 1 when the current task-relevant context was different from that of the last trial, otherwise 0), and the subject-specific random effects (*β*^*0*^). Using the one-sample t-test, we tested the effects of congruence (the mean regression coefficient *β*_*C*_) on the RT.

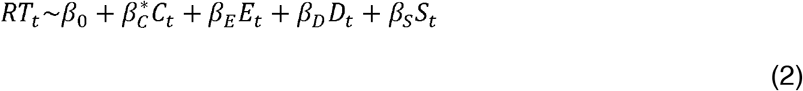

where *β*\* indicate the effect of interest to test the warping in the representation.

In the second model, we substituted context congruence (*C*^*t*^) in the first model with the absolute value of the cosine angle (*A*_*t*_) between two faces deviating from π/4 (the context congruency axis). *A*_*t*_ ranges from 1 when the inference angle is aligned with the congruent axis (y=x) to 0 when the inference angle is aligned with the incongruent axis (y=-x).

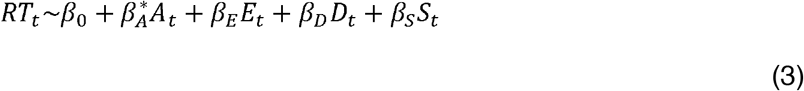

In the third model, we took the task irrelevant rank difference in the inferred trajectory into the model (*I*_*t*_). Specifically, to test if the effects of task irrelevant rank difference varied by

the model (*I*_*t*_). Specifically, to test if the effects of task irrelevant rank difference varied by context congruency, we included the interactions of rank difference and congruency (*C* ×*I*)^*t*^.

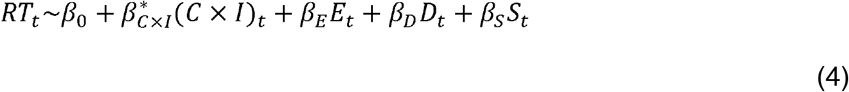

To test the influence of context congruency on the effects of task-irrelevant information, our fourth model separately estimated the effect of task-irrelevant rank differences for congruent and incongruent trials (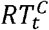 and 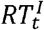 for congruent and incongruent trials respectively). Neutral trials were excluded from this analysis. Specifically, we tested if *β*_*I*_ had a greater negative value indicating faster reaction times in congruent compared to incongruent trials (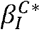 *vs*. 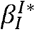).

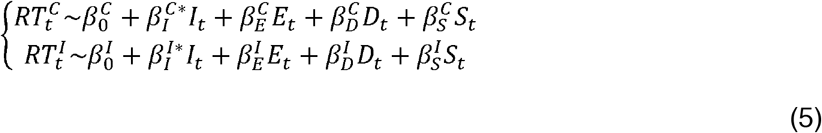

To assess model fit, we calculated and compared the Bayesian information criterion (BIC) based on the residual sum of squares (RSS) from each model’s RT predictions as below where *n* is the number of observations and *k* is the number of free parameters:

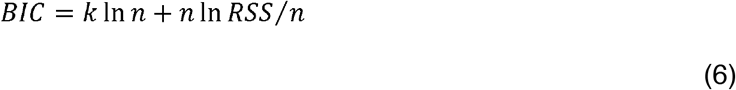

#### Estimating levels of hypothetical warping from reaction times

The previous finding using the same data showed that the RTs of inferences decrease with increases in the Euclidean distances of inferred trajectories ^11^. Given that, if the representation was not warped, the extent to which the Euclidean distances of the inferred trajectory between individuals explain the RTs of inferences (*β*_*E*_) should not be modulated by vector angles of the inferred trajectory (*θ*). On the other hand, if representations are warped, the vector angles of the inferred trajectory would account for variance in the extent to which Euclidean distances of the inferred trajectory explain the RTs in a manner that depended on whether the angles correspond to congruent or incongruent pairs. We estimated the relative levels of warping across participants by approximating the Euclidean distances between individual faces on the subjective representations from the RTs of the inferences.

To infer the subjective level of warping from RTs of inferences, first, we estimated the trial-by-trial variability in the degree to which the Euclidean distance of inferred trajectories explains the RT 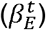 after accounting for the effects of other distance terms and the intercept, as Eq.7 shows below:

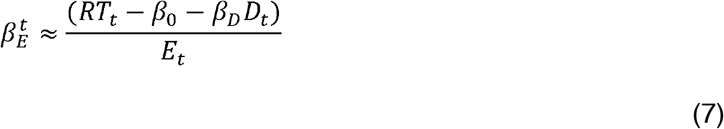

where *t* = {1,2 … *T*} and *T* is the number of inference trials, and the effects of rank differences in the task-relevant dimension on the RTs and *β*^*0*^ of each participant were estimated from a robust linear regression in which we inputted *E*_*t*_ and *D*_*t*_ to predict RTs. Importantly, the Eq.7 does not aim to regress out all the components predicting RTs as much as possible; instead, it selectively removes exogenous task-relevant variables (*D*_*t*_) while preserving potential endogenous variability that may arise from participants’ subjective internal representations. To minimize arbitrary assumptions, we adopted a deliberately conservative formulation, consistent with the prediction that task difficulty varies solely with *D*_*t*_ in an ideal decision maker.

Next, we inferred the norm vector distances 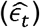 over the subjective representation from 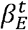. Since the longer the distances, the faster the RTs (while 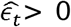 and 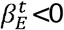 in general), we approximated 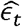 as follows (Eq.8):

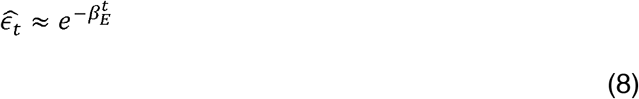

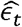 indicates the Euclidean distance of the unit vector of the inference trajectory of a certain vector angle (*θ*_*t*_). That is, if the representation was not warped, 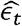 would have consistent values across the trials regardless of the vector angle, *θ*^*t*^. Consequently, the distribution of 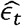 according to *θ*_*t*_ across trials should have a topology of the circle (Supplementary Fig.2). On the other hand, this distribution would have a topology of an ellipse that was elongated along the congruent axis (y=x) if the representation was warped. To test this, we examined the topology of the distribution of 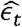 as a function of *θ*_*t*_ in polar coordinates. After excluding outlier trials of each participant (> mean 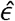 ± 3×standard deviation [SD]), if they exist, we converted the norm vector of every trial, 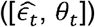 from polar-into cartesian-coordinates as follows (Eq.9):

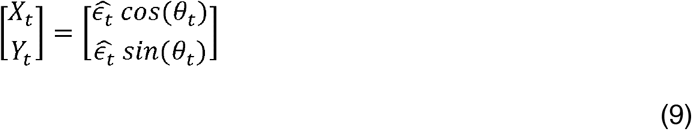

From the distribution of each participant, we measured the covariance between *X* and *Y* (*cov*(*X,Y*) to examine the relationships between two variables composing the distribution. The covariance indicated the direction of the relationship between two variables. That is, a positive covariance indicated that the distribution is elongated along the congruent axis (y=x) while a negative covariance indicated that the distribution is elongated along the incongruent axis (y=-x). Thus, to infer the geometry of the subjective representation of each participant, we computed the covariance matrix (*Σ*) as below (Eq.10):

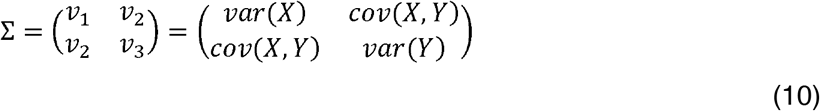

*Σ* allowed us to capture the subjective representation from the distribution of the RTs as follows. First, *cov*(*X,Y*) = 0 if the representations were not warped while *cov*(*X,Y*) >0 if the representations were warped along the congruent axis (Supplementary Fig.2) (c.f., *cov*(*X,Y*)<0 if the representations are warped along the incongruent axis, y=-x). Second, the ratio between and indicates to what extent RTs were explained better by the rank differences in one dimension compared to those in the other dimension. That is, if *var*(*X*) = *var* (*Y*), the participant represents the competence and popularity dimensions similarly, and the distances between faces in different ranks are the same in both dimensions (otherwise the representation is elongated along the vertical or horizontal axis). By applying *Σ* of each participant to the positions of faces in the actual 4×4 social hierarchy 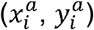, we estimated the new positions of individuals 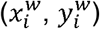 and the Euclidean distances 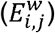 between faces (*i* and *j*) in the subjective representations of the social hierarchy (*Eq*.*11*).

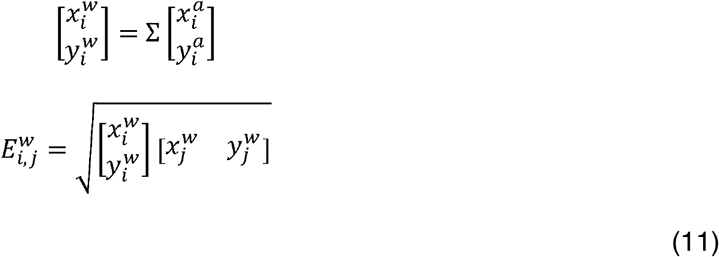

We estimated the level of warping as the ratio (*ω*^*Behvioral*^) between the length of the congruent unit vector, 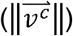 and the length of the incongruent unit vector 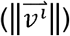 (Eq.12).

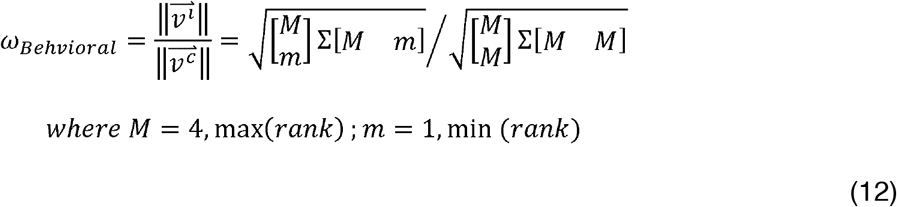

That is, the participant whose representation was more warped would have smaller *ω*_*Behvioral*_. It is important to note that we assumed that the RTs of inferences have an inverse logarithmic relationship with the distances between faces for convenience’s sake (Eq.8). Given that, *ω*_*Behvioral*_ should not be interpreted as the absolute level of warping in the subjective representation. Instead, *ω*^*Behvioral*^ was used for testing for the relative level of warping and ranking participants accordingly.

Note that *ω*_*Behvioral*_ was not estimated directly from the RT residuals but its covariance across trials as a function of decision vector angles. This approach does not require modeling all components that contribute to RT variability; instead, it isolates the component that systematically alters the covariance of RT residuals across trials along specific decision-vector directions. Consequently, the approach is resilient to trial-level fluctuations and insensitive to uniform shifts in RT caused by factors such as decision thresholds or non-decision times. In other words, equivalent results would be obtained using alternative models that incorporate additional RT components, as these factors do not influence the estimated level of representational warping.

To test if the representation was warped significantly along the congruent axis, *ω*^*Behvioral*^ was compared to the baseline (*ω*_*Base*_). This was estimated from each participant by randomly shuffling the relationship between the 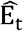 and *θ*_*t*_ (1000 times per subject). The difference between the levels of warping (*mean*[*ω*^*Base*^] − *ω*_*Behvioral*_) were inputted into the one-sample t test.

To validate this method, we performed a simulation. To test if we successfully infer the relative rank of warping (*ω*) across participants from the RTs using the above-mentioned approach, we generated synthetic reaction time data 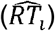 according to Eq.13:

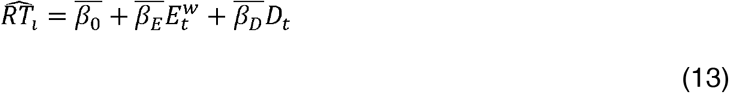

where 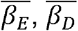, and the noise 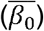 were sampled from normal distributions (ℕ) using the mean and and standard error of those estimated from participants’ data (Eq.2) as follows: 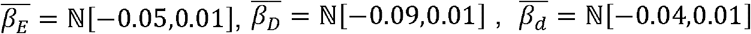 and 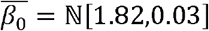. The Euclidean distances between individuals over the warped representations (*E*^*w*^) were estimated from the space to which we applied different levels of warping 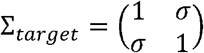 while varying *σ* ranging from 0 to 0.5 (the greater *σ*, the greater the warping) according to *Eq*.*7*. We generated 1000 sets of synthetic RT data. From the synthetic data, we estimated the level of warping 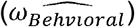 in same way (Eq.6-11). We further tested for the monotonic increase of the 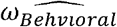 estimated from synthetic RTs as a function of the level of warping (*ω* ^*Behvioral*^) estimated from the actual structure (*Σ* ^*target*^).

The subjective warping estimate should not depend critically on the specific model used to estimate trial-wise RT residuals. In the present task, decision values were balanced across four mirrored directions in the representational space, such that each decision direction had corresponding horizontally mirrored, vertically mirrored, and diagonally opposite trials. This four-fold symmetry ensured that decision values were not biased toward a particular axis. Therefore, applying the same linear or nonlinear mapping from decision value to RT across all directions may change the overall scale of RT predictions, but should not selectively distort one direction of the representational geometry over another. To test the robustness of this assumption, we additionally estimated RT residuals using a DDM-based model and recomputed the warping estimates.

We modeled binary choices and reaction times using a drift diffusion model (DDM), in which noisy evidence accumulates over time until reaching one of two decision boundaries. Trial-wise predicted RTs were computed as shown in Eq.14, where *T*_*er*_ is non-decision time and *a* is the decision boundary. For each trial *i*, the drift rate (v_*i*_) was modeled as a linear function of the signed task-relevant decision variable (*R*_*i*_) and the signed task-irrelevant distractor (*D*_*i*_):

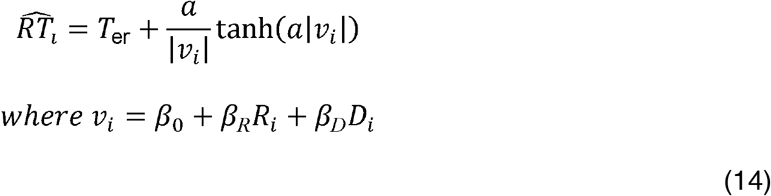

*β*_*R*_ and *β*_*D*_ capture their effects on drift rate, and *β*_*0*_ captures an overall drift bias. Under this model, congruent distractors increase the absolute drift rate and are expected to speed responses, whereas incongruent distractors reduce the absolute drift rate and are expected to slow responses.

#### Neural encoding of inferred trajectories over warped representations of the cognitive map

We tested if the brain activity is better accounted for by the distances between individuals over the subjective representations (*E*^*w*^) rather than the Euclidean distances over the actual 4×4 social hierarchy (*E*). For the univariate analysis images were smoothed using an 8-mm full-width at half maximum Gaussian kernel ^82^. To test this, we modeled the brain activity in the predefined regions of interest (ROIs) using a GLM. To deal with the inherent high correlation between *E*^*w*^ and *E*, we inputted *E* and *E*^*w*^-as parametric regressors into the GLM using a 2s boxcar function. It is noteworthy that we computed *E*^*w*^ by applying *ω* ^*Behavioral*^ to *E*. Though neither *E*^*w*^ nor *ω* ^*Behavioral*^ vary across trials dependent on RTs, we included trial-by-trial RTs as a non-interest regressor to ensure that any observed effects were not driven solely by RTs. In addition, the 6 motion regressors and the button press onsets were also included as regressors of no interest using a stick function. This allowed us to test for the effect of *E*^*w*^ in a parsimonious manner while controlling for the effects of *E* and RTs.

Additionally, to directly test the alternative hypothesis that brain activity might be explained by task switching, rather than the inferred trajectories, we tested additional GLMs. In these models, task switch (*S*_*t*_) was defined as trials when the context changed from one trial to the next at the time of the decision (F2), and included as an additional regressor, instead of *E*^*w*^ − *E*. Statistical inference was made within *a priori* selected ROIs that combined the anatomically defined vmPFC and EC (see ROI definitions).

#### Testing warping in the geometry of neural representations of the social hierarchy

We performed multivariate RSA across the whole brain to directly test the warping from the context-invariant neural representations of the social hierarchy. Unlike the analysis methods using pairwise distances to test different representational geometries, the rank correlation between the brain and model RDMs cannot prove the warping because the pairwise Euclidean distances in a warped representation highly correlate with the pairwise Euclidean distances in the actual 4×4 social hierarchy. Given that, to compute the level of warping from the neural representation, we compared the pattern dissimilarity between representations of the congruent and incongruent pairs. Specifically, first, we computed the average pattern dissimilarity across the pairs that created the same vector 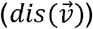 (Supplementary Fig.3b). Second, we computed the ratio between the average pattern dissimilarity of congruent vectors, 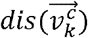 and that of equidistant incongruent vectors, 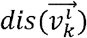. The 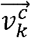 and 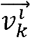 were symmetric. That is, 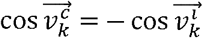 (vector angles) and 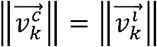 (vector distances). Third, the ratio between mean pattern dissimilarities was measured per pair of the congruent and incongruent vectors (9 vector pairs, *k*={1,2, …, 9}; See Supplementary Fig.3). Last, we estimated the level of warping (*ω*_*Neural*_) from the mean ratio of pattern dissimilarities between these symmetric vectors (Eq.15) per searchlight ROI across whole brain. Therefore, consistent with *ω*_*Behvioral*_, smaller *ω*_*Neural*_ indicated greater warping in the representations. Notably, *ω*_*Neural*_ was only sensitive to the shear transformation (warping) of a context-invariant representation but indifferent to other affine transformations (Supplementary Fig.3).

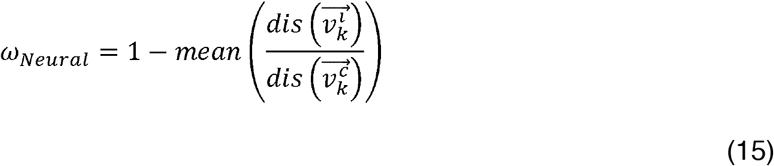

We created a continuous map of *ω*_*Neural*_ in the whole brain per subject by mapping back its value on the central voxel of the corresponding ROI. Each image was smoothed using an 8-mm FWHM Gaussian kernel and inputted into the group-level analysis using the one-sample t-test.

We performed a control analysis while excluding the longest vector pair (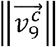 and 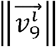) which was created by the representations of the 4 faces whose ranks were at the highest or at the lowest rank in either dimension. Notably, all representations of the remaining 12 faces served to create equidistant congruent and incongruent vectors. That is, the pattern dissimilarities (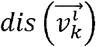 and 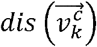) were estimated from different pairs of the same representations. This allowed us to test for warping while excluding the possibility that this effect was driven by a few extreme representations. Moreover, this careful control analysis was instrumental in addressing the potential influence of varying angle distributions resulting from differences in neighboring faces across grid locations. The statistical inference was made within an a priori ROI defined in the anatomically defined right HC (see ROI definitions).

We further tested for brain areas whose level of warping estimated from neural representations (*ω*_*Neural*_) correlates with those estimated from RT (*ω*_*Behvioral*_) across participants by inputting *ω*^*Behvioral*^ (demeaned) as a covariate for the second-level analysis.

#### Intersubject representational similarity analysis

We investigated individual differences in neural representations of the social hierarchy structure reflecting differences in behaviorally estimated levels of warping (*ω*_*Behvioral*_) across participants using the intersubject representational similarity analysis (IS-RSA) ^41–43^. The IS-RSA allowed us to examine the relationship between the neural representations and individual differences in cognitive control (RT biases) without presuming any representational geometric form. To perform this, we first constructed a warping dissimilarity matrix using the difference between the level of warping (*ω*_*Behvioral*_) across participants. We then build the brain RDMs for IS-RSA: We computed the RDMs by comparing the neural activity patterns of faces in two different social hierarchy dimensions estimated from different trials across blocks using Euclidean distances. We computed intersubject neural dissimilarity matrix (size=n×n, where *n*=27 subjects) by comparing brain RDMs estimated from the same searchlight ROI across participants from 1-Pearson correlation coefficients. We computed the intersubject neural similarity per searchlight ROI across the whole brain using the Kendall’s rank correlation between the intersubject neural similarity matrix and the behaviorally estimated warping similarity matrix (Supplementary Fig.4).

The statistical significance for IS-RSA was assessed against the baseline estimated from a permutation test. We generated 1000 similarity matrices while randomly shuffling *ω*_*Behvioral*_ across participants. We then created a null distribution of 1000 correlations between the shuffled matrix and the neural RDM of each searchlight ROI. From the normal distribution defined by the mean and standard deviation of a null distribution, we calculated p-values of each voxel across the whole brain. We corrected for multiple comparisons using the False Discovery Rate (FDR) threshold at q=0.05 across the whole brain ^83^.

In addition, to rule out the possibility that the effects of IS-RSA were driven by any single outlier subject, we performed the IS-RSA for n-1 subjects for n times (*n*=27) while excluding one subject at a time (leave-one-out analysis).

### Reactivation of current task-relevant context at the time of inferences

We tested the hypothesis that the participants who effectively use the task context that was relevant to the current goals to select appropriate actions in incongruent trials were more likely to build and have a more accurate social hierarchy representation, while those who were less likely to resolve incongruence might tend to have warped representations and their inferences would be modulated by the context congruence.

To test for the effect of context utilization on different levels of warping (*ω*_*Behvioral*_) in the representations, we performed the following behavioral and fMRI analyses. First, we tested if the accuracy biases (the tendency to make more accurate inferences on congruent than incongruent trials) related to the effects of congruence on the RTs (*β*_*C*_ in Eq.2). To capture greater variance across individuals, we test this in behavioral data acquired from a larger cohort of participants (*n*=218) (the flexibility test collected on day 2 training).

Next, using fMRI, we examined if participants tend to have greater reinstatement of the current context information at the time of inferences on incongruent than congruent trials. To address this, we performed a series of RSA analyses as follows: (1) We identified brain areas containing task-specific representations by comparing activity patterns at the time of the context cue presentation (See ROI definitions; Supplementary Fig.7a). (2) We estimated the mean activity patterns of each searchlight ROI across the whole brain at the time of inferences, using a boxcar function spanning from the onset of F2 to the reaction time, and modeled them according to the behavioral context (competence or popularity), congruence (congruence or incongruence), and button press (right and left) (8 activity patterns per ROI). The activity patterns associated with (1) and (2) were estimated from a single GLM in which we also included the motion regressors and other regressors of non-interest (stimulus onsets other than F2 and trials with missing responses); (3) We compared the activity patterns associated with context information (acquired at the time of the context cue) to the activity associated with inferences (acquired at the time of decision making) (Supplementary Fig.7c). We created a 10×8 RDM per searchlight ROI by comparing the activity patterns estimated from two independent blocks; (4) We created a model RDM based on a geometrical model (Supplementary Fig.7b) that comprised of independent vectors representing the representations of two context dimensions and those from inferences on the congruent and incongruent relationships, respectively. Based on this model, we created a model RDM in which the dissimilarity between representations was estimated from 1-cosine angle between two vectors (1 − *S*_*C*_). (5) We used a GLM to examine to what extent the model RDM explains variance of the neural RDM across the whole brain. In addition to the model RDM of interest, we included 3 other RDMs into the GLM to control for the potential effects of the different contexts, incongruence, and the different button presses (Supplementary Fig.7c); (6) We created continuous maps capturing the extent to which each model RDM accounted for the neural RDM per subject by mapping back them into a central voxel of each ROI. We smoothed the map of each subject using an 8-mm FWHM Gaussian kernel, and performed one-sample t tests for a group-level analysis; (7) To identify brain areas whose activity patterns associated with context coding was preferentially reinstated during incongruent compared to congruent trials, statistical inference was performed with TFCE within the brain areas having distinctive context representations identified from (1) (See ROI definitions).

### Statistical inferences

We perform group-level inference on whole-brain analyses unless we specified the hypothesis-driven *a priori* ROIs. We reported the results corrected for family-wise error (FWE) for multiple comparisons across the whole brain using threshold-free cluster enhancement (TFCE) ^84^ with 1000 iterations (pTFCE < 0.05). For the ROI analysis, the multiple comparisons were corrected within each of the ROIs using TFCE with 1000 iterations (pTFCE < 0.05). All statistical parametric maps presented in the manuscript are unmasked at the threshold, pTFCE < 0.05.

### ROI definitions

In the current study, we used 3 types of *a priori* ROIs: (1) the searchlight ROIs across the whole brain for multivariate analyses, (2) the anatomical ROIs defined independently from the current task based on a probabilistic map acquired from independent studies, and (3) the functional ROIs based on the results of independent time points of the current task. First, we defined the searchlight ROIs as a sphere including 100 voxels around each of the central voxels across the whole brain. Second, the anatomical ROIs were defined bilaterally in the HC combining only its anterior and middle portions ^85^, EC ^86,87^, amygdala ^88^, and vmPFC/mOFC ^89^. Third, two types of functional ROIs were defined through independent analyses. The first analysis aimed to identify brain areas containing task-relevant information using an orthogonal representation with compression (1*D*). The second analysis focused on identifying brain areas with distinctive context representations at the time of the context cue representation. To define the first set of functional ROIs, we selected the cluster that included the dmPFC/ pre-SMA, dmPFC, and lPFC based on the results of the RSA shown in Fig.3c. For the second set of functional ROIs, a whole-brain RSA was performed by comparing the mean activity patterns during the presentation of context cues across different blocks. Supplementary Fig. 7a described the detailed procedure. The cluster forming threshold for these functional ROIs was set at p_TFCE_<0.05, corrected for multiple comparisons across the entire brain.

### Neural network model

The recurrent neural network (RNN) model was trained on the same task used in the fMRI experiment. On each trial, a cue indicating the relevant social hierarchy dimension (context cue) and a pair of images of faces (F1 and F2) were presented sequentially to the network, which was trained to predict which of the two faces corresponded to the individual that ranked higher on the task-relevant dimension (Fig.1b). The training and test sets for the model consisted of the same sets of trial samples used to train and test the participants in the human fMRI experiment: the model was trained only on limited pairs whose ranks differed by one in the given social hierarchy dimension, but was tested on pairs whose ranks differed by one or more than one - thus requiring transitive inference. Consistent with the human participants, the RNN model did not receive any explicit indication of these groupings, nor of the fact that the faces were arranged in a latent 4×4 grid structure along the two social hierarchy dimensions.

#### Model architecture

The RNN model was built from standard building blocks including a convolutional neural network (CNN) for processing the images of faces and a Long Short-Term Memory (LSTM) network for processing the sequence of stimuli on each trial (Cue, F1, and F2). A recurrent network was chosen in order to better replicate the experience of the participants, who received the stimuli sequentially, and to capture the emergent dynamics of representations unfolding over the course of each trial. The Cue was represented as a one-hot vector (*x*_*C*_ in Eq.16 and Fig.1b) indicating whether the relevant context/dimension was popularity or competence, and was embedded in a 32-dimensional vector (*e*_*C*_ using a linear layer (*W*_*C*_ . We used the same face images with the fMRI experiment to train the neural network model. The two faces (F1 and F2) were given to the network as 64×64 grayscale images (*x*_1_ and *x*_2_), which were embedded in two 32-dimensional vectors (*e*^*1*^ and *e*^*2*^ in Eq.16 and Fig.1b) using the same CNN:

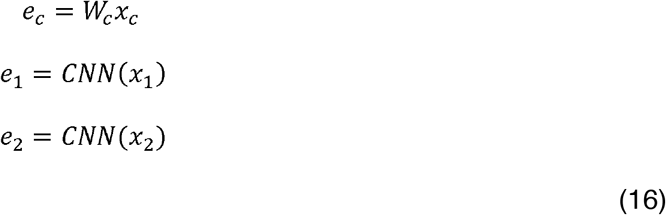

The LSTM then processed these three embeddings in sequence (*e*_*c*_ → *e*_*1*_ → *e*_*2*_):

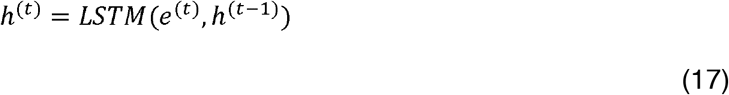

where *h*^(*t)*^ is the hidden state of the LSTM at time step (the sequential processing steps of the RNN) *t* and *e*^*t*^ is the embedding (e_c_, e_1_, or e_2_) at time step *t* (Eq.17). The model produced a prediction 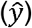 for whether the first or second face ranked higher along the relevant dimension using a linear output layer (*W*^*o*^) applied to the final hidden state (Eq.18):

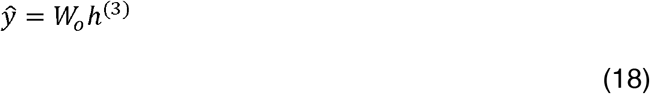

#### Training procedure

The RNN was trained with standard procedures, including a cross-entropy loss function and the backpropagation algorithm. The model was trained in a supervised manner on batches of 32 trials randomly sampled from the entire training set. Unlike the human participants, training the neural network on one dimension at a time in a blocked manner would have caused the network to forget during the second block what had been learned in the first (well-known problem of catastrophic forgetting ^38,39^. The catastrophic forgetting phenomenon represents a relatively well-studied discrepancy between the learning behavior of neural networks as compared with humans. To avoid catastrophic forgetting, the training set for the RNN was randomly sampled from batches combining all pairs presented during learning blocks thoughout 3 days of behavioral training of human participants. Therefore, the task relevant dimensions were randomly intermixed across trials. While this is not central to the issues under investigation in the current study, other studies ^90,91^ including ours ^92^ are exploring the issue of catastrophic forgetting.

#### Implementation details

All neural network simulations were implemented using PyTorch ^93^. The model was optimized with Adam ^94^ with a learning rate of 0.001 and a batch size of 32 for 1000 gradient steps. For each simulation, 20 runs were performed with different random initializations. All parameters were initialized to be small (i.e., in the “rich” regime) using the PyTorch default option. The CNN included two convolutional layers with kernel sizes (3, 3), strides (2, 2) and number of channels (4, 8). Max pooling followed each convolutional layer with kernel sizes (2, 2) and strides (2, 2). The CNN also included a final linear layer to map the output of the last pooling operation to a single vector with 32 dimensions. The hidden layer of the LSTM was 128 dimensions.

### Representational geometry in the RNN

To test our first two hypotheses about the nature of the representations learned by the model, we conducted an RSA analogous to the one used with the fMRI data. Averaged representations for each face in each context were collected from the activity patterns of the RNN hidden layer (*h*), and an RDM was computed using pairwise Euclidean distances. This created a 32×32 RDM, by including the patterns associated with 16 faces in two social hierarchy dimensions, considering that the test trials in the RNN were randomly sampled from the pairs presented in either test blocks in the behavioral training or in the fMRI. The representational geometries of the hidden layers of neural network model was tested using the same GLM with the fMRI experiment (Eq.1) while excluding the group RDM since the pairs were intermixed during training across learning groups in RNN. The GLM was performed after every 50 training steps, allowing us to visualize the emergence of these phenomena throughout the course of training. We further performed one-sample t-tests for group-level analysis across 20 runs.

#### Testing warping in the RNN representations

To test the hypothesis that the map-like representations learned by the model were warped along the congruent diagonal, we measured the ratio (Eq.19; analogous to Eq.8 in behavioral data of human participants) of average Euclidean distances between RNN representations of incongruent pairs of faces 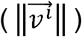 to the average distances between representations of congruent pairs 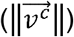. We also conducted a two-sample t-test measuring whether the average distances between congruent pairs of faces were significantly larger than the average distances between incongruent pairs of faces. This analysis was also conducted every 50 training steps.

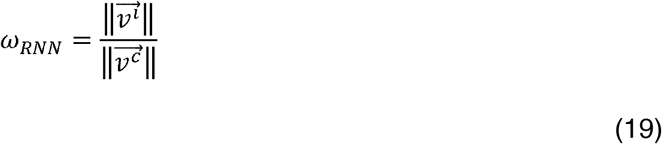

#### Ablation simulation: testing the effects of context information on representational geometry

To better understand the warping, we observed in both the human brain and the neural network model, we performed a simulated “ablation” on the model where we degraded the incoming information from the context cue indicating the current task-relevant social hierarchy dimension (e_c_ in Fig.1b and Eq.20). This was achieved by multiplying the cue’s embedding (e_c_) by a weight (*α*^-1^) ranging from 0 to 1.

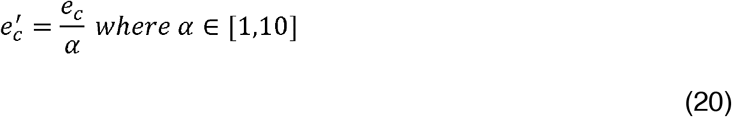

We subsequently measured the warping in the model’s representations after learning using the same method described above (Eq.19). In addition, we performed a linear regression analysis (Eq.21) testing whether there was a significant effect of the ablation weight (*α*^-1^) on *ω*_*RNN*_.

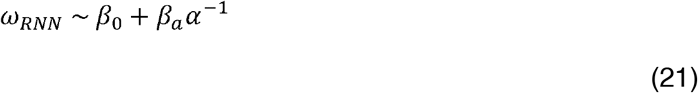

#### Relationship between congruence, warping, and accuracy

To test the hypothesis that the model learned to perform well at congruent trials before it learned to perform well at the incongruent trials, we conducted a logistic regression after every 50 training steps to test whether congruence significantly predicted the task performance accuracy.

We also tested whether the difference between model’s performance on congruent (accuracy^+^) and incongruent trials (accuracy^-^) could be predicted by the degree of warping in the model’s representations. To do this, we performed a linear regression predicting the accuracy ratio at each time step from (1) levels of warping, (2) a linear time variable, and (3) their interaction (Eq.22). Here, we included the linear time predictor (T=[0,20], every 50 training steps) in addition to the warping predictor, as well as their interaction to capture the possibility that the effects of warping could vary over the course of training.

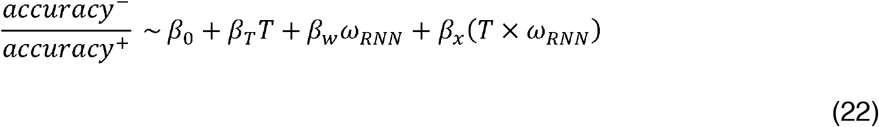

## Data Availability

The behavioral data and unthresholded neuroimaging statistical maps supporting the findings of this study are publicly available on the Open Science Framework (https://osf.io/ra45p; DOI: 10.17605/OSF.IO/RA45P).

## Code Availability

The code used to analyze the data is available on the Open Science Framework (https://osf.io/ra45p; DOI: 10.17605/OSF.IO/RA45P). The code of the neural network model is available on Code Ocean (https://doi.org/10.24433/CO.7457120.v1). The code associated with the behavioral training protocols is available on the Open Science Framework (https://osf.io/bnc3w; DOI: 10.17605/OSF.IO/BNC3W).

## Author Contributions

SAP and EDB conceived the study and designed the experimental paradigm. SAP and DSM programmed the behavioral task and collected the behavioral and fMRI data. SAP analyzed the behavioral and fMRI data. MZ, JR, and RCO designed, implemented, trained, and analyzed the recurrent neural network models. SAP and EDB contributed to the interpretation of the modeling and neuroimaging results. EDB supervised the project and acquired funding. SAP wrote the original draft of the manuscript with input from JR, and EDB. All authors discussed the results, reviewed and edited the manuscript, and approved the final version.

## References

1. Miller, E. K. & Cohen, J. D. An integrative theory of prefrontal cortex function. Annu. Rev. Neurosci. 24, 167–202 (2001).

2. Freund, M. C., Etzel, J. A. & Braver, T. S. Neural Coding of Cognitive Control: The Representational Similarity Analysis Approach. Trends Cogn. Sci. 25, 622–638 (2021).

3. Vaidya, A. R. & Badre, D. Abstract task representations for inference and control. Trends Cogn. Sci. 26, 484–498 (2022).

4. Musslick, S. & Cohen, J. D. Rationalizing constraints on the capacity for cognitive control. Trends Cogn. Sci. 25, 757–775 (2021).

5. Bernardi, S. et al. The geometry of abstraction in the hippocampus and prefrontal cortex. Cell 183, 954–967.e21 (2020).

6. Koechlin, E. Prefrontal executive function and adaptive behavior in complex environments. Curr. Opin. Neurobiol. 37, 1–6 (2016).

7. Higo, T., Mars, R. B., Boorman, E. D., Buch, E. R. & Rushworth, M. F. S. Distributed and causal influence of frontal operculum in task control. Proceedings of the National Academy of Sciences 108, 4230–4235 (2011).

8. Takagi, Y., Hunt, L. T., Woolrich, M. W., Behrens, T. E. & Klein-Flügge, M. C. Adapting non-invasive human recordings along multiple task-axes shows unfolding of spontaneous and over-trained choice. Elife 10, (2021).

9. Flesch, T., Juechems, K., Dumbalska, T., Saxe, A. & Summerfield, C. Orthogonal representations for robust context-dependent task performance in brains and neural networks. Neuron 110, 1258–1270.e11 (2022).

10. Nee, D. E. Integrative frontal-parietal dynamics supporting cognitive control. Elife 10, (2021).

11. Park, S. A., Miller, D. S., Nili, H., Ranganath, C. & Boorman, E. D. Map making: constructing, combining, and inferring on abstract cognitive maps. Neuron 107, 1226–1238.e8 (2020).

12. Park, S. A., Miller, D. S. & Boorman, E. D. Inferences on a multidimensional social hierarchy use a grid-like code. Nat. Neurosci. 24, 1292–1301 (2021).

13. Schuck, N. W., Cai, M. B., Wilson, R. C. & Niv, Y. Human Orbitofrontal Cortex Represents a Cognitive Map of State Space. Neuron 91, 1402–1412 (2016).

14. Garvert, M. M., Dolan, R. J. & Behrens, T. E. J. A map of abstract relational knowledge in the human hippocampal–entorhinal cortex. Elife 6, (2017).

15. Behrens, T. E. J. et al. What is a cognitive map? organizing knowledge for flexible behavior. Neuron 100, 490–509 (2018).

16. Bellmund, J. L. S., Gärdenfors, P., Moser, E. I. & Doeller, C. F. Navigating cognition: Spatial codes for human thinking. Science vol. 362 eaat6766 (2018).

17. Eichenbaum, H. & Cohen, N. J. Can We Reconcile the Declarative Memory and Spatial Navigation Views on Hippocampal Function? Neuron 83, 764–770 (2014).

18. Knudsen, E. B. & Wallis, J. D. Hippocampal neurons construct a map of an abstract value space. Cell 184, 4640–4650.e10 (2021).

19. Fusi, S., Miller, E. K. & Rigotti, M. Why neurons mix: High dimensionality for higher cognition. Curr. Opin. Neurobiol. 37, 66–74 (2016).

20. Buschman, T. J. Balancing Flexibility and Interference in Working Memory. Annu. Rev. Vis. Sci. 7, (2021).

21. Rigotti, M. et al. The importance of mixed selectivity in complex cognitive tasks. Nature 497, 585–590 (2013).

22. Badre, D., Bhandari, A., Keglovits, H. & Kikumoto, A. The dimensionality of neural representations for control. Curr. Opin. Behav. Sci. 38, 20–28 (2020).

23. Ehrlich, D. B. & Murray, J. D. Geometry of neural computation unifies working memory and planning. Proc. Natl. Acad. Sci. U. S. A. 119, e2115610119 (2022).

24. Garner, K. G. & Dux, P. E. Knowledge generalization and the costs of multitasking. Nat. Rev. Neurosci. 24, 98–112 (2023).

25. Mante, V., Sussillo, D., Shenoy, K. V. & Newsome, W. T. Context-dependent computation by recurrent dynamics in prefrontal cortex. Nature 503, 78–84 (2013).

26. Aoi, M. C., Mante, V. & Pillow, J. W. Prefrontal cortex exhibits multidimensional dynamic encoding during decision-making. Nat. Neurosci. 23, 1410–1420 (2020).

27. Bichot, N. P., Xu, R., Ghadooshahy, A., Williams, M. L. & Desimone, R. The role of prefrontal cortex in the control of feature attention in area V4. Nature Communications 2019 10:1 10, 1–12 (2019).

28. Ho, M. K. et al. People construct simplified mental representations to plan. Nature 606, 129–136 (2022).

29. Barsalou, L. W. Ad hoe categories. Mem. Cognit. 11, 211–227 (1983).

30. Mack, M. L., Love, B. C. & Preston, A. R. Dynamic updating of hippocampal object representations reflects new conceptual knowledge. Proc. Natl. Acad. Sci. U. S. A. 113, 13203–13208 (2016).

31. Nieh, E. H. et al. Geometry of abstract learned knowledge in the hippocampus. 80 | Nature | 595, (2021).

32. Tang, E. et al. Effective learning is accompanied by high-dimensional and efficient representations of neural activity. Nat. Neurosci. 22, 1000–1009 (2019).

33. MacDowell, C. J., Tafazoli, S. & Buschman, T. J. A Goldilocks theory of cognitive control: Balancing precision and efficiency with low-dimensional control states. Curr. Opin. Neurobiol. 76, 102606 (2022).

34. Rougier, N. P., Noelle, D. C., Braver, T. S., Cohen, J. D. & O’Reilly, R. C. Prefrontal cortex and flexible cognitive control: Rules without symbols. Proc. Natl. Acad. Sci. U. S. A. 102, 7338–7343 (2005).

35. Panichello, M. F. & Buschman, T. J. Shared mechanisms underlie the control of working memory and attention. Nature 592, 601–605 (2021).

36. Hochreiter, S. & Schmidhuber, J. Long Short-Term Memory. Neural Comput. 9, 1735–1780 (1997).

37. Park, S. A., Miller, D. S. & Boorman, E. D. Protocol for building a cognitive map of structural knowledge in humans by integrating abstract relationships from separate experiences. STAR Protoc. 2, 100423 (2021).

38. McClelland, J. L., McNaughton, B. L. & O’Reilly, R. C. Why there are complementary learning systems in the hippocampus and neocortex: Insights from the successes and failures of connectionist models of learning and memory. Psychol. Rev. 102, 419–457 (1995).

39. McCloskey, M. & Cohen, N. J. Catastrophic Interference in Connectionist Networks: The Sequential Learning Problem. Psychology of Learning and Motivation - Advances in Research and Theory 24, 109–165 (1989).

40. Egner, T. Congruency sequence effects and cognitive control. Cogn. Affect. Behav. Neurosci. 7, 380–390 (2007).

41. Cohen, J. D. et al. Computational approaches to fMRI analysis. Nature Neuroscience 2017 20:3 20, 304–313 (2017).

42. Hasson, U., Nir, Y., Levy, I., Fuhrmann, G. & Malach, R. Intersubject Synchronization of Cortical Activity During Natural Vision. Science (1979). 303, 1634–1640 (2004).

43. Chen, P. H. A., Jolly, E., Cheong, J. H. & Chang, L. J. Intersubject representational similarity analysis reveals individual variations in affective experience when watching erotic movies. Neuroimage 216, 116851 (2020).

44. Theves, S., Fernández, G. & Doeller, C. F. The Hippocampus Maps Concept Space, Not Feature Space. The Journal of Neuroscience 40, 7318–7325 (2020).

45. Barbas, H. & Blatt, G. J. Topographically specific hippocampal projections target functionally distinct prefrontal areas in the rhesus monkey. Hippocampus 5, 511–533 (1995).

46. Collins, A. G. E. & Frank, M. J. Cognitive control over learning: Creating, clustering, and generalizing task-set structure. Psychol. Rev. 120, 190–229 (2013).

47. Collins, A. G. E. & Frank, M. J. Motor Demands Constrain Cognitive Rule Structures. PLoS Comput. Biol. 12, e1004785 (2016).

48. Usher, M. & McClelland, J. L. The time course of perceptual choice: the leaky, competing accumulator model. Psychol. Rev. 108, 550–592 (2001).

49. Constantinescu, A. O., O’Reilly, J. X. & Behrens, T. E. J. Organizing conceptual knowledge in humans with a gridlike code. Science (1979). 352, 1464–1468 (2016).

50. Bao, X. et al. Grid-like Neural Representations Support Olfactory Navigation of a Two-Dimensional Odor Space. Neuron 102, 1066–1075.e5 (2019).

51. Bellmund, J. L. S., Deuker, L., Schröder, T. N. & Doeller, C. F. Grid-cell representations in mental simulation. Elife 5, (2016).

52. Doeller, C. F., Barry, C. & Burgess, N. Evidence for grid cells in a human memory network. Nature 463, 657–661 (2010).

53. Viganò, S., Rubino, V., Soccio, A. Di, Buiatti, M. & Piazza, M. Grid-like and distance codes for representing word meaning in the human brain. Neuroimage 232, 117876 (2021).

54. Theves, S., Fernandez, G. & Doeller, C. F. The Hippocampus Encodes Distances in Multidimensional Feature Space. Current Biology 29, 1226–1231.e3 (2019).

55. Nitsch, A., Garvert, M. M., Bellmund, J. L. S., Schuck, N. W. & Doeller, C. F. Grid-like entorhinal representation of an abstract value space during prospective decision making. Nature Communications 2024 15:1 15, 1198–(2024).

56. Muhle-Karbe, P. S. et al. Goal-seeking compresses neural codes for space in the human hippocampus and orbitofrontal cortex. Neuron 111, 3885–3899.e6 (2023).

57. Kumaran, D., Melo, H. L. & Duzel, E. The Emergence and Representation of Knowledge about Social and Nonsocial Hierarchies. Neuron 76, 653–666 (2012).

58. Kumaran, D., Banino, A., Blundell, C., Hassabis, D. & Dayan, P. Computations Underlying Social Hierarchy Learning: Distinct Neural Mechanisms for Updating and Representing Self-Relevant Information. Neuron 92, 1135–1147 (2016).

59. Bickart, K. C., Wright, C. I., Dautoff, R. J., Dickerson, B. C. & Barrett, L. F. Amygdala volume and social network size in humans. Nature Neuroscience 2010 14:2 14, 163–164 (2010).

60. Noonan, M. A. P. et al. A Neural Circuit Covarying with Social Hierarchy in Macaques. PLoS Biol. 12, e1001940 (2014).

61. Lewis, P. A., Rezaie, R., Brown, R., Roberts, N. & Dunbar, R. I. M. Ventromedial prefrontal volume predicts understanding of others and social network size. Neuroimage 57, 1624–1629 (2011).

62. Sallet, J. et al. Social network size affects neural circuits in macaques. Science (1979). 334, 697–700 (2011).

63. Testard, C. et al. Social connections predict brain structure in a multidimensional free-ranging primate society. Sci. Adv. 8, 5794 (2022).

64. Rudorf, S. & Hare, T. A. Interactions between Dorsolateral and Ventromedial Prefrontal Cortex Underlie Context-Dependent Stimulus Valuation in Goal-Directed Choice. The Journal of Neuroscience 34, 15988–15996 (2014).

65. Grueschow, M., Polania, R., Hare, T. A. & Ruff, C. C. Automatic versus Choice-Dependent Value Representations in the Human Brain. Neuron 85, 874–885 (2015).

66. Mahmoodi, A. et al. Causal role of a neural system for separating and selecting multidimensional social cognitive information. Neuron 111, 1152–1164.e6 (2023).

67. Moneta, N., Garvert, M. M., Heekeren, H. R. & Schuck, N. W. Task state representations in vmPFC mediate relevant and irrelevant value signals and their behavioral influence. Nat. Commun. 14, 3156 (2023).

68. Botvinick, M. M., Carter, C. S., Braver, T. S., Barch, D. M. & Cohen, J. D. Conflict monitoring and cognitive control. Psychol. Rev. 108, 624–652 (2001).

69. Braver, T. S., Reynolds, J. R. & Donaldson, D. I. Neural mechanisms of transient and sustained cognitive control during task switching. Neuron 39, 713–726 (2003).

70. Gillan, C. M., Kosinski, M., Whelan, R., Phelps, E. A. & Daw, N. D. Characterizing a psychiatric symptom dimension related to deficits in goal directed control. Elife 5, (2016).

71. Schafer, M. & Schiller, D. The Hippocampus and Social Impairment in Psychiatric Disorders. Cold Spring Harb. Symp. Quant. Biol. 83, 105–118 (2018).

72. Shields, C. N. & Gremel, C. M. Review of Orbitofrontal Cortex in Alcohol Dependence: A Disrupted Cognitive Map? Alcohol. Clin. Exp. Res. 44, 1952–1964 (2020).

73. Whittington, J. C. R., McCaffary, D., Bakermans, J. J. W. & Behrens, T. E. J. How to build a cognitive map. Nature Neuroscience 2022 25:10 25, 1257–1272 (2022).

74. Hassabis, D., Kumaran, D., Summerfield, C. & Botvinick, M. Neuroscience-Inspired Artificial Intelligence. Neuron 95, 245–258 (2017).

75. Saxe, A., Nelli, S. & Summerfield, C. If deep learning is the answer, what is the question? Nature Reviews Neuroscience 2020 22:1 22, 55–67 (2020).

76. Lake, B. M., Ullman, T. D., Tenenbaum, J. B. & Gershman, S. J. Building machines that learn and think like people. https://doi.org/10.1017/S0140525X16001837 (2017) doi:10.1017/S0140525X16001837.

77. Russin, J., O’reilly, R. C. & Bengio, Y. Deep learning needs a prefrontal cortex. In International Conference on Learning Representations (ICLR) (2020).

78. Kriegeskorte, N. Representational similarity analysis – connecting the branches of systems neuroscience. Front. Syst. Neurosci. 2, 4 (2008).

79. Nili, H. et al. A Toolbox for Representational Similarity Analysis. PLoS Comput. Biol. 10, e1003553 (2014).

80. Diedrichsen, J. & Kriegeskorte, N. Representational models: A common framework for understanding encoding, pattern-component, and representational-similarity analysis. PLoS Comput. Biol. 13, e1005508 (2017).

81. Friston, K. J., Diedrichsen, J., Holmes, E. & Zeidman, P. Variational representational similarity analysis. Neuroimage 201, 115986 (2019).

82. Mikl, M. et al. Effects of spatial smoothing on fMRI group inferences. Magn. Reson. Imaging 26, 490–503 (2008).

83. Benjamini, Y. & Hochberg, Y. On the Adaptive Control of the False Discovery Rate in Multiple Testing with Independent Statistics. Journal of Educational and Behavioral Statistics 25, 60 (2000).

84. Smith, S. M. & Nichols, T. E. Threshold-free cluster enhancement: Addressing problems of smoothing, threshold dependence and localisation in cluster inference. Neuroimage 44, 83–98 (2009).

85. Yushkevich, P. A. et al. Quantitative comparison of 21 protocols for labeling hippocampal subfields and parahippocampal subregions in in vivo MRI: Towards a harmonized segmentation protocol. Neuroimage 111, 526–541 (2015).

86. Amunts, K. et al. Cytoarchitectonic mapping of the human amygdala, hippocampal region and entorhinal cortex: Intersubject variability and probability maps. Anat. Embryol. (Berl). 210, 343–352 (2005).

87. Zilles, K. & Amunts, K. Centenary of Brodmann’s map conception and fate. Nat. Rev. Neurosci. 11, 139–145 (2010).

88. Tzourio-Mazoyer, N. et al. Automated anatomical labeling of activations in SPM using a macroscopic anatomical parcellation of the MNI MRI single-subject brain. Neuroimage 15, 273–289 (2002).

89. Neubert, F.-X., Mars, R. B., Sallet, J. & Rushworth, M. F. S. Connectivity reveals relationship of brain areas for reward-guided learning and decision making in human and monkey frontal cortex. Proc. Natl. Acad. Sci. U. S. A. 112, 1–10 (2015).

90. Flesch, T., Balaguer, J., Dekker, R., Nili, H. & Summerfield, C. Comparing continual task learning in minds and machines. Proc. Natl. Acad. Sci. U. S. A. 115, E10313–E10322 (2018).

91. Flesch, T., Nagy, D. G., Saxe, A. & Summerfield, C. Modelling continual learning in humans with Hebbian context gating and exponentially decaying task signals. https://doi.org/10.48550/arxiv.2203.11560 (2022) doi:10.48550/arxiv.2203.11560.

92. Russin, J., Zolfaghar, M., Park, S. A., Boorman, E. & O’Reilly, R. C. A neural network model of continual learning with cognitive control. Cognitive Science Society (CogSci 2022) https://doi.org/10.48550/arXiv.2202.04773 (2022) doi:10.48550/arXiv.2202.04773.

93. Paszke, A. et al. PyTorch: An Imperative Style, High-Performance Deep Learning Library. Adv. Neural Inf. Process. Syst. 32, (2019).

94. Kingma, D. P. & Ba, J. L. Adam: A Method for Stochastic Optimization. in 3rd International Conference on Learning Representations, ICLR 2015 (International Conference on Learning Representations, ICLR, 2014). doi:10.48550/arXiv.1412.6980.

