## Supplementary for "The representational geometry of cognitive maps under cognitive control"

Supplementary Figure S1
Supplementary Figure S2
Supplementary Figure S3
Supplementary Figure S4
Supplementary Figure S5
Supplementary Figure S6
Supplementary Figure S7
Supplementary Figure S8
Supplementary Figure S9

Supplementary Table 1
Supplementary Table 2
Supplementary Table 3
Supplementary Table 4
Supplementary Table 5
Supplementary Table 6
Supplementary Table 7
Supplementary Table 8

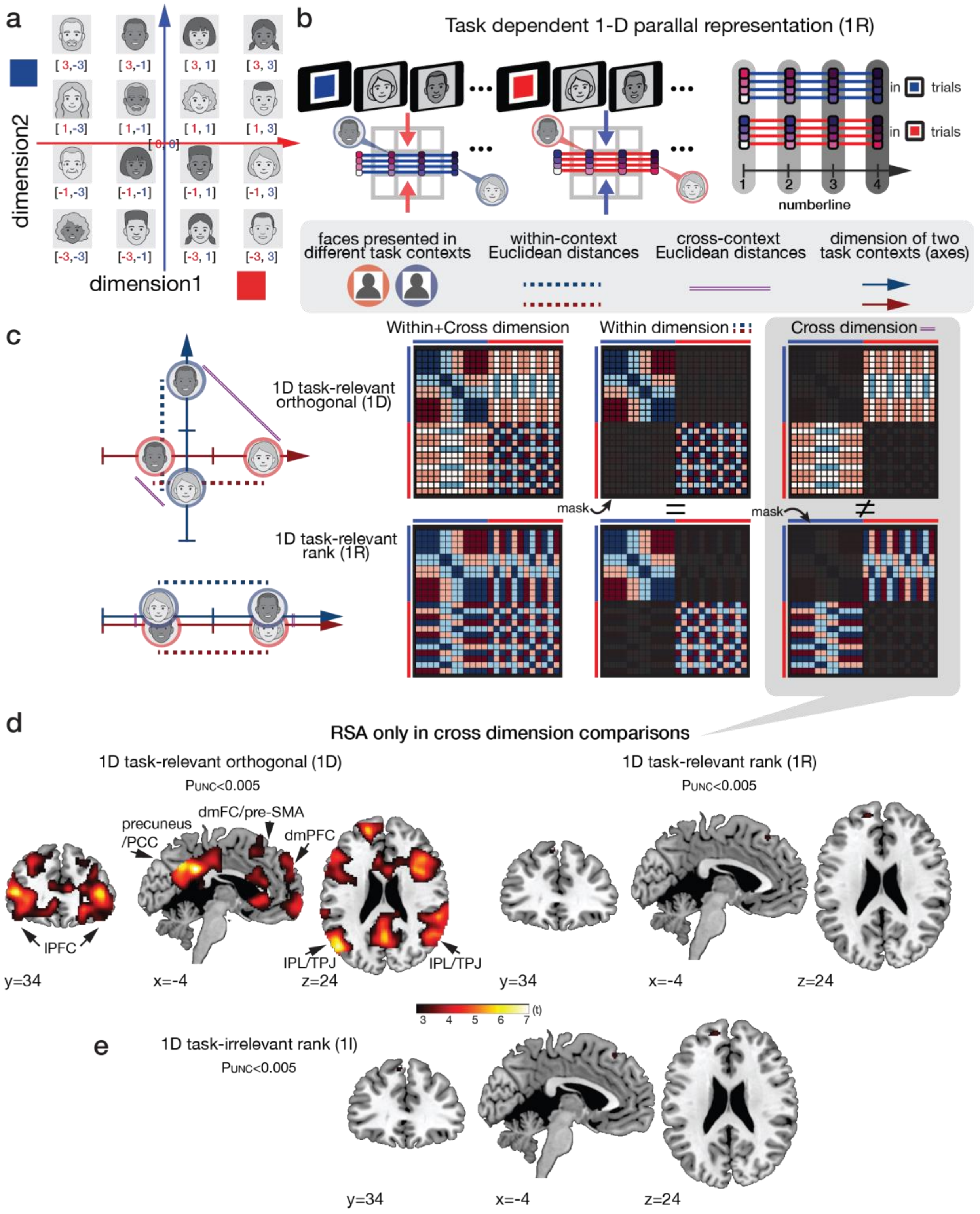


**Supplementary Figure 1. The multivariate representational similarity analysis (RSA) using a general linear model (GLM) to examine the representational geometry in the brain and RNN**

**a.** The coordinates of 16 faces that indicate their position in the context-invariant 2-D representation. It was used to compute pairwise Euclidean distances between faces and create the model RDMs. The coordinates of the task-irrelevant axis were set to zero when computing Euclidean distances between faces in the context-dependent 1D orthogonal representation.

**b.** Illustration of the one-dimensional (1-D) task-relevant representation on a shared axis (*1R* model). In *1R*, both task contexts are represented along a single shared axis. Representations of each face preserve their relative ordering within each context, but because both contexts map onto the same axis, cross-context Euclidean distances differ from those predicted by the orthogonal 1-D model. Within-context distances remain identical between the two models. *1R* is an alternative form of 1-D representation that is geometrically distinct from the orthogonal 1-D model (*1D* in **Fig. 2c**). Both are 1-D representations of task-relevant information, but they differ in how the two task dimensions are arranged in representational space. To determine which model better explains the variability in representation dissimilarity, we included both regressors jointly in the same GLM during the RSA analysis (**Fig. 3a**).

**c.** Distinguishing Orthogonal 1-D (1D) and Shared-Axis 1-D (1R) Models via pairwise distances in a representational space. Comparison of model-predicted representational dissimilarity matrices (RDMs) for the orthogonal 1-D (1D) and shared-axis 1-D (1R) models. Within-context distances (dotted regions) are identical for both models and thus do not provide discriminative power. In contrast, cross-context distances (solid regions) differ systematically because the 1D model represents contexts on orthogonal axes whereas the 1R model maps both contexts onto a single shared dimension. This shows that proper model comparisons should focus on cross-context distances. Given this, we conducted direct model comparisons using an additional RSA to evaluate the ability of the 1D and 1R models to explain multivariate activity patterns, particularly in cross-dimension comparisons where they diverge most, while masking out within-context structure.

**d.** Direct model comparisons between the 1-D orthogonal representation with compression ($1D$) and the model representing only the relevant dimension without compression ($1R$). We conducted RSA to assess the abilities of the 1D and 1R models to explain multivariate activity patterns, particularly in cross-dimension comparisons where they exhibit the most divergence. The cross-dimension RSA results show that this effect indeed driven by the context-dependent 1-D orthogonal representation, even when considering only cross-context pattern dissimilarity and not within-context pattern dissimilarity, where distinguishing between the $1D$ and $1R$ models becomes challenging (left panel). Additionally, we found that the effects of the task-relevant information ($1R$) become statistically insignificant when exclusively examined in cross-context comparisons (right panel).

**e.** Even when applying a lenient threshold ($p < 0.05$, uncorrected), we found no indication of significant effects related to task-irrelevant information ($1I$).

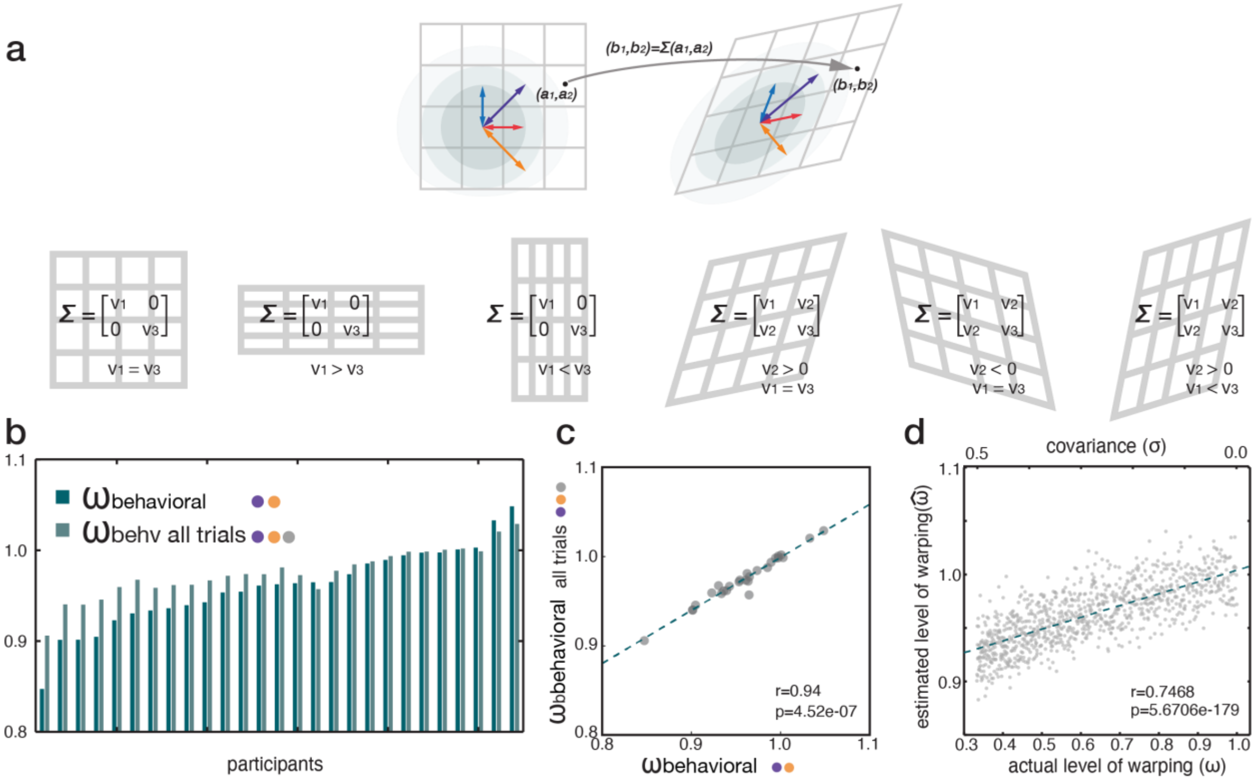


**Supplementary Figure 2. The procedure for approximating the level of warping in subjective representations from the reaction times (RT) of inferences**

**a.** The level of warping was estimated using a subject-specific covariance matrix ($\Sigma$). This matrix was obtained by examining the relationship between inferential angles (i.e., inference directions) and the extent to which Euclidean distances (E) in the true 4x4 structure accounted for RTs (**Eq.7-10**). Individual face positions within the warped space were then calculated by applying each participant's $\Sigma$ to their coordinates in the true 4x4 structure (**Eq.12**). The $\Sigma$ allows capturing not only the potential warping along the context invariant axis but other possible transformations, if there are any, in the 2-D representation.

**b.** $\omega_{\text{behavioral}}$ can be estimated from the $\Sigma$ from the distribution of $\varepsilon$ including the neutral trials ($\cos(\theta_t)$ equals to 0 or 1; grey dots) as well. The level of warping was estimated from RTs of each participant including all trials ($\omega_{\text{behavioral all trials}}$; lighter color) juxtaposed to the sorted $\omega_{\text{behavioral}}$ that were estimated only from RTs of congruent and incongruent trials (darker color).

**c.** The rank of $\omega_{\text{behavioral all trials}}$ was barely different from the rank of $\omega_{\text{behavioral}}$, suggesting results of the analysis using $\omega_{\text{behavioral}}$ won't be different when using $\omega_{\text{behavioral all trials}}$ instead.

**d.** Simulation results to validate the current methods. The level of warping estimated from RTs ($\omega_{\text{behavioral}}$) linearly correlates with the modulated actual level of warping ($\omega$) in the hidden subjective representations. To do this, we generated synthetic RTs by applying the mean regression coefficients and intercept estimated from participants' data to Euclidean distances of inferred trajectories of the same inference trials with the fMRI task on the representations that we modulated the levels of warping ($\omega$). We modulated $\omega$ while applying different levels of $\Sigma$ while varying $\sigma$ ($v_2$ and $v_3$ components in $\Sigma$) from 0.0 (no warping) to 0.5 (large warping). From the synthetically generated RTs, we estimated $\omega_{\text{behavioral}}$ using the same method.

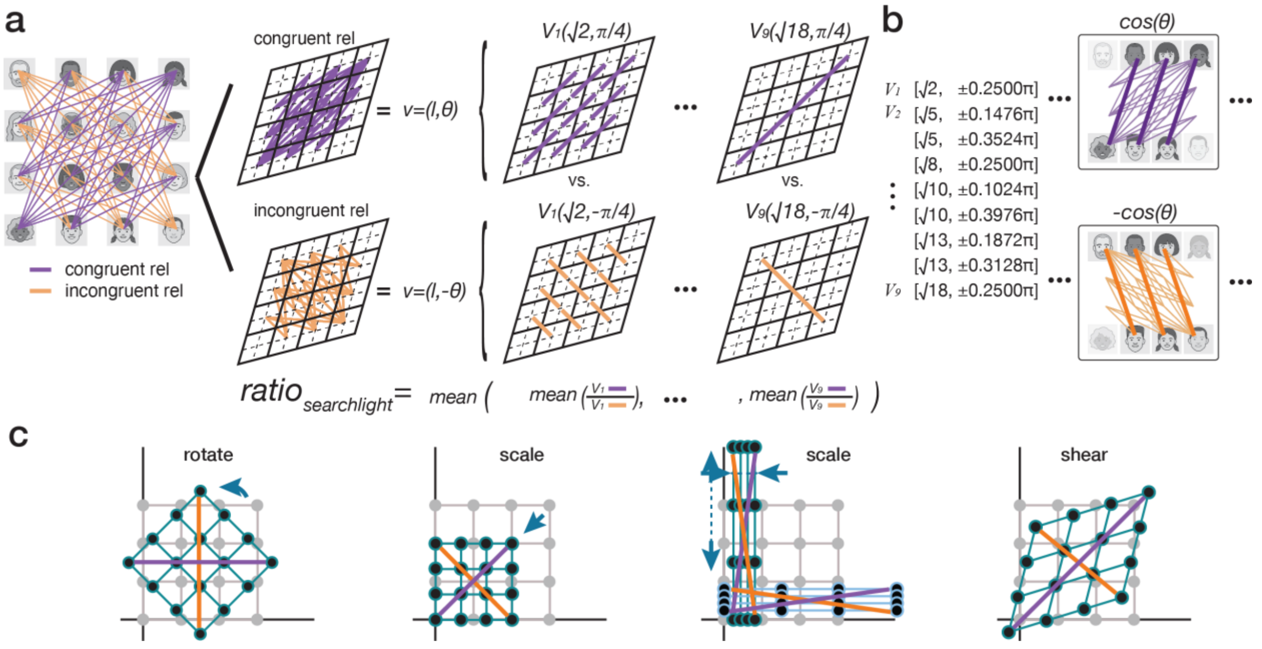


**Supplementary Figure 3. The procedure to measure the level of warping ($\omega_{\text{neural}}$) in the 2-D representation from activity patterns in the brain**

**a.** To estimate $\omega_{\text{neural}}$, we compared neural activity patterns of all possible pairs of congruent and incongruent relationships and computed the pairwise pattern dissimilarity using Euclidean distances. The sets of pattern dissimilarity estimated from congruent pairs (purple) were separated from those of incongruent pairs (orange). We computed the mean pattern dissimilarity for the pairs sharing the same vector in which both angles ($\theta$) and Euclidean distances ($l$) between two faces on the 2D social hierarchy are the same. Then, we computed the ratio of the mean pattern dissimilarity of the congruent pairs sharing the same vector (e.g. $v^C = [l, \theta^C]$) to that of the equidistant vectors of incongruent pairs that had symmetric vector angle (i.e. $v^I = [l, \theta^I]$ where $cos(\theta^C) = -cos(\theta^I)$). To test for warping, we computed the mean ratio between pattern dissimilarity of congruent pairs and that of their symmetric incongruent pairs in each searchlight ROI across the whole brain (**Eq.15**).

**b.** The nine vector sets ([Euclidean distance, vector angle]) are used for computing $\omega_{\text{Neural}}$ using representations of all 16 faces. In the additional control analysis, we tested for warping using representations of 12 faces while excluding those whose rank was the highest or lowest in both social hierarchy dimensions. The number of congruent/incongruent pairs sharing the same vector changes when excluding the representations. To elaborate further, while certain faces were used to calculate the mean activity patterns associated with specific angles in congruent vectors (e.g., $\theta$), the same faces were also used to compute the mean activity patterns associated with their counterpart angles in equidistant incongruent vectors ($\pi$-$\theta$ while $cos\,\theta = -\cos(\pi - \theta)$). In doing so, we were able to measure pattern dissimilarity not only from vectors with the same Euclidean distances but also from activity patterns of different pairs of identical faces. This careful control analysis allowed us to test whether warping in the representation was not driven by some representations having extreme ranks.

**c.** Measuring ratio between lengths of two diagonals allows testing of warping, defined by a shear transformation rather than other affine transformations (translation, rotation, or scaling in either dimension).

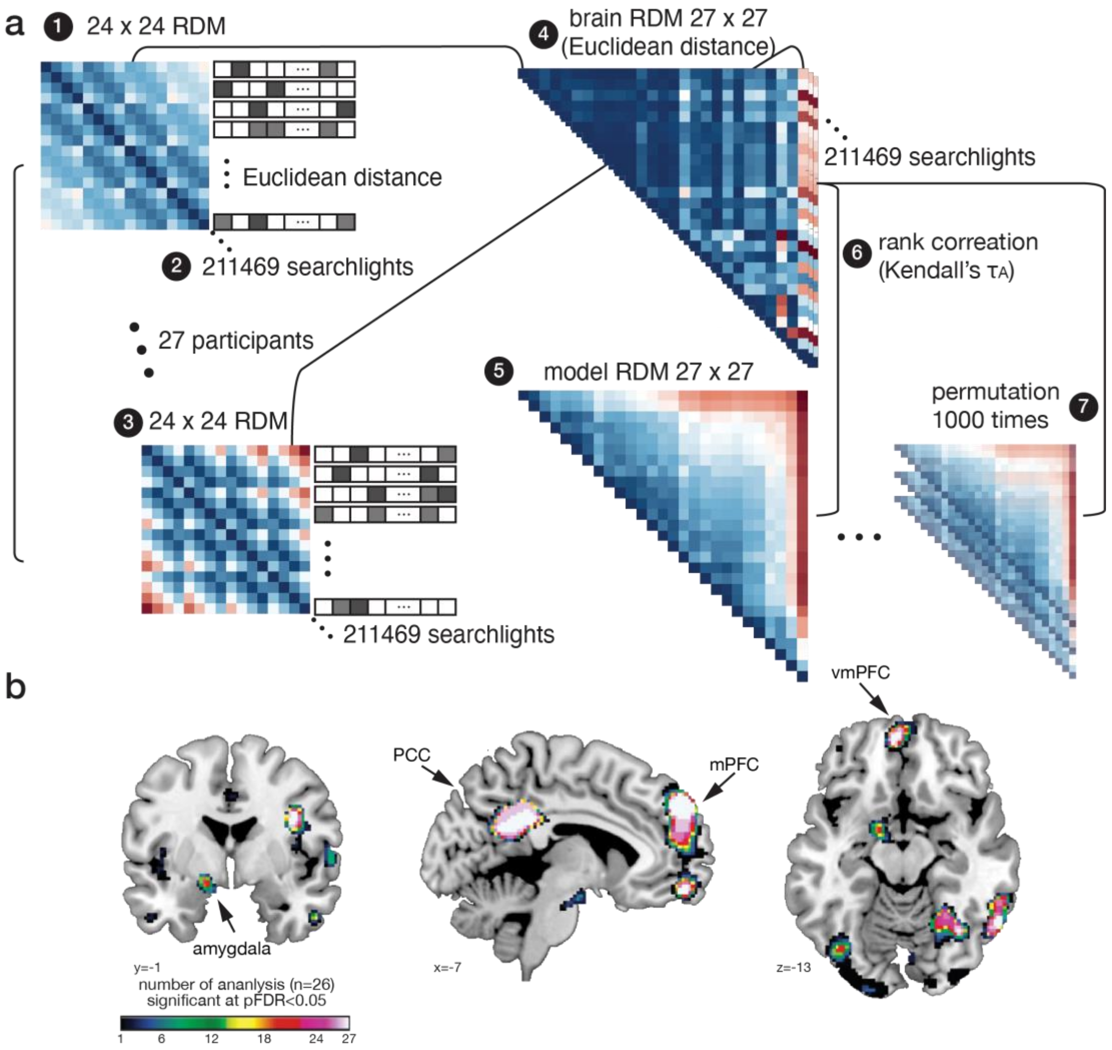


**Supplementary Figure 4. Intersubject representational similarity analysis (IS-RSA)**

**a.** The procedure of IS-RSA: (1) creating a representational dissimilarity matrix (RDM) by comparing the mean activity patterns associated with each face presented in each of the two task contexts using Euclidean distances; (2) An RDM was computed from each of 211,469 searchlight ROIs across the whole brain; (3) This procedure was repeated across participants (n=27); (4) The brain RDM for IS-RSA was created by comparing RDMs estimated from the same searchlight ROI across participants using Euclidean distances. This created an n×n sized RDM per ROI; (5) The model RDM for IS-RSA was computed from pairwise dissimilarity of the behaviorally estimated level of warping ($\omega_{\text{behavioral}}$; **Fig.4d**) across participants. (6) The rank correlation between the brain RDM of each searchlight and the model RDM was computed using Kendall's $\tau_A$ across the whole brain; (7) The rank correlation between the brain RDM and the model RDM was computed repeatedly while randomly shuffling the labels of $\omega_{\text{behavioral}}$ 1000 times to create the baseline distribution of Kendall's $\tau_A$. The t and p values were computed over the baseline using a normal distribution generated from the mean and standard deviation of the distribution of 1000 Kendall's $\tau_A$. This procedure was repeated for each searchlight ROI across the whole brain.

**b.** The results of leave-one-subject-out IS-RSA included data from 26 participants while excluding one participant at a time. We repeated this 27 times. The color map indicates the number of analyses showing significant intersubject similarity across 26 participants at the threshold $p_{FDR}<0.05$

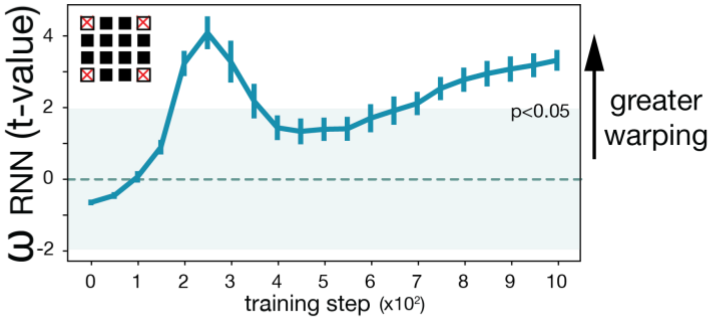


**Supplementary Figure 5. Time course changes of level of warping in RNN representations** while excluding the representations of faces who had the highest or lowest rank in either social hierarchy dimension. This result shows that the warping was not driven by the representations of certain faces but observed systematically across all face representations.

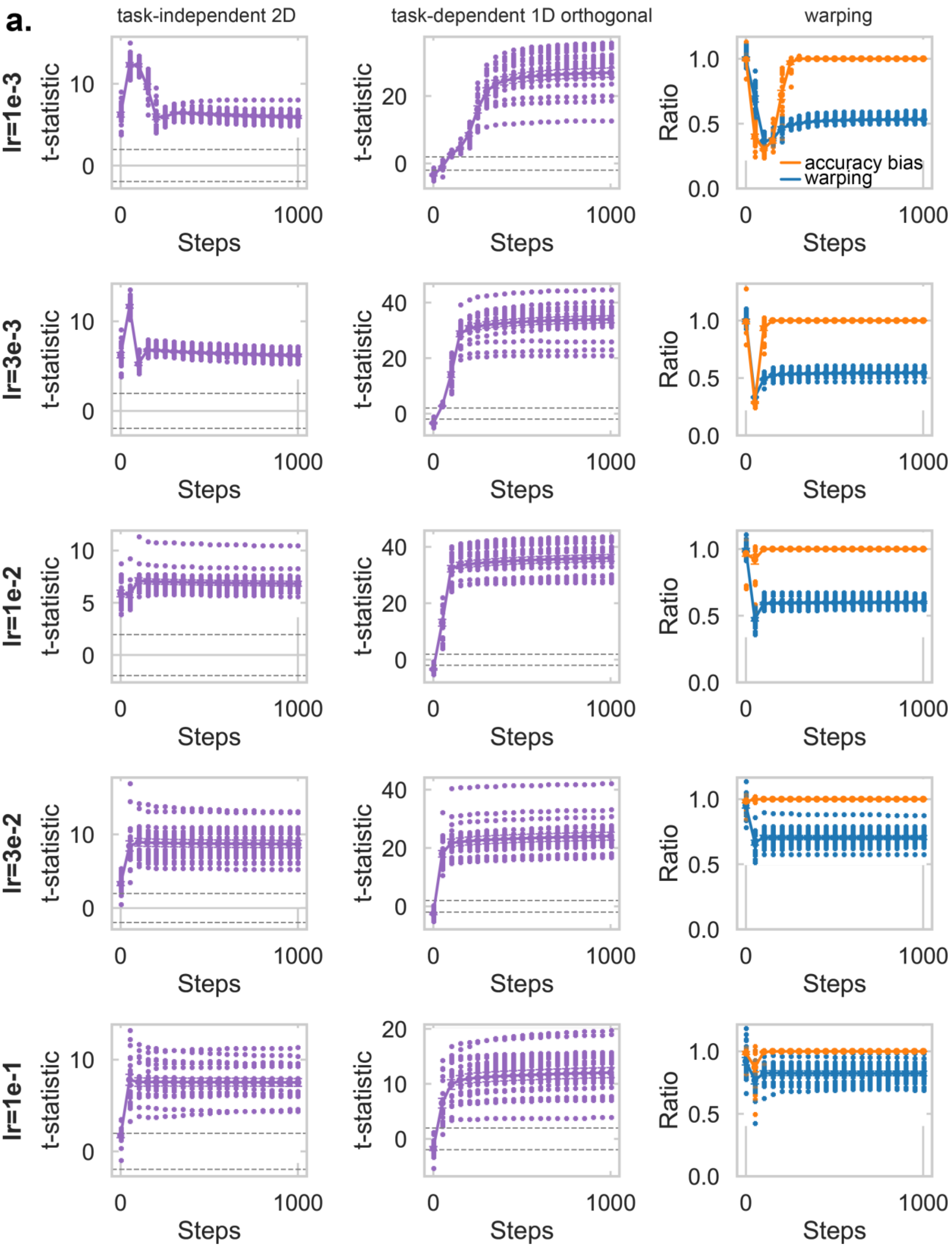
a.
task-independent 2D
task-dependent 1D orthogonal
warping
lr=1e-3
lr=3e-3
lr=1e-2
lr=3e-2
lr=1e-1
t-statistic
Ratio
Steps
accuracy bias
warping

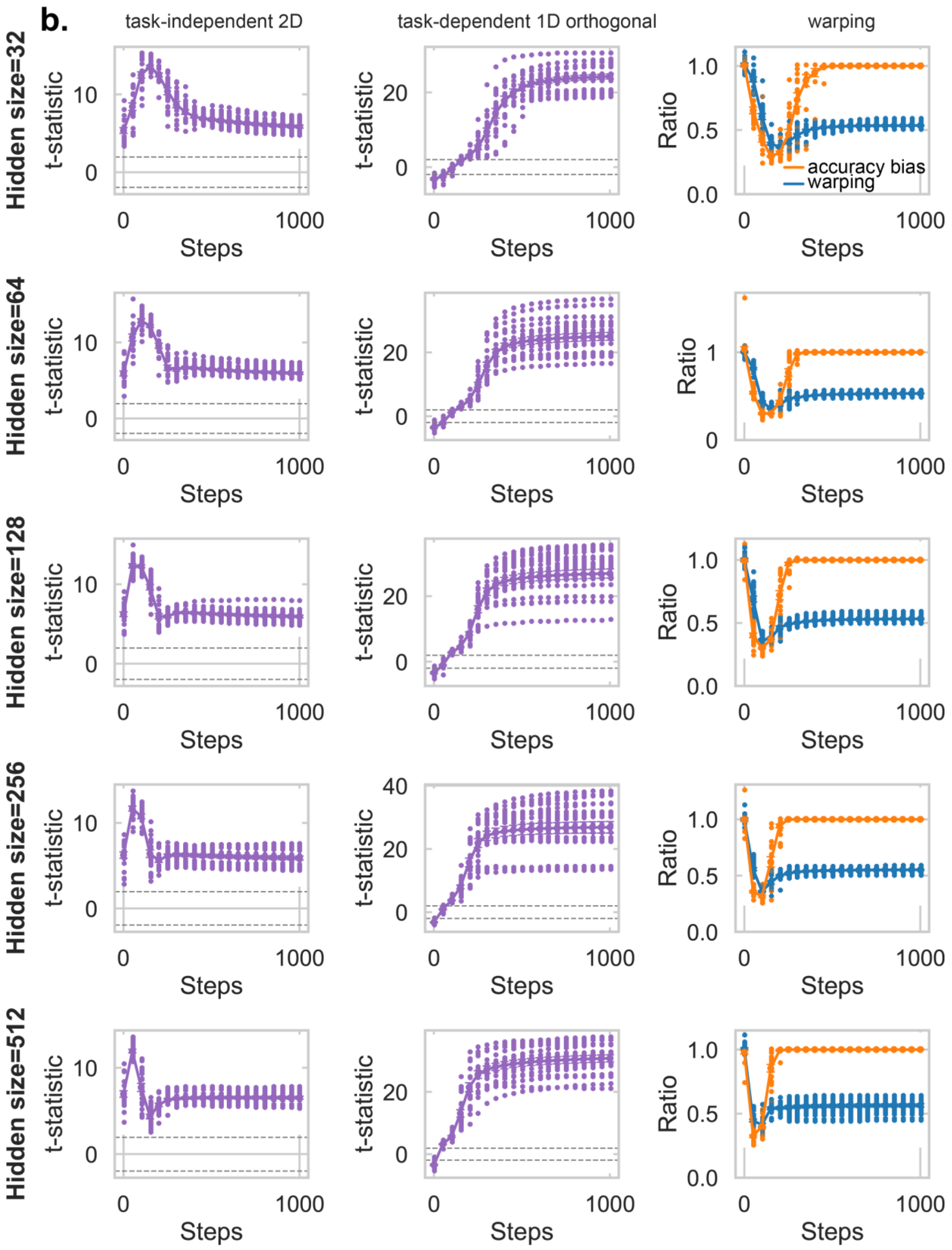
b.
task-independent 2D
task-dependent 1D orthogonal
warping
Hidden size=32
Hidden size=64
Hidden size=128
Hidden size=256
Hidden size=512
t-statistic
Ratio
Steps
accuracy bias
warping

c.

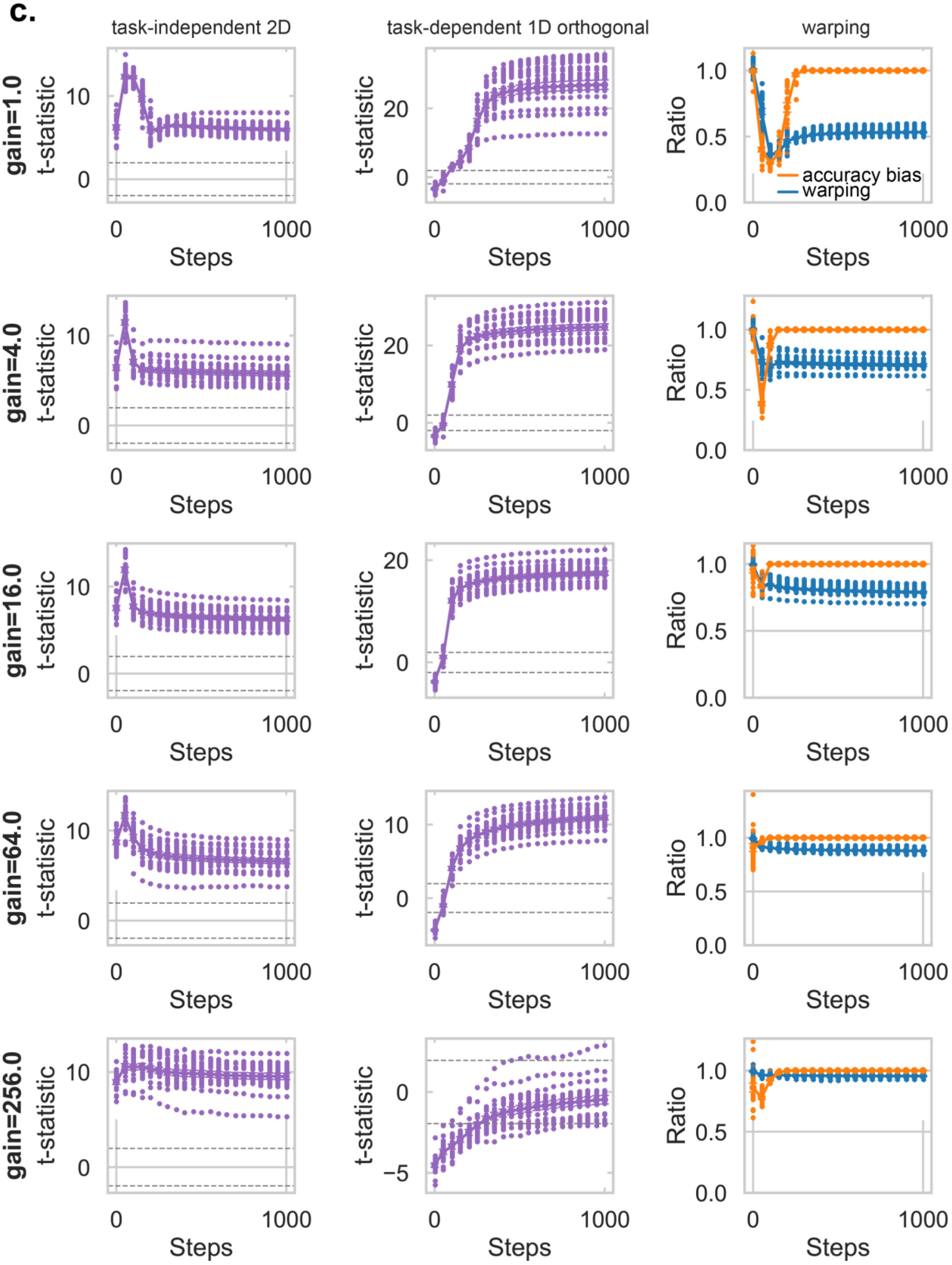
task-independent 2D
task-dependent 1D orthogonal
warping
gain=1.0
gain=4.0
gain=16.0
gain=64.0
gain=256.0
t-statistic
Ratio
Steps
accuracy bias
warping

**Supplementary Figure 6. Robustness test of RNN models.**

To assess the robustness of the RNN results, we conducted a series of control analyses in which we systematically varied key model parameters: the learning rate, hidden layer size, and the gain parameter controlling the variance of the Xavier-normal weight initialization[1]. Specifically, we tested **(a)** five learning rates (0.001, 0.003, 0.01, 0.03, 0.1), **(b)** five hidden sizes (32, 64, 128, 256, 512), and **(c)** five initialization gains (1.0, 4.0, 16.0, 64.0, 256.0); Increasing the gain (i.e., the variance of weight initialization) shifts the network toward a less rich and more lazy learning regime, in which feature learning is reduced and representations change minimally during training. For each manipulation, one parameter was varied while the others were fixed at their default values. For each model configuration, we performed the same RSA as described above to quantify the relative strength of the 2D (left panel) and orthogonal 1D (middle panel) representational geometries where the levels of 2D and 1D representations are shown as t-values. We also assessed the level of warping (right panel). Warping was indexed by the distance ratio between incongruent and congruent pairs (blue; lower values indicate greater warping) and by the accuracy bias between incongruent and congruent trials (orange). The results show that both 2D and 1D orthogonal representational formats consistently emerged across the full range of learning rates and hidden sizes. Warping effects were also stable across most configurations, though they decreased slightly at higher learning rates. When varying the initialization gain, the 2D representations remained robust, whereas the 1D orthogonal and warping effects weakened under higher gain conditions, corresponding to “less rich” or “more lazy” training regimes. Together, these findings indicate that the emergence of both 2D and orthogonal 1D representational structures is a robust outcome of learning in recurrent neural networks, rather than a consequence of specific parameter choices.

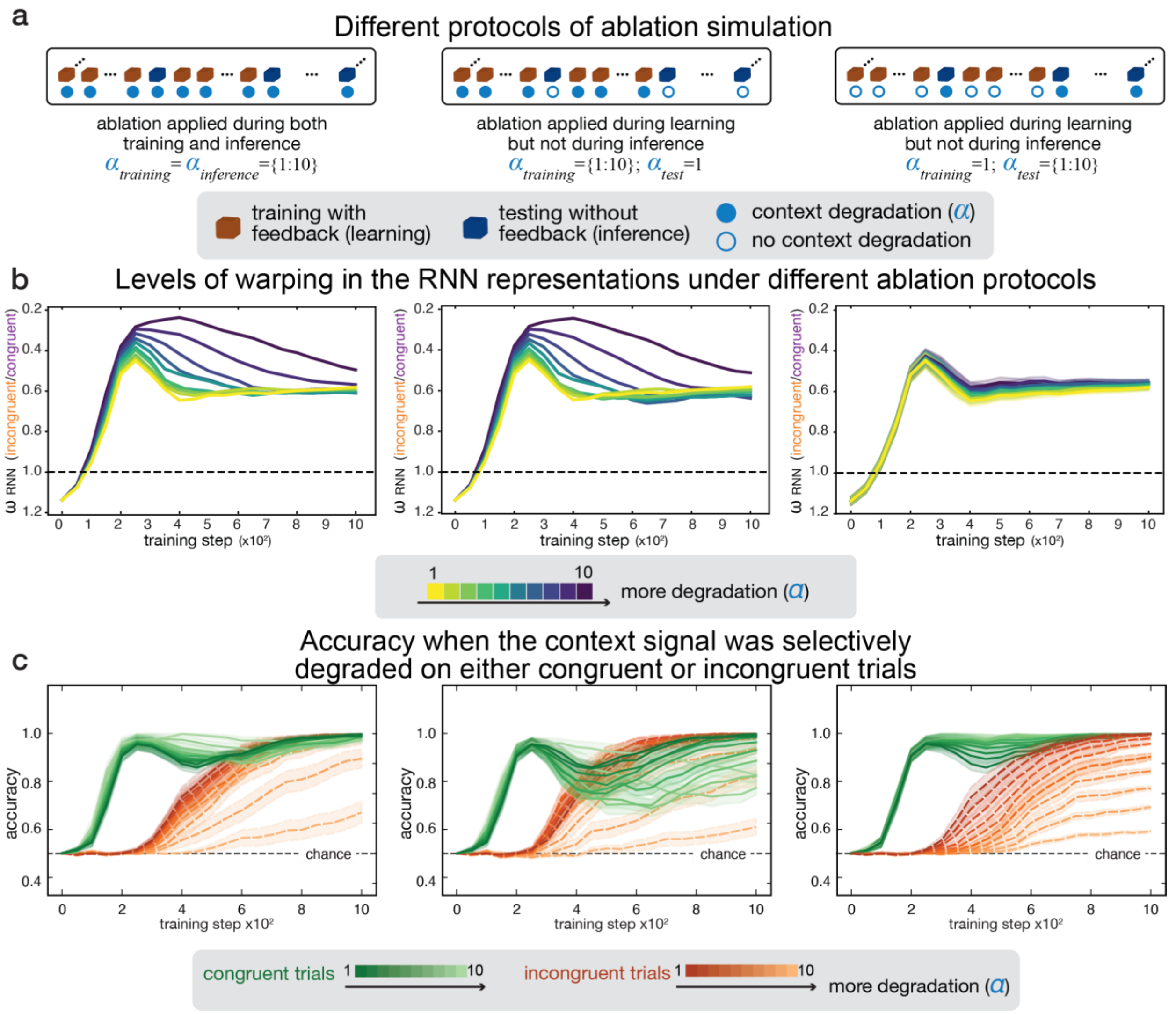


**Supplementary Figure 7. Effects of degrading context information during training and/or inference on representational warping and inference accuracy in RNN models**

**a.** Ablation protocols. Schematic illustration of the three ablation (context-degradation) protocols applied to the RNN. Left: Context ablation during both training and inference (α_training ∈ {1…10}, α_test ∈ {1…10}); Middle: Context ablation during training only (α_training ∈ {1…10}, α_test = 1); Right: Context ablation during inference only (α_training = 1, α_test ∈ {1…10}).

**b.** Warping of representational geometry in the hidden layer under different ablation protocols. Representational geometry was estimated during each test block, with one test block occurring after every 50 training blocks across a total of 1,000 training blocks. Each panel shows the evolution of representational warping ($\omega_{RNN}$ = distance incongruent / distance congruent) across training steps for one ablation protocol .Left: Warping when context was degraded during both training and inference; Middle: Warping when context was degraded only during training; Right: Warping when context was degraded only during inference. Warping increased substantially with greater context degradation ($\alpha$) only when the degradation was applied during training (left and middle panels), whereas degrading context information exclusively during inference had minimal effect on the representational geometry (right panel).

**c.** Effects of selective context degradation on inference accuracy. Inference accuracy was measured during test blocks while context degradation was applied during training and/or testing, selectively targeting either congruent (green) or incongruent (orange) trials. Plotted are the congruent-trial accuracies (green) and incongruent-trial accuracies (orange) across training steps for each ablation protocol. Left: Context degradation applied only on congruent trials during inference; Middle: Context degradation applied only on incongruent trials during inference; Right: Benchmark condition illustrating the stable performance when no degradation is applied during inference. Stronger degradation selectively impaired performance on incongruent trials, whereas accuracy on congruent trials remained largely unaffected. Importantly, degrading context during inference altered behavior but did not change representational warping (**Supplementary Fig.S7b**, right panel). This pattern arises because context information is needed to choose between competing responses even when the underlying representation is not warped, confirming that warping is induced during feedback-based learning and remains stable during inference.

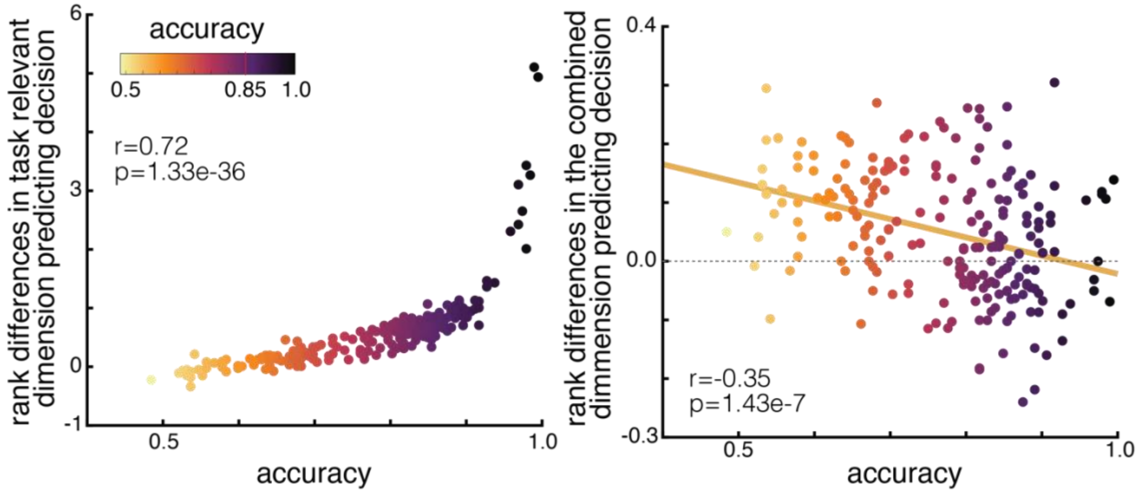


**Supplementary Figure 8. Relationships between individual differences in inference accuracy and their tendency of using context-dependent (left panel) and context-independent (right panel) decision values (DVs)**

When learning the relative ranks of faces through pairwise comparisons, participants could adopt different learning strategies. In a context-dependent learning strategy, participants learn rank relationships separately within each task dimension, such as competence versus popularity, thereby maintaining the independence of the two dimensions. During inference, they selectively retrieve rank information from the task-relevant dimension, consistent with a structured representation of task context. In contrast, a context-independent learning strategy does not preserve the distinction between dimensions. Instead, participants integrate experiences across tasks into a single combined dimension based on the overall win–loss ratio, reflecting how often one face was ranked higher or lower across all comparisons regardless of task context. This win–loss ratio serves as a context-independent decision value (DV), a scalar estimate derived from accumulated reinforcement history rather than context-specific relational structure. In the current task, accurate inference required participants to use rank differences in the task-relevant dimension, corresponding to the context-dependent DV, equivalent to distances between individuals in the task-relevant dimension in the high-dimensional representation shown in **Fig.2a** (right panel). Participants should not rely on rank differences in the combined dimension, corresponding to the context-independent DV, equivalent to distances between individuals in the low-dimensional representation shown in **Fig.2a** (left panel). Across participants (n=218), higher inference accuracy was associated with greater use of context-dependent DVs (left panel; $r=0.72$, $p=1.33\times10^{-36}$) and reduced use of context-independent DVs (right panel; $r=-0.35$, $p=1.43\times10^{-7}$). These relationships remained significant after partialling out individual RT congruency costs. RT congruency cost was estimated from a multiple linear regression predicting RTs on correct trials, quantifying the extent to which trial congruency explained RT while controlling for decision difficulty, as in **Fig.7d**. The relationships remained significant after this control analysis (context-dependent DV: partial $r=0.70$, $p=6.01\times10^{-33}$; context-independent DV: partial $r=-0.30$, $p=1.23\times10^{-6}$), indicating that these effects were not simply explained by generic congruency-related slowing. Participants needed to reach above 85% accuracy to continue participating in the fMRI experiment.

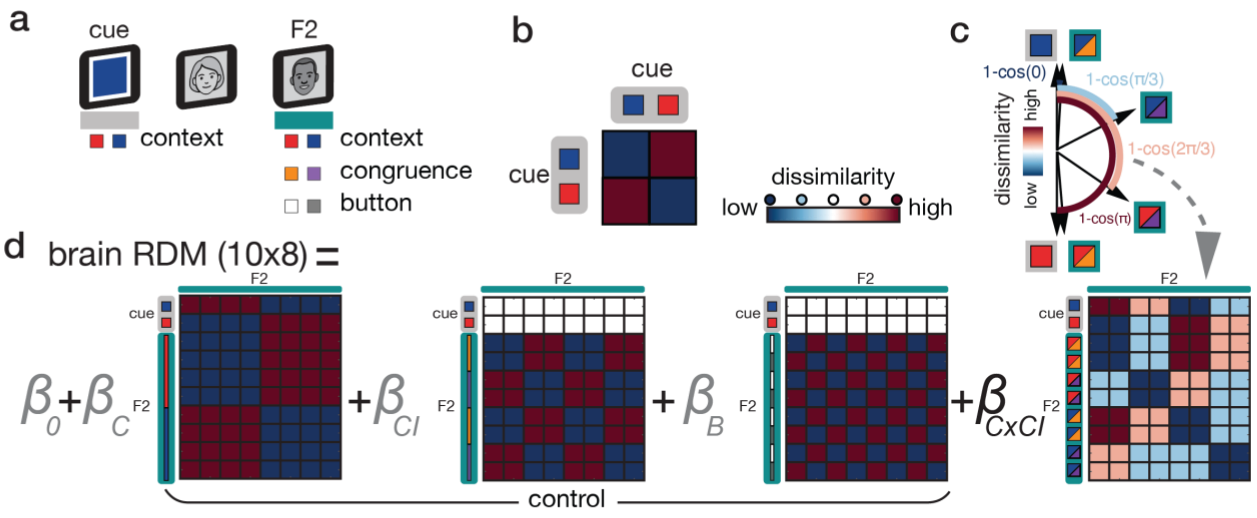


**Supplementary Figure 9. The multivariate representational similarity analysis (RSA) to test for greater task context reinstatement at the time of inferences on the incongruent compared to congruent trials**

**a.** For an additional RSA, we estimated the mean activity patterns of each task context (red: competence; blue: popularity) at the time of context presentation, and mean activity patterns of F2 grouped by task-relevant context (red: competence; blue: popularity), context congruence (orange: incongruent trial; purple: congruent trial), and the button press (white: right choice; grey: left choice) at the time of decision

**b.** The brain regions containing task context-specific representations were identified by comparing the mean activity patterns associated with the two task contexts at the time of context cue presentation.

**c.** A graphical model for creating a model representational dissimilarity matrix (RDM) to test the hypothesis that context reinstatement at the time of inferences is greater on the incongruent trials compared to congruent ones. Each arrow indicates the vector of representation of two task contexts (gray outline) and that of F2 grouped by context congruence in each of two task-relevant contexts (green outline). The model RDM reflects $1 - cos(angle)$ between two arrows. i.e. Dissimilarity between two task context-specific representations: $1 - cos(\pi)$; Dissimilarity between representations of the task-relevant context (red or blue) and incongruent decision under the corresponding task context (red+orange or blue+orange): $1 - cos(0)$; Dissimilarity between representations of the task-relevant context (red or blue) and congruent decision under the corresponding task context (red+purple or blue+purple): $1 - cos(\pi/3)$.

**d.** A brain RDM was created per searchlight ROI across the whole brain. The 8×10 sized RDM was created by comparing the activity patterns estimated at the two context cue presentations and those of eight types of inferences. To test the hypothesis while controlling for the other potential confounding representations, we used a general linear model (GLM) in which we included other regressors. The three regressors of non-interest included current task-relevant context (C), congruence/incongruence (CI), and button press (B). Using the GLM, we tested the hypothetical representation (the interaction effect between C and CI shown in **c**). This representation was tested within the independently defined ROI including the brain areas having a context-specific representation (as the result of **Fig.S9b**).

| Brain areas | Laterality | cluster size (k) | Peak activity (MNI) x | y | z | T | $P_{TFCE}$ |
|---|---|---|---|---|---|---|---|
| 2-D context-invariant representation | | | | | | | |
| Ventromedial prefrontal cortex / Medial orbitofrontal cortex (vmPFC/ mOFC) | Bilateral (B) | 844 | 14 | 42 | -22 | 5.83 | <0.01 |
| Hippocampus (HC) | Right (R) | 832 | 28 | -2 | -18 | 4.67 | 0.01 |
| Subgenual area | R | 832 | 12 | 8 | -20 | 4.15 | 0.02 |
| lateral OFC | Left (L) | 1844 | -24 | 26 | -16 | 4.55 | 0.01 |
| lateral OFC | R | 514 | 28 | 22 | -22 | 3.96 | 0.02 |
| Entorhinal cortex (EC) | R | 160 | 20 | 0 | -36 | 4.02 | 0.02 |
| 1-D orthogonal context-dependent representation | | | | | | | |
| Dorsomedial prefrontal cortex (dmPFC) | B | 16680 | -4 | 54 | 24 | 5.99 | 0.01 |
| Dorsomedial frontal cortex/ Pre-supplementary motor area (dmFC/pre-SMA) | B | | 4 | 14 | 48 | 5.59 | <0.01 |
| Lateral PFC | L | | -42 | 24 | 20 | 13.48 | <0.01 |
| Lateral PFC | R | | 42 | 26 | 18 | 9.03 | <0.01 |
| Precuneus/ posterior cingulate cortex (PCC) | B | 15321 | -4 | -64 | 28 | 13.6 | <0.01 |
| Inferior parietal lobule/ Temporoparietal junction (IPL/ TPJ) | R | | 46 | -64 | 22 | 5.2 | <0.01 |
| IPL/ TPJ/ Posterior superior temporal sulcus (STSp) | L | | -48 | -72 | 18 | 6.72 | <0.01 |
| Fusiform gyrus | L | | -40 | -60 | -20 | 6.44 | <0.01 |
| Fusiform gyrus | R | | 30 | -84 | -10 | 5.54 | <0.01 |

**Supplementary Table 1. Context-invariant 2-D and context-dependent orthogonal 1-D representations using whole-brain searchlight representational similarity analysis (RSA)**

The upper table, associated with **Fig. 3b**, shows brain areas where the dissimilarity between activity patterns for faces significantly correlates with their pairwise Euclidean distances computed from the idealized context-invariant representation (i.e. the true 2D social hierarchy).

The lower table, associated with **Fig. 3c**, shows brain areas where the dissimilarity between activity patterns for faces significantly correlates with their pairwise Euclidean distances computed from the idealized 1D orthogonal context-dependent representation. Group-level statistical significance was assessed using one-sided one-sample t-tests on the individual participants' searchlight RSA regression coefficients, with TFCE-based family-wise error correction applied for whole-brain multiple comparisons.

| | | 1D orthogonal representations | | | |
|---|---|---|---|---|---|
| | | All (pTFCE<0.05) | dmFC/ pre-SMA | lPFC | Precuneus/ PCC |
| 2D map-like representations | All (pTFCE<0.05) | 0.277 (0.171) | 0.238 (0.243) | 0.312 (0.121) | -0.247 (0.225) |
| | vmPFC/mOFC | -0.050 (0.810) | -0.202 (0.322) | 0.078 (0.704) | -0.182 (0.372) |
| | EC | 0.127 (0.536) | 0.095 (0.644) | 0.119 (0.564) | 0.143 (0.485) |
| | HC | 0.022 (0.915) | 0.283 (0.161) | -0.045 (0.827) | 0.228 (0.262) |

**Supplementary Table 2. The relationship between context-invariant 2-D and context-dependent orthogonal1-D representations in the brain across individuals**

Pearson's correlation coefficients (r) and two-sided P-values (in parentheses) are reported for the relationship between mean brain activity ($\beta$) of brain regions that represent 2D map-like and 1D orthogonal representations across participants. The $\beta$ values were estimated either from all brain areas that significantly represent 2D or 1D representations at a cluster-corrected threshold, $P_{TFCE}<0.05$, or from one of the selected clusters. The lack of a significant negative correlation (P>0.05) suggests that the brain represents both types of representations, rather than some individuals only showing one or the other representation.

**a. Multiple linear regression (log-transformed RTs)**

| Model 1: | Constant | Context (C) | Euclidean distance (E) | Task-relevant rank difference (D) | Task switching (S) |
|---|---|---|---|---|---|
| p: | 3.54E-05 | 2.18E-05 | 5.72E-08 | 1.40E-08 | 9.58E-01 |
| T: | 0.98** | -5.164** | -7.506** | -8.101** | 0.053 |
| B: | 1.17E-01 | -2.67E-02 | -4.82E-02 | -1.46E-01 | 4.21E-04 |
| se: | 2.39E-02 | 5.26E-03 | 6.54E-03 | 1.83E-02 | 8.13E-03 |

| Model 2: | Constant | Angle (A) | Euclidean distance (E) | Task-relevant rank difference (D) | Task switching (S) |
|---|---|---|---|---|---|
| p: | 8.37E-07 | 3.23E-04 | 5.63E-08 | 1.09E-08 | 9.65E-01 |
| T: | 6.422** | -4.142** | -7.513** | -8.209** | 0.044 |
| B: | 1.52E-01 | -5.30E-02 | -4.85E-02 | -1.47E-01 | 3.49E-04 |
| se: | 2.42E-02 | 1.30E-02 | 6.58E-03 | 1.82E-02 | 8.05E-03 |

| Model 3: | Constant | Interaction (C×I) | Euclidean distance (E) | Task-relevant rank difference (D) | Task switching (S) |
|---|---|---|---|---|---|
| p: | 3.24E-05 | 1.08E-05 | 5.28E-08 | 1.49E-08 | 8.84E-01 |
| T: | 5.014** | -5.431** | -7.539** | -8.072** | 0.148 |
| B: | 1.18E-01 | -2.02E-02 | -4.89E-02 | -1.45E-01 | 1.18E-03 |
| se: | 2.39E-02 | 3.79E-03 | 6.60E-03 | 1.83E-02 | 8.11E-03 |

| Model 4: | Constant | Task-irrelevant rank difference (I) | Euclidean distance (E) | Task-relevant rank difference (D) | Task switching (S) |
|---|---|---|---|---|---|
| | | | Congruent | | |
| p: | 5.34E-01 | 8.79E-07 | 3.41E-02 | 1.30E-01 | 8.16E-01 |
| T: | 0.63 | -6.403** | -2.237* | -1.566 | 0.235 |
| B: | 2.15E-02 | -1.18E-01 | -2.87E-02 | -3.78E-02 | 3.24E-03 |
| se: | 3.47E-02 | 1.87E-02 | 1.31E-02 | 2.46E-02 | 1.41E-02 |
| | | | Incongruent | | |
| p: | 7.82E-01 | 1.05E-03 | 2.71E-01 | 3.96E-03 | 5.50E-01 |
| T: | 0.279 | -3.688** | -1.126 | -3.162** | -0.606 |
| B: | -7.92E-03 | -4.74E-02 | -1.29E-02 | -7.67E-02 | -1.10E-02 |
| se: | 2.89E-02 | 1.31E-02 | 1.17E-02 | 2.47E-02 | 1.85E-02 |
| | | | Congruent - Incongruent | | |
| p: | 3.36E-01 | 1.94E-03 | 2.82E-01 | 2.29E-01 | 5.60E-01 |
| T: | 0.981 | -3.448** | -1.098 | 1.232 | 0.59 |
| B: | 2.94E-02 | -7.02E-02 | -1.58E-02 | 3.89E-02 | 1.42E-02 |
| se: | 3.05E-02 | 2.07E-02 | 1.47E-02 | 3.22E-02 | 2.46E-02 |

**b. Multiple logistic regression (Accuracy)**

| Model 1: | Constant | Context (C) | Euclidean distance (E) | Task-relevant rank difference (D) | Task switching (S) |
|---|---|---|---|---|---|
| p: | 9.35E-02 | 1.64E-02 | 1.02E-03 | 6.22E-02 | 9.70E-01 |
| T: | 0.741 | 2.566 | 3.7** | 1.949 | -0.038 |
| B: | 3.72E+00 | 3.34E-01 | 4.62E-01 | 7.50E+00 | -1.22E-01 |
| se: | 2.18E+00 | 1.33E-01 | 1.27E-01 | 3.92E+00 | 3.30E+00 |

| Model 2: | Constant | Angle (A) | Euclidean distance (E) | Task-relevant rank difference (D) | Task switching (S) |
|---|---|---|---|---|---|
| p: | 1.05E-01 | 4.91E-01 | 1.05E-03 | 6.26E-02 | 9.69E-01 |
| T: | 1.678 | -0.699 | 3.689** | 1.946 | -0.04 |
| B: | 3.93E+00 | -3.11E-01 | 4.67E-01 | 7.46E+00 | -1.28E-01 |
| se: | 2.39E+00 | 4.54E-01 | 1.29E-01 | 3.91E+00 | 3.28E+00 |

| Model 3: | Constant | Interaction (C×I) | Euclidean distance (E) | Task-relevant rank difference (D) | Task switching (S) |
|---|---|---|---|---|---|
| p: | 9.92E-02 | 1.54E-02 | 4.91E-04 | 6.12E-02 | 9.75E-01 |
| T: | 1.71 | 2.594 | 3.981** | 1.957 | -0.031 |
| B: | 3.66E+00 | 3.13E-01 | 4.99E-01 | 7.52E+00 | -1.02E-01 |
| se: | 2.18E+00 | 1.23E-01 | 1.28E-01 | 3.91E+00 | 3.33E+00 |

| Model 4: | Constant | Task-irrelevant rank difference (I) | Euclidean distance (E) | Task-relevant rank difference (D) | Task switching (S) |
|---|---|---|---|---|---|
| | | | Congruent | | |
| p: | 3.01E-03 | 3.21E-03 | 5.54E-01 | 4.26E-01 | 4.64E-01 |
| T: | 3.273** | 3.247** | -0.599 | 0.808 | 0.743 |
| B: | 5.26E+01 | 2.04E+01 | -4.86E+00 | 8.53E+00 | 8.36E+00 |
| se: | 1.64E+01 | 6.40E+00 | 8.27E+00 | 1.08E+01 | 1.15E+01 |
| | | | Incongruent | | |
| p: | 7.09E-05 | 9.17E-01 | 7.41E-01 | 2.05E-02 | 3.13E-01 |
| T: | 4.717** | 0.106 | 0.334 | -2.469* | -1.028 |
| B: | 9.64E+01 | 1.06E+00 | 2.99E+00 | -2.38E+01 | -9.32E+00 |
| se: | 2.08E+01 | 1.02E+01 | 9.11E+00 | 9.84E+00 | 9.24E+00 |
| | | | Congruent - Incongruent | | |
| p: | 9.95E-02 | 8.20E-02 | 5.34E-01 | 4.37E-02 | 2.48E-01 |
| T: | -1.708 | 1.809 | -0.63 | 2.121* | 1.181 |
| B: | -4.38E+01 | 1.93E+01 | -7.85E+00 | 3.24E+01 | 1.77E+01 |
| se: | 2.62E+01 | 1.09E+01 | 1.27E+01 | 1.56E+01 | 1.53E+01 |

**Supplementary Table 3. Four models utilizing multiple linear regression and multiple logistic regression to predict reaction times (RTs) and accuracy, respectively.**

**a.** Associated with **Fig. 4a**. Four models employing multiple linear regression were used to predict the log-transformed RTs of inferences (**Eq.2-5**). It is important to note that these four models were not designed to test different mechanisms underlying biases in RTs or different representational structures. Instead, their purpose was to assess whether the representation could be warped, with the null hypothesis assuming that congruence effects did not result in representational distortion. Each model employed distinct approaches to examine the relationships between warped representation and RTs, on a trial-by-trial basis, in a compatible manner that mitigated issues related to multicollinearity. Statistical significance of regression coefficients was assessed using two-sided t-tests for each model. No multiple-comparison correction was applied because each regression model tested a predefined hypothesis. **<0.01; *<0.05; B: mean beta value; se: standard error.

**b.** The study employed four models using multiple logistic regressions with the same regressors to predict accuracy. However, in this study, we gave priority to RTs as the primary dependent measure instead of accuracy. This decision was due to the limited variability in accuracy across trials among extensively trained participants. To thoroughly investigate the potential impact of warping in the representation, it was essential to ensure that participants had developed and possessed a representation that facilitated accurate inferences. While participants could perform the task without forming a representation and simply compare win/lose counts against others, we only included participants who achieved an accuracy of over 85% during the behavioral training phase to proceed with the fMRI experiment. This stringent selection approach resulted in consistently high and stable levels of accuracy, which did not exhibit substantial fluctuations at the trial level to capture fine-grained variations in cognitive control demands. Statistical significance of regression coefficients was assessed using two-sided t-tests for each model. No multiple-comparison correction was applied because each regression model tested a predefined hypothesis. **<0.01; *<0.05; B: mean beta value; se: standard error.

| Brain areas | Laterality | cluster size (k) | Peak activity (MNI) x | y | z | T | $P_{TFCE}$ |
|---|---|---|---|---|---|---|---|
| Group level | | | | | | | |
| HC | R | 105 | 28 | -12 | -22 | 4.5 | 0.05 |
| Supramarginal gyrus | R | 259 | 52 | -30 | 42 | 5.51 | 0.02 |
| Inferior temporal gyrus | R | 149 | 64 | -12 | -30 | 4.79 | 0.05 |
| Fusiform/ Parahippocampal gyrus | R | 238 | 44 | -38 | -14 | 4.6 | 0.04 |
| Dorsolateral prefrontal cortex | R | 104 | 42 | 34 | 24 | 4.56 | 0.05 |
| Individual differences | | | | | | | |
| Dorsomedial prefrontal cortex (dmPFC) | R | 357 | 22 | 48 | 16 | 9.9 | <0.01 |
| Amygdala | R | 4962 | 28 | 0 | -12 | 6.67 | 0.01 |
| Amygdala | L | | -20 | -8 | -16 | 3.3 | 0.03 |
| Lateral OFC | R | | 34 | 32 | -18 | 4.68 | 0.02 |
| Fusiform gyrus | L | | -36 | -32 | -18 | 4.24 | 0.02 |
| Ventromedial/ Medial prefrontal cortex (vmPFC/mPFC) | B | | -5 | 34 | 2 | 4.52 | 0.01 |
| Fusiform gyrus | R | 3027 | 28 | -54 | -6 | 5.88 | 0.01 |

**Supplementary Table 4. Warped map-like representations using whole-brain searchlight RSA**

The upper table, associated with **Fig. 5b**, shows brain areas showing significant warping in their map-like representations. The lower table, associated with **Fig. 5c**, shows brain areas with map-like representations whose level of warping correlates with individual differences in behaviorally estimated warping from reaction times (RTs) (ratio between diagonals on congruent and incongruent relationships between individuals, where their distances were approximated from the relationship between Euclidean distance and RTs). Group-level statistical significance was assessed using one-sided one-sample t-tests on individual participants' searchlight RSA maps, with TFCE-based family-wise error correction ($P < 0.05$) applied for multiple comparisons.

| Brain areas | Laterality | cluster size (k) | Peak activity (MNI) | | | |
| --- | --- | --- | --- | --- | --- | --- |
| | | | x | y | z | T |
| Posterior cingulate cortex (PCC) | B | 1728 | -8 | -48 | 28 | 7.28 |
| Fusiform gyrus | R | 1367 | 24 | -50 | -6 | 7.05 |
| Fusiform gyrus | L | 103 | -20 | -54 | -10 | 4.3 |
| Temporoparietal junction (TPJ) | L | 900 | -34 | -74 | 16 | 6.72 |
| TPJ | R | 606 | 40 | -64 | 18 | 6.48 |
| Medial/ Dorsomedial prefrontal cortex (mPFC/dmPFC) | B | 940 | -8 | 56 | 38 | 6.28 |
| Parahipocampal gyrus | R | 117 | 50 | -18 | -22 | 5.82 |
| Amygdala | L | 178 | -16 | -2 | -12 | 5.73 |
| Inferior frontal gyrus | R | 336 | 38 | 0 | 26 | 5.56 |
| Precentral gyrus | R | 453 | 36 | -18 | 40 | 5.34 |
| Dorsolateral frontal cortex (dlFC) | R | 172 | 38 | 24 | 40 | 5.01 |
| Medial orbitofrontal cortex (mOFC) | B | 248 | -4 | 56 | -14 | 4.87 |
| Inferior frontal gyrus/ Frontal inferior operculum | L | 370 | -46 | 14 | 16 | 4.73 |
| Inferior temporal gyrus | R | 237 | 54 | -46 | -26 | 4.39 |

**Supplementary Table 5. Intersubject RSA (IS-RSA)**

Associated with **Fig. 5d**, the brain areas where the similarity between representational geometry is associated with individual differences in behaviorally estimated warping from reaction times across participants. T-values compared to baseline are computed from 1000 permutations with random shuffling at the threshold $P_{FDR}<0.05$.

| Training step | T | P |
|---|---|---|
| 0 | 0.00E+00 | 1.00E+00 |
| 50 | 1.21E+00 | 2.25E-01 |
| 100 ✻ | 3.53E+00 | 4.10E-04 |
| 150 ✻ | 1.59E+01 | 4.12E-57 |
| 200 ✻ | 3.05E+01 | 1.94E-204 |
| 250 ✻ | 3.07E+01 | 2.32E-206 |
| 300 ✻ | 2.89E+01 | 4.19E-183 |
| 350 ✻ | 2.34E+01 | 7.18E-121 |
| 400 ✻ | 1.46E+01 | 3.05E-48 |
| 450 ✻ | 8.16E+00 | 3.30E-16 |
| 500 ✻ | 4.32E+00 | 1.56E-05 |
| 550 ✻ | 3.39E+00 | 6.97E-04 |
| 600 | -1.43E+00 | 1.54E-01 |
| 650 | -1.54E+00 | 1.25E-01 |
| 700 | 3.46E-01 | 7.29E-01 |
| 750 | 2.91E+00 | 3.57E-03 |
| 800 | 3.06E+00 | 2.18E-03 |
| 850 | 3.09E+00 | 1.97E-03 |
| 900 | -8.72E-01 | 3.83E-01 |
| 950 | 9.26E-01 | 3.54E-01 |
| 1000 | 1.93E-01 | 8.47E-01 |

**Supplementary Table 6. Timecourse of the effect of context congruency on RNN inference accuracy.**

Associated with **Fig. 6b**, logistic regression analyses were used to estimate the effect of context congruency on RNN inference accuracy for each independent RNN agent. Group-level statistical significance of the congruency effect was assessed using two-sided one-sample t-tests on the logistic regression coefficients across 20 independent RNN agents at each training step. ✻ indicates training steps where the congruency effect was significant ($P < 0.001$, uncorrected), illustrating the temporal evolution of the effect across training.

| Brain areas | Laterality | cluster size (k) | Peak activity (MNI) | | | | |
|---|---|---|---|---|---|---|---|
| | | | x | y | z | T | $P_{SVC}$ with TFCE |
| SMA | B | 8778 | -6 | 12 | 44 | 6.59 | <0.01 |
| PCC | B | | -4 | -28 | 28 | 6.35 | <0.01 |
| Anterior insula | L | | -30 | 18 | -8 | 5.48 | <0.01 |
| Lateral OFC | L | | -42 | 30 | 0 | 5.3 | 0.02 |
| Inferior temporal gyrus | L | | -48 | -6 | -32 | 3.7 | 0.02 |
| Inferior frontal gyrus | L | | -58 | 6 | 14 | 3.42 | 0.02 |
| HC | L | | -26 | -8 | -22 | 3.25 | 0.02 |
| Insula/ Lateral OFC | R | 5021 | 36 | 22 | -2 | 6.44 | <0.01 |
| Inferior frontal gyrus | R | | 58 | 4 | 18 | 5.73 | <0.01 |
| Lateral PFC | R | | 32 | 32 | 16 | 4.89 | <0.01 |
| vmPFC | B | 409 | -12 | 48 | -4 | 3.19 | 0.03 |

**Supplementary Table 7. Reinstatement of current task-relevant context during inferences on incongruent trials.**

Associated with **Fig. 8a**, we examined brain areas where the activity patterns elicited by the task-relevant context cue were reinstated to a greater extent during inferences on incongruent trials than congruent trials. This effect was tested within an independently defined region of interest (ROI) that combined all brain areas showing context-specific activity patterns at the time of context cue presentation, using small volume correction (SVC). Multiple comparisons were corrected within the ROI using TFCE.

| Brain areas | Laterality | cluster size (k) | Peak activity (MNI) | | | |
|---|---|---|---|---|---|---|
| | | | x | y | z | T |
| Dorsolateral Frontal Cortex (dlFC) | L | 1519 | -42 | 22 | 42 | 4.74 |
| Pre-supplementary motor area (preSMA) | B | | 0 | 24 | 50 | 3.22 |
| Dorsomedial Frontal Cortex (dmFC) | B | 237 | 26 | 24 | 50 | 3.78 |
| Dorsolateral Prefrontal Cortex (dlPFC) | B | 307 | 50 | 38 | 30 | 4.1 |

**Supplementary Table 8. The brain areas where the activity patterns represent the level of task-irrelevant information in the 1-D orthogonal representation having a correlation with the behaviorally estimated warping ($\omega_{Behavioral}$) across participants.**

Associated with **Fig. 8b**, we found the positive correlation between the behaviorally estimated level of warping in the 2-D representation ($\omega_{Behavioral}$) and the degree to which neural activity patterns were accounted for by task-irrelevant information in the 1-D context-dependent orthogonal representation. This finding implies that participants who exhibited less bias in their response times (RTs) during inferences were more effective in suppressing distances in the task-irrelevant dimension compared to the task-relevant dimension in brain regions displaying a 1-D task-dependent orthogonal representation. The threshold for determining a significant effect is established at $p<0.005$ uncorrected.
